# Early detection of pancreatic ductal adenocarcinoma by single-molecule profiling of pancreatic enzyme activities

**DOI:** 10.64898/2026.06.05.730533

**Authors:** Shingo Sakamoto, Hideto Hiraide, Junpei Hatakeyama, Takashi Kobayashi, Kazuki Takahashi, Takashi Inagaki, Shizuka Takagi-Niidome, Mayano Minoda, Tatsuya Kawaguchi, Toru Shibayama, Nozomi Iwakura, Masami Kawana, Tadahaya Mizuno, Mitsuyasu Kawaguchi, Hidehiko Nakagawa, Kyohhei Fujita, Yasuteru Urano, Atsuhiro Masuda, Juri Ikemoto, Yasutaka Ishii, Shiro Oka, Keiji Hanada, Yuzo Kodama, Toru Komatsu, Yu Kagami

## Abstract

Pancreatic ductal adenocarcinoma (PDAC) remains a leading cause of cancer-related mortality, largely due to diagnosis at advanced stages. Early detection through minimally invasive liquid biopsy holds promise for improving patient outcomes. Here, we report a blood-based liquid biopsy platform based on single-molecule enzyme activity profiling (SEAP), which detects proteoform-level alterations in circulating pancreatic enzymes at single-molecule resolution. Using a tissue-centric biomarker discovery strategy, we identified activity signatures of pancreas-specific digestive enzymes associated with PDAC. A combinatorial classifier detected PDAC (stage I–IV) with 95.5% specificity and 75.0% sensitivity (74.2% for stage I–II) across 690 blood samples collected from multiple hospitals and biobanks. Performance was further validated in an independent cohort enriched for early-stage disease, where 54.2% (13/24) of stage IA, 64.3% (9/14) of stage IB, and 21.4% (3/14) of stage 0 lesions were classified as positive. These findings support the clinical potential of SEAP for early detection of PDAC.

## Introduction

Pancreatic ductal adenocarcinoma (PDAC) remains one of the most lethal malignancies, with a 5-year survival rate of 12%^1,2^. Prognosis remains poor largely because most patients are diagnosed at advanced stages (stages III-IV). Detecting PDAC at an earlier stage (0–II), when resection is possible, could significantly improve survival outcomes^3,4^.

Liquid biopsy has emerged as a minimally invasive strategy for detecting cancers through analysis of blood-based biomarkers^5^. While this approach holds particular promise for the early detection of PDAC, most efforts have focused on circulating tumor DNA (ctDNA), exosomes, or protein markers, yet clinical performance remains insufficient for early-stage detection^3^. Several critical challenges remain: (1) many studies rely on single-center cohorts, limiting their generalizability across populations; (2) existing markers often lack cancer-type specificity, making it difficult to distinguish PDAC from other malignancies; and (3) most platforms show limited sensitivity for detecting early-stage disease.

In the present study, we developed a liquid biopsy platform based on single-molecule enzyme activity profiling (SEAP) of a panel of pancreatic digestive enzymes to address these limitations. By integrating fluorogenic substrate probes with a microfabricated chamber array, this method achieved high-sensitivity analysis of enzyme activity at single-molecule resolution (**Figure 1a–c**)^6–8^. A key advantage of this platform is its ability to resolve distinct proteoforms arising from altered protein–protein interactions or post-translational modifications, which are often masked in conventional bulk assays^8^. Using this SEAP platform, we previously identified a specific reduction of DPP4-FAPα heterodimers in the plasma of PDAC patients^9^. Although this marker shows responsiveness even in stage I-II disease, its overall patient coverage remains limited. In this study, we sought to extend our approach by integrating a multi-proteoform panel designed to capture the broader landscape of the disease. Central to this evolution was a “tissue-centric” discovery strategy, which prioritizes enzymes with highly restricted expression to isolate organ-specific signals from systemic biological noise in circulation.

**Figure 1.**
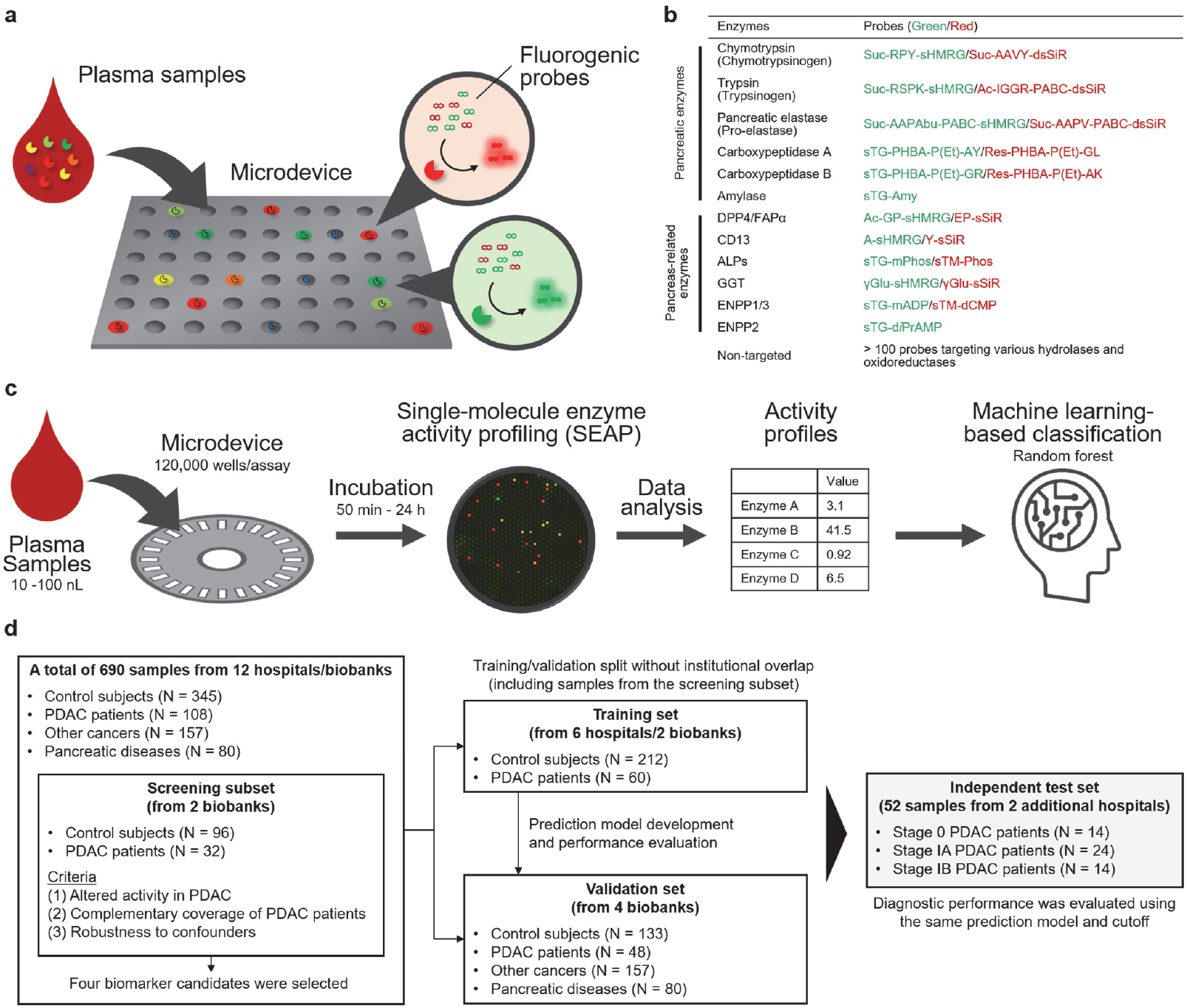
Single-molecule enzyme activity profiling (SEAP)-based liquid biopsy platform for the detection of PDAC. (a) Concept of SEAP. Enzyme activities in plasma are measured using multicolored (green and red) fluorogenic probes in a microdevice containing multiple reactors at the single-molecule level. (b) Fluorogenic probes used to measure the activities of the targeted enzymes. Chemical structures of the probes are provided in **Supplementary Figures**, and the optimized assay conditions are provided in **Table S4.** (c) SEAP-based classification of PDAC. Diluted plasma is mixed with fluorogenic probes, loaded into the microdevice, and incubated for enzymatic probe cleavage. Fluorescence signals in individual microreactors generate enzyme activity profiles that are integrated for PDAC classification. (d) Study design for biomarker discovery and validation. Plasma samples from multiple hospitals and biobanks were analyzed to develop a prediction model. Samples were divided into training and validation sets without institutional overlap, and performance was evaluated in the validation set and an independent test set enriched for stage 0–I PDAC cases.

Blood contains proteins derived from diverse tissues. When targeting ubiquitously expressed proteins, their systemic fluctuations do not necessarily reflect the pathological state of a specific organ. Furthermore, such proteins are often confounded by leakage from blood cells, which is a major source of inter-institutional variation. In contrast, proteins with highly restricted tissue expression are inherently less susceptible to such systemic noise. The pancreas is suitable for this “tissue-centric” biomarker discovery approach using the SEAP platform because it uniquely synthesizes a diverse array of digestive enzymes, allowing for high tissue specificity. While pancreatic enzymes such as elastase, amylase, and lipases have already been measured clinically using ELISA or conventional biochemical assays, the proteoform resolution of the SEAP platform allows us to capture their activity landscapes arising from heterogeneous enzyme states. This proteoform-level analysis enables the extraction of information with sufficient depth to achieve accurate detection of PDAC.

### Study design and screening strategy

The SEAP assay was performed by measuring the activity of single enzyme molecules confined in small reactors of a microdevice through turnover-based metabolism of fluorogenic substrates. By loading diluted biological samples into a microdevice bearing multiple reactors, theoretically 0 or 1 enzyme molecule is present in each reactor, and the unique reactivity profile of each enzyme molecule is measured as a fluorescence readout (**Figure 1a**)^8,10^. Recent advances in assay development have expanded the repertoire of targetable enzymes, now spanning diverse functional classes, such as proteases/peptidases^7,9,11^, glycosidases^12^, phosphatases^6,13^, esterases^14^, and oxidoreductases^15^. Under the concept of tissue-centric biomarker discovery approach, we curated a list of proteins with pancreas-specific expression profiles (**Table S2**) and developed a panel of single-molecule assays that cover major pancreas-specific enzymes, including chymotrypsin (CTRB1, CTRB2), trypsin (PRSS1, PRSS2), pancreatic elastase (CELA3A), carboxypeptidase A (CPA), carboxypeptidase B (CPB), and amylase (**Figure 1b**). In addition to these tissue-specific targets, we screened more than 150 SEAP assays targeting a variety of enzymes to identify systemic or microenvironmental alterations associated with pancreatic pathologies^16,17^ (**Table S3**).

We constructed an assay workflow using a mass-producible cyclic olefin copolymer (COP) microdevice to perform the SEAP assay with high throughput, reliability, and repeatability (**Figure 1c**)^9^. Using this platform, biomarker screening was performed in three steps. (1) Characterization of candidates by comparing 32 samples from patients with PDAC and 96 non-cancer control subjects to identify those showing altered activity patterns; (2) selection of biomarkers that can complement patient coverage; and (3) confirmation of minimal variation owing to pre-analytical confounders, such as hemolysis and sample handling conditions.

Markers that met these three criteria were selected, and their performance was validated using 690 blood samples (345 control subjects, 108 patients with PDAC, 80 patients with other pancreatic diseases, and 157 patients with other cancers) collected from 12 hospitals and biobanks in Japan and the United States (**Figure 1d**). This international, multi-institutional validation effectively filtered out candidate biomarkers susceptible to cohort-specific variations or site-specific biases. After the selection, we identified three enzyme classes, DPP4/FAPα, chymotrypsin (CTRB2 and CTRB1), and CELA3A, which provide complementary patient coverage (**Figure 2a, 2f, S2, S3**) and stability against various potential confounders and inter-institutional differences. Below, we summarize the screening workflow and the results of the analysis of 690 blood samples for identified biomarkers.

**Figure 2.**
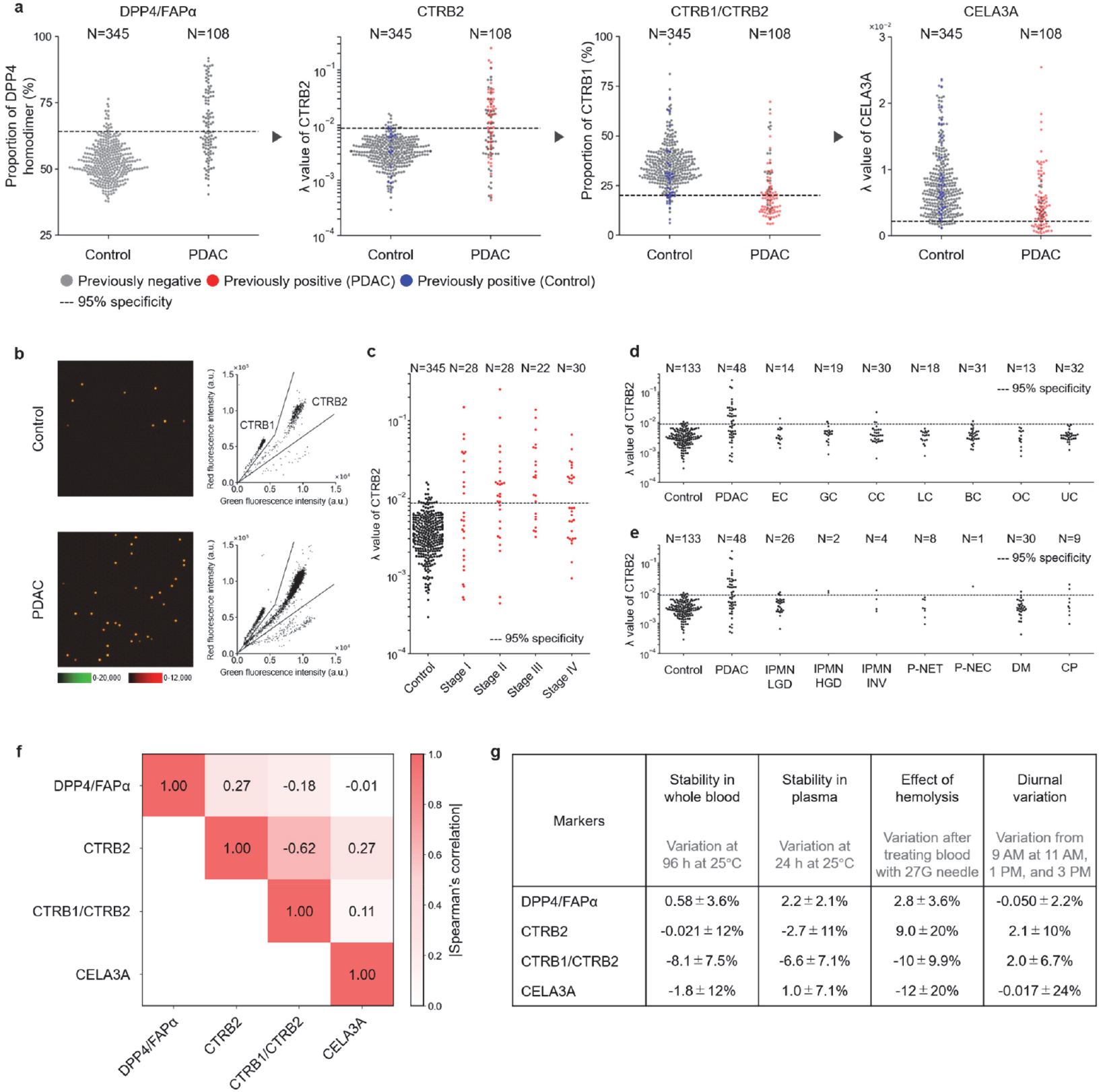
Screening workflow of PDAC biomarker enzymes. (a) Stepwise screening workflow prioritizing orthogonal patient coverage. Dot plots show biomarker distributions in patients with PDAC and control subjects from the combined training and validation sets (PDAC, N = 108; Control, N = 345). Dashed lines indicate cutoffs corresponding to 95% specificity (i.e., 95% of control samples are classified as negative). From the second panel onward, red and blue dots represent samples already classified as positive in the preceding steps (red, PDAC; blue, control), whereas gray dots indicate samples carried forward to the current step. Accordingly, gray dots in the PDAC group represent samples that were false negatives in the preceding step. (b-e) Measurements of chymotrypsin activity as a representative biomarker identified in this study. (b) Representative fluorescence images of the microdevice and corresponding scatter plots obtained after loading Suc-RPY-sHMRG (30 μM), Suc-AAVY-dsSiR (30 μM), IR-dye 800 (30 μM), and plasma samples (1:2,500 dilution) from control subjects or PDAC patients into HEPES-Na buffer (100 mM, pH 7.4) containing MgCl_2_ (1 mM), CaCl_2_ (5 mM), NaCl (1 M), DTT (100 μM), trypsin (25 μg/mL), and Triton X-100 (150 μM), followed by incubation at 25°C for 20 h. (c) Distribution of lambda (signal numbers per all analyzed chambers) value of CTRB2 in control subjects and patients with PDAC, stratified by disease stage. (d) Distribution of lambda value of CTRB2 in control subjects and patients with cancers from various tissues. (e) Distribution of lambda value of CTRB2 in control subjects and patients with PDAC or other diseases related to pancreatic dysfunction. (f) Pairwise correlation matrix of Spearman’s rank correlation coefficients constructed using data from the combined training and validation sets (PDAC, N = 108; Control, N = 345). Correlation coefficients were calculated using PDAC samples only. (g) Stability assessment of the measurements. Changes after storage were calculated from the mean of technical replicates (n = 3) for each sample and summarized as mean ± s.d. across biological replicates (N = 5). Changes after hemolysis treatment are expressed as mean ± s.d. across biological replicates (N = 3), with a single measurement per sample. Diurnal variation was calculated as the mean relative change from the 9:00 time point to 11:00, 13:00, and 15:00 for each individual and summarized as mean ± s.d. across biological replicates (N = 8). Numbers above the plots indicate sample size. **Abbreviations:** EC, esophageal cancer; GC, gastric cancer; CC, colorectal cancer; LC, lung cancer; BC, breast cancer; OC, ovarian cancer; UC, uterine cancer; IPMN-LGD, intraductal papillary mucinous neoplasm with low-grade dysplasia; IPMN-HGD, IPMN with high-grade dysplasia; IPMN-INV, IPMN with an associated invasive carcinoma; P-NET, pancreatic neuroendocrine tumor; P-NEC, pancreatic neuroendocrine carcinoma; DM, diabetes mellitus; CP, chronic pancreatitis; s.d., standard deviation.

### Identification and validation of PDAC biomarker signatures of pancreas-specific enzymes

First, we validated the performance of the DPP4-FAPα complex, a previously reported biomarker candidate^7,9^, using 690 blood samples. Specifically, 54 of 108 patients with PDAC exhibited an elevated value in the ratio of DPP4– DPP4/(DPP4–FAPα + DPP4–DPP4), exceeding the 95% specificity cutoff defined using control subjects (N = 345, **Figure 2a**). The DPP4/FAPα signature also retained comparable coverage in patients with stage I–II PDAC (30 of 56 stage I-II cases, **Figure S4f**). We also confirmed that the elevation was rarely observed in other cancers (e.g., gastric, colorectal, and lung cancers, N = 157, **Figure S4g**) or other pancreatic diseases (e.g., diabetes and chronic pancreatitis, N = 80, **Figure S4h**), indicating high specificity for PDAC.

Building on this validated framework, we performed activity-based screening to identify additional biomarker candidates. Consistent with our design, the identified hits were predominantly pancreas-specific enzymes, rather than ubiquitously expressed targets. The first hit was chymotrypsin B2 (CTRB2) activity after proteolytic activation with trypsin. Chymotrypsin is a digestive enzyme expressed exclusively in the pancreas in the form of a zymogen (chymotrypsinogen) and is secreted into the gastrointestinal tract^18,19^. Despite its pancreas-restricted expression, chymotrypsin has received limited attention as a circulating biomarker because of its established role as an exocrine digestive enzyme. Chymotrypsin activity could be detected in blood samples using the SEAP platform, especially when the blood samples were treated with trypsin to proteolytically activate chymotrypsinogen (**Figure S5b**). Under trypsin-activated conditions, the optimized fluorogenic probes revealed the presence of multiple active species, assigned to chymotrypsin B1 (CTRB1) and B2 (CTRB2), based on comparison with recombinant enzymes (**Figure 2b, Figure S5c**). Patients with PDAC demonstrated a consistent increase in CTRB2 levels (57 of 108 patients; **Figure 2a, S5e**). Importantly, CTRB2 retained comparable coverage in patients with stage I–II PDAC (29 of 56 patients; **Figure 2c**). The increase was uniquely observed in PDAC compared to other cancers (**Figure 2d**), confirming the validity of targeting tissue-specific enzymes. The marker did not respond to major pancreatic diseases and showed high specificity for PDAC (**Figure 2e**). Critically, patients with altered CTRB2 activity were orthogonal to those responding to DPP4/FAPα (**Figure 2f, S2**) and the combination of the two biomarkers significantly increased diagnostic performance (**Figure 2a**).

The second biomarker also involved chymotrypsin activity. A detailed analysis of the activity patterns observed in the chymotrypsinogen analysis indicated that CTRB1 and CTRB2 activities, although generally correlated, exhibited a notable discrepancy in a subset of patients with PDAC. We observed that the attenuation of CTRB1 relative to that of CTRB2 was characteristic of the disease. To capture this signature, we introduced a ratiometric parameter, CTRB1/(CTRB1+CTRB2) (**Figure 2a, S6a, S6b**), which further enhanced the sensitivity of our panel by identifying patients who would otherwise have been missed based on absolute activity levels alone. Alteration of this new parameter, named the CTRB1/CTRB2, was uniquely observed in PDAC compared to other cancers (**Figure S6d**), and was not observed in other pancreatic diseases (**Figure S6e**).

We studied the activity of trypsin (derived from trypsinogen), another pancreatic enzyme. When the blood samples were treated with enterokinase and chymotrypsin, multiple activity clusters composed mainly of two trypsin subtypes, PRSS1 and PRSS2, were observed (**Figure S8**). While enterokinase was considered sufficient for trypsinogen activation, the addition of chymotrypsin was necessary for reliable activity detection, probably because it neutralizes excess serpins (serine protease inhibitors) in the blood to keep the activated trypsin intact. Patients with PDAC showed increased activity of PRSS1 and PRSS2 (**Figure S9b, S9c**), but patients with increased PRSS1 and PRSS2 activity overlapped with those with increased CTRB2 activity (**Figure S3, S9e**). This implied that the increase in pancreatic proteases shares the same underlying mechanism as their release into the bloodstream. Because CTRB2 identified more PDAC cases than PRSS1 and PRSS2 at 95% specificity, we retained CTRB2 as the final candidate (**Figure S9f**).

Similarly, other pancreas-specific enzymes, such as CPAs, CPBs, and amylase, possessed unique activity clusters that were elevated in patients with PDAC. However, the positive cases for these markers overlapped with those responding to CTRB2, indicating that their inclusion did not significantly improve diagnostic accuracy. Extended screening for non-tissue-specific enzymes identified several candidates, including specific activity clusters of CD13, TNAP, GGT, ENPP1, and ENPP3, which showed unique activity alterations in patients with PDAC. Nevertheless, these markers were highly correlated with the DPP4/FAPα signature and were therefore excluded from the final panel. A correlation analysis of all of these candidate markers clearly delineated two distinct clusters: one overlapping with the DPP4/FAPα signature (from non-specific enzymes) and another comprising the pancreas-specific enzymes (**Figure S3**).

After the continued search for orthogonal biomarkers from these two classes, we identified the altered activity of pancreatic elastase (CELA3A) as an additional biomarker (**Figure 2a, 2f, S2, S3**). CELA3A activity was not detectable with the conventional amide-modifying fluorogenic probe design of sHMRG^9^, likely due to steric hindrance limiting substrate access to the catalytic pocket, whereas fluorogenic probes with an extended linker structure showed sufficient reactivity toward CELA3A for use in SEAP assays (**Figure S19a**). Subtypes of elastases derived from the pancreas and neutrophils were distinguished using an optimized set of probes (**Figure S19b**). We observed a decrease in active CELA3A in a subset of patients with PDAC (26 of 108 samples) (**Figure 2a, S20b, S20c, S20d**). The decrease was also observed in patients with stage I–II PDAC (15 of 56 stage I-II cases, **Figure S20e**). In this case, the blood sample was not treated with trypsin or other proteases to proteolytically activate the zymogen, so the assay detected only the “active” CELA3A population. The number of active elastases did not directly correlate with the total elastase levels measured by conventional ELISA^20,21^ or SEAP under trypsin-treated conditions to proteolytically activate pro-elastase^22,23^ (**Figure S21d, S21e**). These results suggest that the active fraction, rather than the total protein, of CELA3A provides superior diagnostic utility for PDAC. Circulating elastase activity is affected by protease inhibitors, such as serpins and macroglobulins^24,25^. Macroglobulin-associated elastase remains active toward small molecule probes while being shielded from protein substrates and serpins^26^. Consistent with this, we observed that active elastase in circulation was not blocked by the addition of serpins, raising the possibility that the observed active species were conjugated to macroglobulin rather than the intact protein (**Figure S28**). Therefore, the observed decline in active CELA3A likely reflects alterations in the balance between protease activity and its inhibition in circulation, potentially driven by changes in protease inhibitor dynamics within the PDAC microenvironment^27,28^.

### Evaluation of the stability of the assay readouts

Having identified four complementary biomarker signatures (DPP4/FAPα, CTRB2, CTRB1/CTRB2, and CELA3A) that collectively cover a broad range of patients with PDAC, we next evaluated their technical robustness and susceptibility to potential confounding factors. The technical reproducibility of the assay showed generally low coefficients of variation, with mean CVs of 1.0% for DPP4/FAPα, 8.0% for CTRB2, 5.9% for CTRB1/CTRB2, and 16% for CELA3A. In addition, their activity readouts were stable over various pre-analytical variables, such as storage conditions, plasma preparation methods, and timing of blood collection (**Figure 2g**). We confirmed that all markers remained stable (activity variation <15%) when whole blood was stored at 25°C for up to 96 h (**Figure S22a**) and when plasma was stored at 25°C for up to 24 h (**Figure S22b**). This is consistent with the general stability of plasma proteins^29^, which are less susceptible to degradation than RNA or metabolites, particularly because of the presence of endogenous protease inhibitors that suppress protease activity in the blood. We also evaluated the effects of hemolysis, a common source of error in biomarker analysis^30^, and found that its influence was minimal (**Figure S23**). This is likely because the major target enzymes are selectively expressed in the pancreas, reducing the effect of signal contamination from blood cell-derived proteins. This tissue specificity may partly explain the stable assay performance observed across samples collected from multiple hospitals and biobanks. The assay results were not significantly affected by the timing of blood sample collection from healthy subjects within a day (**Figure S24**).

In addition to these pre-analytical variables, we assessed potential pharmacological interference from DPP4 inhibitors, which are commonly prescribed for patients with diabetes^31^, a recognized risk factor for PDAC^32^. When blood samples were treated with a panel of DPP4 inhibitors at clinically relevant maximum plasma concentrations (C_max_), DPP4 activity was partially suppressed by several compounds. Nevertheless, DPP4–DPP4 and DPP4–FAPα remained distinguishable through the preserved FAPα-associated activity, and the resulting DPP4/FAPα score remained largely unchanged (**Figure S25**). Finally, we evaluated the equivalence of plasma and serum samples. Comparable activity patterns were observed for DPP4/FAPα and chymotrypsin. For elastases, neutrophil elastase activity increased in serum relative to plasma, consistent with release during coagulation^33^. However, because the assay clearly distinguished pancreatic and neutrophil elastases, CELA3A activity remained unaffected by sample type (**Figure S26**).

### Data analysis using a random forest classifier and performance of combined markers

We then combined the four biomarker candidates to establish a unified platform to distinguish patients with PDAC from control subjects. For data analysis, the dataset was split into a training set (N = 272; PDAC, N = 60; control subjects, N = 212) and a validation set (N = 181; PDAC, N = 48; control subjects, N = 133), ensuring that the two sets did not include samples from the same institutions (**Table S5**). The remaining 237 non-PDAC samples (other cancers, N = 157; other pancreatic diseases, N = 80) were used to assess cross-reactivity.

A prediction model for PDAC detection was developed using a random forest classifier^34^ with candidate biomarker activities from the training set as input features. The optimal model was selected by hyperparameter tuning using cross-validation. In the validation set, the model yielded an area under the receiver operating characteristic curve (AUROC) of 0.913 (95% CI: 0.857–0.969) (**Figure 3a**). When the analysis was limited to stage I–II PDAC, the AUROC was 0.910 (95% CI: 0.840–0.980) (**Figure 3b**). In the validation cohort, an operating cutoff of 0.369 was selected to achieve the prespecified specificity target of ≥93% among control subjects. At this cutoff, the model showed a sensitivity of 75.0% (95% CI, 60.4%–86.4%) and a specificity of 95.5% (95% CI, 90.4%–98.3%). Notably, the sensitivity was maintained at 74.2% (95% CI: 55.4%–88.1%) even in stage I–II patients. Although the model successfully detected PDAC across all stages, the prediction scores did not strongly correlate with clinical stage (Spearman’s ρ = 0.24, *p* = 0.11), suggesting that the classifier captures disease-associated biochemical alterations that are not simply proportional to tumor burden or stage progression (**Figure 3c, Table S6**).

**Figure 3.**
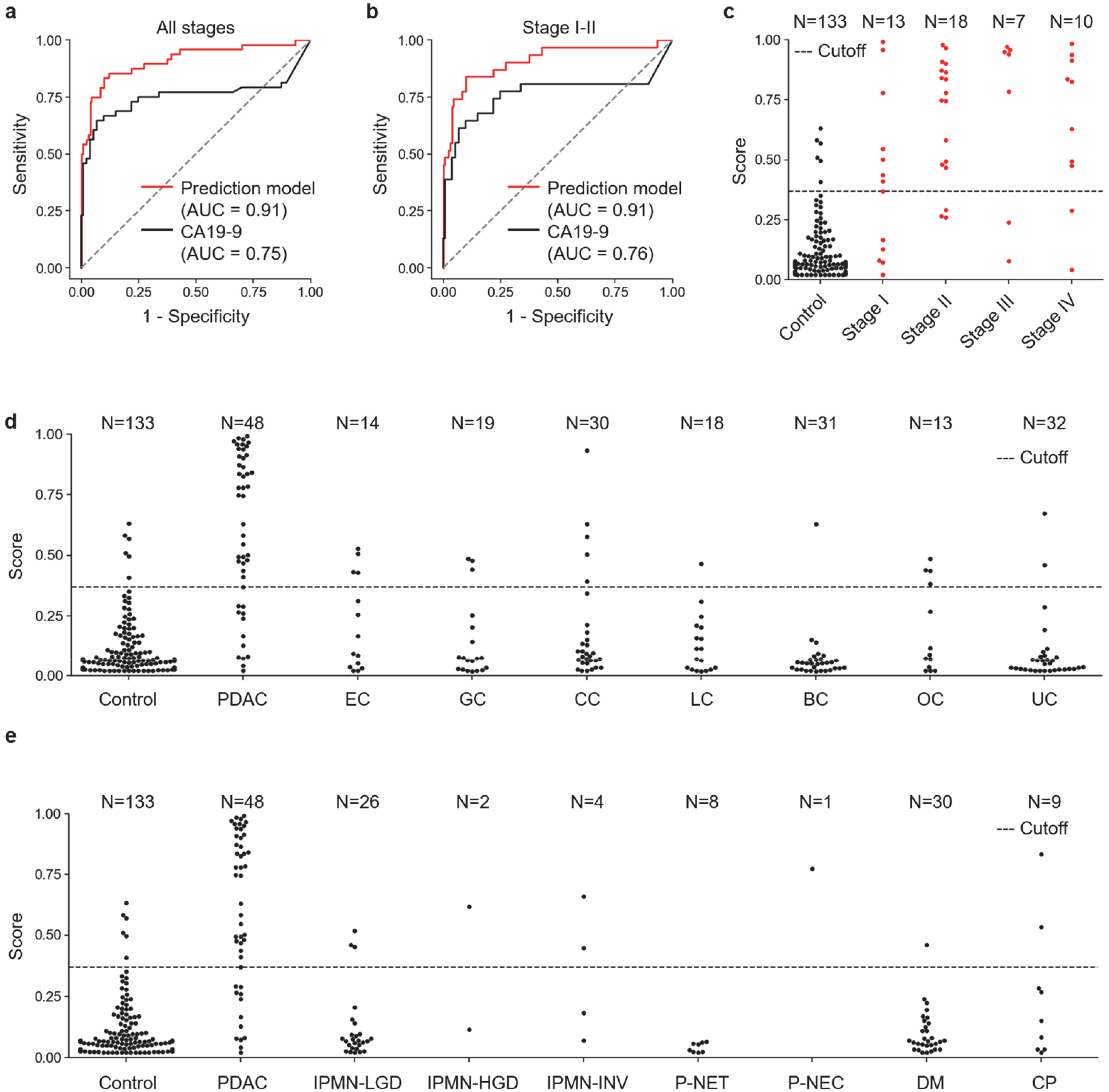
Construction of a random forest classifier–based prediction model for PDAC detection. (a) Receiver operating characteristic (ROC) curves for the combined biomarkers (red) and CA19-9 (black) in the validation set (PDAC, N = 48; control, N = 133). (b) ROC curves for the same analysis restricted to stage I–II PDAC cases (PDAC, N = 31; control, N = 133). (c) Distribution of prediction scores generated by the model for discriminating patients with PDAC from control subjects in the validation set, stratified by disease stage. The dotted line indicates the cutoff value. (d) Distribution of prediction scores generated by the model for control subjects and patients with cancers from various tissues. (e) Distribution of prediction scores generated by the model for control subjects, PDAC, and other diseases related to pancreatic dysfunction. Numbers above the plots indicate sample size. **Abbreviations:** EC, esophageal cancer; GC, gastric cancer; CC, colorectal cancer; LC, lung cancer; BC, breast cancer; OC, ovarian cancer; UC, uterine cancer; IPMN-LGD, intraductal papillary mucinous neoplasm with low-grade dysplasia; IPMN-HGD, IPMN with high-grade dysplasia; IPMN-INV, IPMN with an associated invasive carcinoma; P-NET, pancreatic neuroendocrine tumor; P-NEC, pancreatic neuroendocrine carcinoma; DM, diabetes mellitus; CP, chronic pancreatitis; s.d., standard deviation.

Next, we evaluated the performance of the model using 157 samples from patients with other types of cancer to assess cross-reactivity. The detection rates were 28.6% (4/14) for esophageal, 15.8% (3/19) for gastric, 16.7% (5/30) for colorectal, 5.6% (1/18) for lung, 3.2% (1/31) for breast, 30.8% (4/13) for ovarian, and 6.3% (2/32) for uterine cancers, with an overall detection rate of 12.7% (20/157; **Figure 3d, Table S7**). Most of the false-positive cases exhibited an elevated DPP4–DPP4/DPP4–FAPα ratio.

To further evaluate specificity, we examined the model’s performance on a subset of 80 samples with other pancreatic diseases. The detection rate was 50.0% (3/6) for intraductal papillary mucinous neoplasms (IPMNs) with high-grade dysplasia or associated invasive carcinoma, 11.5% (3/26) for IPMNs with low-grade dysplasia, and 22.2% (2/9) for chronic pancreatitis. Among the individuals with diabetes mellitus, the detection rate was 3.3% (1/30). Notably, no pancreatic neuroendocrine tumors (P-NETs) were detected (0/8), whereas a single available case of pancreatic neuroendocrine carcinoma (P-NEC) was classified as positive using the model (**Figure 3e, Table S7**). One of the two patients with chronic pancreatitis who showed increased scores was in the compensated phase, and the elevation of CTRB2 appeared to be the major contributing factor.

Finally, we compared the performance of the model with that of CA19-9. In the same validation set, CA19-9 yielded an AUROC of 0.753 (95% CI: 0.645–0.861) (**Figure 3a**), which was significantly lower than that of the model (DeLong test, *p* = 0.010). Using the conventional cutoff value of 37 U/mL, CA19-9 showed a sensitivity of 58.3% (95% CI: 43.2%– 72.4%) and a specificity of 94.7% (95% CI: 89.5%–97.9%). In stage I–II PDAC, the sensitivity was 51.6% (95% CI: 33.1%–69.8%) (**Table S6**). Importantly, the model detected 80.0% of CA19-9–negative PDAC cases in stage I-II combined (12/15) and overall (16/20), indicating a substantial complementarity between the two markers. When combined using an either-positive rule (model or CA19-9), the sensitivity increased to 90.3% in stages I–II and 91.7% overall, with a specificity of 90.2% (**Table S6**). Additionally, the detection rate of CA19-9 for other cancers was 20.4% (32/157), which was higher than that observed for the model, indicating reduced specificity in this context (**Table S7**).

### Clinical performance in resectable early-stage PDAC and stage 0 signatures

To test the reproducibility of the performance of our platform for early-stage disease, including carcinoma *in situ* (stage 0) and resectable stage I PDAC, we analyzed an independent test cohort of stage 0 and stage I cases (N = 52) collected at the JA Onomichi General Hospital and Hiroshima University Hospital (**Table S8**). This cohort was retrospectively assembled with a focus on pathologically confirmed stage 0/I lesions, and samples were collected and analyzed independently of the discovery and validation cohorts. Stage 0 was pathologically defined as carcinoma in situ, according to the Japanese classification and registry criteria. Detection at these stages offers the best opportunity for curative surgical intervention and long-term survival; patients diagnosed with stage 0 and stage I (tumor size ≤ 40 mm) achieve 5-year survival rates of 87% and 71%, respectively^4^. However, these early-stage cases currently represent only a small fraction of clinical diagnoses owing to the limited sensitivity of conventional screening modalities and the absence of reliable blood-based biomarkers^35^.

Using the same predefined prediction model and cutoff value established in the preceding analyses without retraining or recalibration, we identified 54.2% (13/24) stage IA patients and 64.3% (9/14) of stage IB patients (**Figure 4a, 4b**). These findings confirmed the robust diagnostic performance for stage I PDAC, which remains the primary target of early intervention. Furthermore, we found that a subset of stage 0 lesions was detectable. Although stage 0 is notoriously difficult to detect using blood-based assays, as exemplified by the limited sensitivity of CA19-9 in this group (7.1%, 1/14), our platform identified 21.4% (3/14) of cases. One case exhibited a particularly robust signal response (notably for CTRB2, **No.6** in **Table S8**). Clinical imaging of this patient revealed focal pancreatic parenchymal atrophy and a stricture of the main pancreatic duct with upstream dilatation, despite the absence of a detectable mass (**Figure 4c–g**). This suggests that even at the carcinoma *in situ* stage, physical alterations such as ductal obstruction can trigger the release of specific enzymes into the circulation. While less pronounced than the aforementioned case, the other two detected stage 0 cases exhibited subtle yet discernible enzymatic activity profiles that remained distinguishable from the control group. Rather than relying on tumor-derived mass antigens, our system may capture broader physiological alterations reflected in the dysregulation of pancreas-specific enzymatic activity that may occur at the onset of neoplastic transformation. Taken together, these findings suggest that, although the enzymatic changes observed in stage IA–IB may already be present in a subset of stage 0 lesions, the enzymatic abnormalities captured by our platform become more pronounced with the progression from carcinoma in situ to invasive stage I disease. From this perspective, the more limited detectability of stage 0 lesions relative to invasive diseases is biologically plausible.

**Figure 4.**
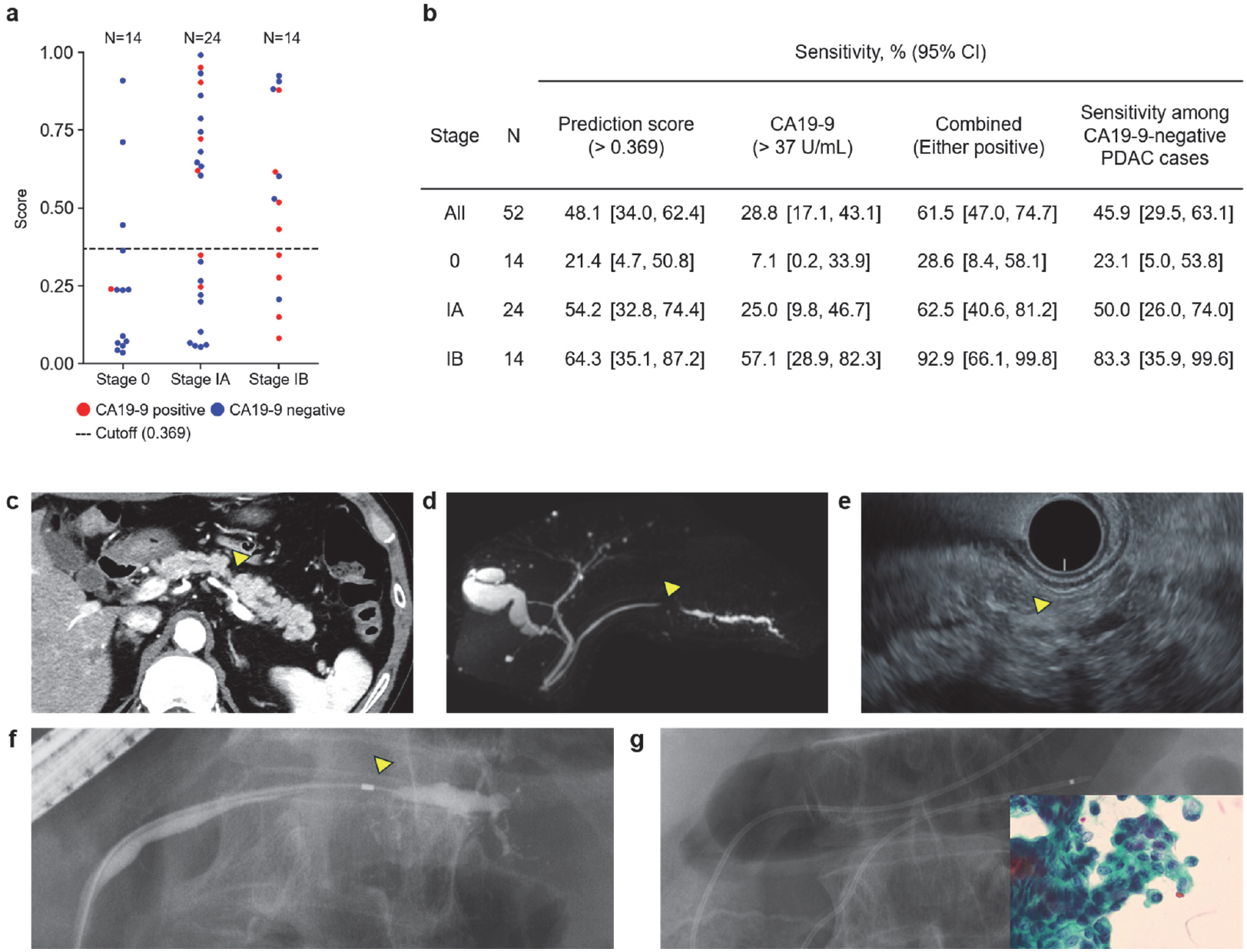
Clinical performance in an independent early-stage cohort. (a) Distribution of prediction scores generated by the model for patients with resectable early-stage PDAC, stratified by disease stage. Blue and red points indicate patients with negative CA19-9 levels (≤37 U/mL) and positive CA19-9 levels (>37 U/mL), respectively. The dashed line indicates the predefined cutoff value of 0.369, determined in the validation cohort. Numbers above the plots indicate the sample size. (b) Sensitivity of the prediction model-derived score, CA19-9, and the combined rule in the test set. Sensitivity is shown overall and stratified by disease stage. The prediction model-derived score was considered positive at >0.369, and CA19-9 was considered positive at >37 U/mL. The combined rule was defined as positive when either the prediction score or CA19-9 was positive. The prediction score in CA19-9-negative PDAC cases column shows the proportion of CA19-9-negative PDAC cases with a positive prediction score. Values are percentages with exact (Clopper–Pearson) 95% confidence intervals in brackets. (c–g) Representative case of a 76-year-old man with stage 0 PDAC who had a positive prediction score. (c) Contrast-enhanced computed tomography demonstrated focal pancreatic parenchymal atrophy in the pancreatic body with dilatation of the distal main pancreatic duct, without an identifiable mass lesion. (d) Magnetic resonance cholangiopancreatography revealed an irregular stricture of the main pancreatic duct in the pancreatic body, with upstream ductal dilatation. (e) Endoscopic ultrasonography showed irregular narrowing of the main pancreatic duct, without a definite mass lesion. (f) Endoscopic retrograde cholangiopancreatography confirmed a stricture of the main pancreatic duct in the pancreatic body, with non-visualization of branch ducts at the stenotic segment. An endoscopic nasopancreatic drainage tube was placed, and serial pancreatic juice aspiration cytologic examination was performed. (g) Cytology from serial pancreatic juice aspiration revealed adenocarcinoma, and the final pathological diagnosis after surgical resection was pancreatic carcinoma in situ. Arrowheads indicate the relevant lesions or ductal abnormalities.

## Discussion

Despite substantial progress in blood-based diagnostics, the clinical application of liquid biopsies for pancreatic ductal adenocarcinoma (PDAC) continues to face several major challenges. These include (1) limited validation across institutionally distinct cohorts^36–40^, (2) variable specificity against non-PDAC malignancies^41,42^, and (3) the inherent difficulty in detecting early-stage disease (stages 0–II), where intervention is most effective^42^. In this study, we aimed to address these challenges by combining single-molecule enzyme activity profiling (SEAP) with tissue-centric biomarker discovery. To strengthen validation across institutionally distinct cohorts (1), we incorporated a large-scale cohort of 690 samples from multiple institutions across Japan and the United States. By employing pooled sample screening to mitigate inter-individual noise, followed by multi-stage individual validation, we identified a biomarker panel that retained discriminative power across distinct clinical settings. To address specificity (2), we prioritized enzymes with pancreas-restricted expression, such as chymotrypsin and pancreatic elastase. While the systemic circulation reflects physiological changes in all organs, our platform focuses on pancreas-specific proteoforms. Based on a tissue-centric biomarker discovery approach, we achieved a high specificity of 95.5% with limited interference from non-PDAC malignancies. The most critical challenge, early-stage detection (3), was addressed through the ability of SEAP to capture the activity of pancreas-specific enzymes at single-molecule resolution. In contrast to tumor-derived markers, such as mutation-based ctDNA, which primarily reflect tumor burden, enzyme activity profiles can report broader biological changes within the pancreatic microenvironment. In the early stages, these alterations are subtle and often fall below the resolution of conventional assays. The sensitivity and single-molecule resolution of SEAP enable the detection of molecular signatures even in stage 0–IB lesions. These findings highlight the potential of capturing premalignant alterations at the level of the tissue microenvironment, rather than the tumor cells themselves. Our approach is conceptually aligned with efforts such as the Human Tumor Atlas Network (HTAN)^43^ and the Pre-Cancer Atlas (PCA)^44^, which aim to resolve tumor evolution and precancerous states in a tissue-level context.

Having established a strategic rationale for our multimarker panel, we next considered the biological context underlying the observed enzymatic changes. The distinct yet complementary behaviors of chymotrypsinogen-derived activities, active CELA3A, and the DPP4–FAPα heterodimer suggest that the panel captures multiple aspects of microenvironmental alterations during early-stage PDAC (**Figure 2f, S2**). Coordinated changes across multiple pancreatic digestive enzymes, including chymotrypsinogen, trypsinogen, carboxypeptidases, and amylases (**Figure S3**), indicate a shared mechanism of entry into the circulation, potentially reflecting the increased leakage from the pancreatic ductal system reported in early-stage disease^45,46^. Consistent with this notion, elevated CTRB2 activity was observed in a stage 0 patient with pancreatic duct constriction identified by endoscopic examination (**Figure 4e**), suggesting a possible association between early ductal abnormalities and circulating enzyme activity. In contrast, CELA3A exhibited a distinct pattern in which the active form decreased without exogenous activation, despite an overall increase in total enzyme levels. This divergence suggests that beyond enzyme release, regulatory mechanisms in the circulation contribute to the observed signal. Given that circulating protease activity is tightly controlled by endogenous inhibitors, including serpins and macroglobulins^25,26^, the CELA3A signal may reflect a shift in the balance between the active and inhibited states (**Figure S28**)^27,28^. Regarding the DPP4-FAPα heterodimer, which we previously reported as a biomarker candidate^9^, its tissue-of-origin is of particular interest in the context of this study. Although DPP4 and FAPα are broadly expressed (**Figure S1**), our observations raise the possibility that the DPP4-FAPα heterodimer may reflect a more pancreas-restricted biology. This interpretation is supported by tissue-level profiling in murine samples, which suggested relative enrichment of the DPP4–FAPα heterodimer in the pancreas (**Figure S29**). Notably, the decrease in heterodimer levels in patients with PDAC did not correlate with CTRB2 levels, suggesting a mechanism distinct from that of digestive enzyme release. In contrast, an increase in the DPP4–DPP4 homodimer was observed in a subset of patients and overlapped with markers associated with biliary obstruction (TNAP and GGT1; **Figure S30**)^47^. These observations suggest that the ratiometric index captures multiple biologically distinct processes, including the loss of pancreas-associated signals and secondary effects related to tumor burden, thereby contributing to improved diagnostic performance. Further mechanistic studies are underway to clarify how these enzymatic signals reflect alterations in the pancreatic microenvironment during early-stage PDAC.

Although our study included a large retrospective multicenter cohort (N = 690) and an independent early-stage test cohort (N = 52), prospective clinical validation is required. Notably, the stage 0 cohort was small and enriched at high-expertise centers; therefore, findings in carcinoma in situ should be considered preliminary while providing clinically relevant proof-of-concept evidence. In addition to expanding the sample size to stage 0, prospective validation in larger, risk-enriched cohorts is ongoing to further define the clinical performance and utility (NCT07605819).

## Conclusion

In conclusion, we developed a blood-based liquid biopsy platform that integrates SEAP with a tissue-centric biomarker discovery strategy. By focusing on the functional activity of pancreas-specific enzymes rather than conventional mass-based antigens, our platform achieved 95.5% specificity and 75.0% sensitivity, while maintaining 74.2% sensitivity in stage I–II PDAC and detecting a subset of lesions at the earliest, potentially curable stages (stage 0– IB). Low detection rates across most non-pancreatic cancer types further support the disease selectivity of this approach. Taken together, these findings, which were supported by multi-institutional validation across diverse cohorts, underscore the reproducibility and potential clinical utility of this technology.

The ability to capture proteoform-level enzymatic dysregulation offers a unique window into the early physiological onset of pancreatic malignancy, thereby addressing a critical gap in current diagnostic capabilities. We anticipate that the SEAP-based platform may facilitate the early detection of PDAC in high-risk populations, but also serve as a versatile framework for developing organ-specific diagnostics for other challenging malignancies. Ultimately, this approach may provide a promising pathway toward shifting the clinical management of PDAC from late-stage palliation to early curative intervention, with the potential to improve long-term patient survival.

## Data availability

The data supporting the findings of this study are available from the corresponding authors upon reasonable request.

## Author contributions

T. Komatsu and Y. Kagami conceived the project. S. S., H. H. and J. H. led the development and optimization of the single-molecule enzyme assays. Experimental data were acquired and analyzed by S. S., H. H., J. H., M. M., and T. Kawaguchi. N. I. and M. Kawana coordinated the large-scale SEAP data acquisition. S. S., K. T., M. M., and T. Komatsu performed the chemical synthesis of fluorogenic probes. M. Kawaguchi and H. N. provided the fluorogenic probes for ENPPs. K. F., Y. U. and T. Komatsu provided the fluorogenic probes for amylases. T. Kobayashi, A. M., and Y. Kodama were responsible for the collection and clinical characterization of blood samples from Kobe University and related hospitals. J. I., Y. I., S. O., and K. H. were responsible for the collection and clinical characterization of blood samples from Hiroshima University Hospital and JA Onomichi General Hospital (Stage 0-I), and for providing the corresponding clinical imaging findings. T. I. and S. T-N. managed the organization and processing of blood samples from multicenter biobanks and hospitals. S. S. and T. M. developed the computational data analysis platform. Interpretation of the data and clinical validation were supervised by S. S., K. H., Y. Kodama, T. Komatsu, and Y. Kagami. The manuscript was written by S. S., T. S., T. Komatsu, and Y. Kagami, with input from all authors.

## Acknowledgements

We thank all employees of Cosomil, Inc. We especially thank Toshihiro Uno for his coordination of the multi-institutional collaboration essential to this study. We also thank Okadai Biobank and JIHS Biobank for providing the study materials, clinical information, and technical support. Blood sample collection was generously supported by Dr. Akinori Shimizu and Dr. Tetsuro Hirano of Department of Gastroenterology, Onomichi General Hospital.

This work was financially supported by the New Energy and Industrial Technology Development Organization (Y. Kagami) (JPNP14012 and JPNP23019).

