## Supplementary Information for "Early detection of pancreatic ductal adenocarcinoma by single-molecule profiling of pancreatic enzyme activities"

**Contents**

**Methods**

**Supplementary tables and figures**

**Supplementary data for preparation of compounds**

**Supplementary references**

### Methods

#### Enzymes

| REAGENT OR RESOURCE | SOURCE | IDENTIFIER |
|---|---|---|
| CTRB1 (*Homo sapiens*) | MedChemExpress | Cat. #HY-P70005<br>Lot #111342 |
| CTRB2 (*Homo sapiens*) | CUSABIO | Cat. #MP006180HU<br>Lot #DA06328k1g0 |
| PRSS1 (*Homo sapiens*) | R&D Systems | Cat. #3848-SE-010<br>Lot #PTF0720121 |
| PRSS2 (*Homo sapiens*) | R&D Systems | Cat. #3586-SE<br>Lot #PKW0423011 |
| PRSS3 (*Homo sapiens*) | R&D Systems | Cat. #3710-SE-010<br>Lot #PDY0221121 |
| CELA3A (*Homo sapiens*) | MyBioSource | Cat. #MBS8248775<br>Lot #NPAX310 |
| Neutrophil elastase (*Homo sapiens*) | Merck | Cat. #324681<br>Lot #4173142 |
| CPB1 (*Homo sapiens*) | R&D Systems | Cat. #2897-ZN<br>Lot #OKA0224031 |
| AMY2A (*Homo sapiens*) | MyBioSource | Cat. #MBS173014<br>Lot #12K9031 |

| | | |
|---|---|---|
| CD13 (*Homo sapiens*) | R&D Systems | Cat. #3815-ZN<br>Lot #POP0223081 |
| GGT1 (*Homo sapiens*) | R&D Systems | Cat. #10977-GT-20<br>Lot #DPRV0124071 |
| ENPP1 (*Homo sapiens*) | R&D Systems | Cat. #6136-EN<br>Lot #TOK0922111 |
| ENPP2 (*Homo sapiens*) | R&D Systems | Cat. #5255-EN<br>Lot #RPZ10213021 |
| ENPP3 (*Homo sapiens*) | Origene | Cat. #TP324818<br>Lot #10CC5C |
| Alkaline phosphatase, recombinant | Roche | Cat. #03359123001<br>Lot #73901638 |
| Trypsin, recombinant | Roche | Cat. #06369880103<br>Lot #80223100 |
| Trypsin, recombinant | Yeasen | Cat. #20416ES60<br>Lot #R6519170 |
| Trypsin, proteomics grade | Roche | Cat. #3708985001<br>Lot #66148823 |
| Enterokinase (*Homo sapiens*) | R&D Systems | Cat. #10438-SE-020<br>Lot #DNDJO124111 |
| TLCK-treated Chymotrypsin | MedChemExpress | Cat. #HY-108910B<br>Lot #617812 |
| Serpin A1 (*Homo sapiens*) | R&D Systems | Cat. #1268-PI-010<br>Lot #HRR0324051 |
| α2-macroglobulin (*Homo sapiens*) | R&D Systems | Cat. #1938-PI-050<br>Lot #MNX1224111 |

**Activation of recombinant enzymes obtained as zymogens**

Recombinant CELA3A, PRSS1, PRSS2, and PRSS3 were obtained as zymogens and pre-activated prior to use. CELA3A (10 μg/mL) was activated with trypsin (1 μg/mL) in HEPES-Na buffer (50 mM, pH 7.4) containing $CaCl_2$ (10 mM), NaCl (150 mM), and CHAPS (0.05% w/v) at 25°C for 2 h. PRSS1, PRSS2, and PRSS3 (10 μg/mL) were activated with enterokinase (1 μg/mL) in HEPES-Na buffer (50 mM, pH 7.4) containing $CaCl_2$ (10 mM), NaCl (150 mM), and CHAPS (0.05% w/v) at 25°C for 1 h. For recombinant CTRB1, CTRB2, and CPB1, the zymogens were activated in situ within the single-molecule assay system by incorporating the appropriate proteolytic enzymes directly into the reaction mixture. Detailed activation protocols for each target are provided in the respective assay sections.

**Plasma samples and ethics statement**

Plasma samples were collected from the Kobe University Hospital (Kobe, Japan) and related institutions

(Asahara Clinic [Akashi, Japan], Kita-Harima Medical Center [Ono, Japan], Japanese Red Cross Kyoto Daini Hospital [Kyoto, Japan], Hotel Okura Kobe Clinic [Kobe, Japan], Yodogawa Christian Hospital [Osaka, Japan]), JA Onomichi General Hospital (Onomichi, Japan), Hiroshima University Hospital (Hiroshima, Japan), biobanks in Japan (Biobank of Japan Institute for Health Security [Tokyo, Japan], Okayama University Hospital Biobank [Okayama, Japan]), biobanks in the US (PrecisionMed, LLC. [California, US], REPROCELL, Inc. [Maryland, US], Dx Biosamples, LLC. [California, US], BioIVT, LLC. [New York, US]), and Cosomil, Inc. (Tokyo, Japan).

Ethical approval for this study was obtained from Kobe University Hospital (B230079), Craif Institutional Review Board (IEF-24012401), PHRF-IRB (24B0001), and Shiba Palace Clinic Ethics Review Committee (161516_rn-42062).

Demographic and clinical characteristics of patients with PDAC and control subjects (sex, age, race, ethnicity, disease stage, tumor location, and institutions) are summarized in **Table S5**. Disease stage was assigned according to the Union for International Cancer Control (UICC) TNM classification, 8th edition.

**Study design and cohorts**

Plasma samples were collected from patients with pancreatic, esophageal, gastric, colorectal, lung, breast, ovarian, and uterine cancers, intraductal papillary mucinous neoplasms (IPMNs) with low-grade dysplasia, high-grade dysplasia, and an associated invasive carcinoma, pancreatic neuroendocrine tumors, pancreatic neuroendocrine carcinoma, chronic pancreatitis, diabetes mellitus, and control subjects with no cancers or pancreatic diseases. For patients with malignancies, samples were collected before any treatment. For PDAC cases classified as stage III or below, inclusion was restricted to those with histologically confirmed adenocarcinoma or anaplastic carcinoma. Throughout the study, measurements and signal extraction from each blood sample were performed in a blinded manner without access to patient information.

To screen the biomarker candidates, we used 32 patients with PDAC and 96 non-cancer control subjects from biobanks in Japan.

To evaluate the detection performance of the prediction model, we split the collected samples into a training set (N = 272; PDAC, N = 60; control subjects, N = 212) and a validation set (N = 181; PDAC, N = 48; control subjects, N = 133), ensuring that the two sets did not include samples from the same institutions. The training set consisted of samples collected at the Kobe University Hospital and related institutions, PrecisionMed, LLC., and REPROCELL, Inc., whereas the validation set comprised samples from biobanks in Japan, Dx Biosamples, LLC., and BioIVT, LLC. The remaining 237 samples (other cancers, N = 157; other pancreatic diseases, N = 80), collected at biobanks in Japan, were used to evaluate specificity. To test the reproducibility of the performance of the constructed prediction model, especially for early-stage PDAC, we analyzed an independent test cohort of stage 0 and stage I cases (N = 52) collected at JA Onomichi General Hospital and Hiroshima University Hospital (**Figure 1d**, **Table S1**).

**Plasma/serum preparation**

For samples collected at Kobe University Hospital (Kobe, Japan) and related institutions, blood was drawn into EDTA-containing tubes (Terumo, VP-NA050K). The collected blood samples were then centrifuged at 4°C, 3,000 rpm for 15 min to separate plasma.

For samples collected at Japan Institute for Health Security, blood was drawn into EDTA-containing tubes (Terumo, VP-NA070K). The collected blood samples were then centrifuged at 15°C, 3,000 rpm for 10 min to separate plasma.

For samples collected at Okayama University Hospital, blood was drawn into EDTA-containing tubes. The samples were first centrifuged at 2,200 × g for 10 min at 20°C to separate plasma. The resulting plasma was transferred to microtubes and subjected to a second centrifugation at 16,000 × g for 10 min at 20°C. The supernatant was then collected for analysis.

For samples collected at PrecisionMed LLC., blood was drawn into EDTA-containing tubes. The samples were first centrifuged at 1,200 RCF for 10 min at room temperature to separate plasma. The resulting plasma was transferred to microtubes and subjected to a second centrifugation at 1,200 RCF for 10 min at room temperature. The supernatant was then collected for analysis.

For samples collected at REPROCELL, Inc., blood was drawn into EDTA-containing tubes. The collected blood samples were then centrifuged at 4°C, 2,000 rpm for 10 min to separate plasma.

For samples collected at Dx Biosamples, LLC., blood was drawn into EDTA-containing tubes. The collected blood samples were then centrifuged at room temperature, 1,500 × g for 10 min to separate plasma.

For samples collected at a blood collection facility operated by BioIVT, LLC., blood was drawn into EDTA-containing tubes. The collected blood samples were then centrifuged at 5°C at 5,000 × g for 15 min to separate plasma.

For samples collected at hospitals affiliated with BioIVT, LLC., detailed information regarding sample processing was not available.

For samples collected at JA Onomichi General Hospital, peripheral venous blood was collected under standardized conditions into serum-separating tubes. The blood samples were allowed to clot at room temperature for 30 min and then centrifuged at 1,500 × g for 10 min at 4°C. The serum was carefully transferred to avoid cellular contamination, aliquoted into pre-labeled cryovials, and stored at −80°C until analysis. Multiple aliquots were prepared for each sample to avoid repeated freeze–thaw cycles.

For samples collected at Hiroshima University Hospital, peripheral venous blood was collected under standardized conditions into EDTA-containing tubes. The blood samples were centrifuged at 3,500 rpm for 10 min at 4°C, and the plasma supernatant was carefully transferred to avoid contamination with the buffy coat or cellular components. The plasma was then subjected to a second centrifugation at 12,000 rpm for 10 min at 4°C to remove residual cellular debris and platelets. The clarified plasma was aliquoted into pre-labeled cryovials and stored at −80°C until analysis. Multiple aliquots were prepared for each sample to avoid repeated freeze–thaw cycles.

**Plasma preparation for stability test**

Plasma samples used for stability testing were collected at Cosomil, Inc. Blood was drawn into EDTA-containing tubes. The collected blood samples were generally centrifuged at room temperature at 2,200 × g for 10 min to separate plasma. In cases where alternative conditions were applied as part of the experimental design, the specific procedures are described in the **Stability analysis** section.

**Stability analysis**

**Stability in whole blood**

To evaluate the stability of enzyme activities in whole blood, samples collected into EDTA-containing tubes were kept at 25°C without centrifugation. Plasma was separated by centrifugation after 0 h (immediately after collection), 24 h, 48 h, and 96 h. The resulting plasma was promptly frozen at −80°C until analysis. Enzyme activities were measured in plasma samples obtained at each time point (**Figure S22a**). For the summary analysis shown in **Figure 2g**, the relative change at 96 h from 0 h was calculated using the mean of three technical replicates for each biological sample and summarized as mean ± s.d. across biological replicates.

**Stability in plasma**

To evaluate the stability after plasma separation, blood samples were centrifuged immediately after collection, and plasma was aliquoted. Plasma was frozen at −80°C either immediately (0 h), after 8 h at 25°C, or after 24 h at 25°C. Enzyme activities were measured in plasma samples obtained at each time point (**Figure S22b**). For the summary analysis shown in **Figure 2g**, the relative change at 24 h from 0 h was calculated using the mean of three technical replicates for each biological sample and summarized as mean ± s.d. across biological replicates.

**Effect of hemolysis**

To evaluate the effect of hemolysis, whole blood from each donor was divided into two aliquots. One aliquot was subjected to mechanical stress by passing it 10 times through a 27G needle to induce hemolysis, while the other was left untreated. Both samples were subsequently centrifuged to obtain plasma, which was frozen at −80°C until analysis. Enzyme activities were measured in plasma from hemolyzed and non-hemolyzed samples. Hemoglobin concentrations in the same plasma samples were measured as an indicator of hemolysis using a Hemoglobin Assay Kit (BioChain Institute, Z5030026) according to the manufacturer's instructions (**Figure S23**). For the summary analysis shown in **Figure 2g**, the relative change between hemolyzed and non-hemolyzed samples was calculated for each biological sample and summarized as mean ± s.d. across biological replicates.

**Diurnal variation**

To evaluate potential diurnal variation, eight donors were assigned to either a morning group (AM group, N = 4) or an afternoon group (PM group, N = 4). The AM group fasted from 22:00 on the day prior to blood collection, while the PM group fasted from 07:00 on the day of collection. For each donor, blood was collected at 09:00, 11:00, 13:00, and 15:00. Donors in the AM group consumed lunch between the 11:00 and 13:00 collections, whereas those in the PM group remained fasting until the 15:00 collection. Enzyme activities were measured in plasma obtained at each time point (**Figure S24**). For the summary analysis shown in **Figure 2g**, relative changes from the 09:00 value were calculated at 11:00, 13:00, and 15:00 for each donor, and the mean of these three relative changes was used as the representative value for each donor. These values were then summarized as mean ± s.d. across biological replicates.

**Calculation of change in values**

For all stability analyses, relative change was calculated as:

$$\text{Relative change (\%)} = \frac{\text{value}_{\text{after}} - \text{value}_{\text{before}}}{\text{value}_{\text{before}}} \times 100.$$

Here, $\text{value}_{\text{before}}$ denotes the initial value for each comparison (0 h, non-hemolyzed, or 09:00), and $\text{value}_{\text{after}}$ denotes the corresponding later or treated value. When technical replicates were available, their mean was used for the calculation.

**Comparison of plasma and serum measurements**

To compare enzyme activity measurements between plasma and serum, paired blood samples were collected from three donors at the same time into EDTA-Na-containing tubes for plasma preparation and serum-separating tubes for serum preparation. Enzyme activities were measured in paired plasma and serum samples (**Figure S26**).

**Preparation of mouse tissue lysate**

Ethical approval for the study using animals was obtained from Animal Care and Use Committee of the University of Tokyo (P4-21, P31-9). Preparation of tissue lysate followed the reported protocol with minor modification[1]. Mice (C57BL/6JJcl, 6-week-old, male) were sacrificed and tissues (brain, liver, lung, heart, spleen, pancreas, kidney, testis) were isolated and flash frozen in liquid nitrogen. Frozen tissue samples were weighed and placed into a dounce tissue homogenizer with 2 mL ice-cold PBS (pH 7.4) with 10 strokes of the pestle. Each sample was centrifuged at 2,000 × g (4°C, 5 min) to separate insoluble material. The lysates were collected, aliquoted for single-use, flash frozen in liquid nitrogen, and stored at -80°C. Protein concentration of the lysate was measured by Bradford assay using BSA as a calibrant.

**Preparation of mouse blood**

Ethical approval for the study using animals was obtained from Animal Care and Use Committee of The University of Tokyo (P4-21, P31-9). Mice (C57BL/6JJcl, 6-week-old, male) were euthanized and the blood was collected through an inferior vena cava into 1.5 mL tube containing 1.5 μL heparin (Yoshindo Inc., Japan). The collected blood sample was centrifuged at 1,700 × g (4°C for 15 min) for plasma separation. The plasma samples were collected, aliquoted, flash frozen in liquid nitrogen, and stored at -80°C.

**Tissue-dominance analysis**

Gene expression data were obtained from the GTEx v11 gene-level TPM matrix and the corresponding GTEx sample-attribute table. Samples were matched using GTEx sample identifiers. Tissue labels were defined from the “Sample Tissue” annotation (SMTS), and only samples present in both the expression matrix and the attribute table were retained. All analyses were performed on $\log2(TPM + 1)$-transformed expression values.

For a given target tissue, let $T\_med$ denote the median expression across samples assigned to the target tissue. For non-target expression, we defined $O\_med$ as the median across all non-target samples, $O\_max$ as the maximum of the tissue-wise non-target medians, and $O\_q95$ as the 95$^{th}$ percentile across all non-target samples. Using these quantities, we defined the following gene-level scores: $margin = T_{\mathrm{med}} - O_{\mathrm{max}}$, $margin\_med = T_{\mathrm{med}} - O_{\mathrm{med}}$, and $gap\_q95 = T_{\mathrm{med}} - O_{\mathrm{q95}}$.

The primary tissue-dominance score was $margin$, which measures separation from the most highly expressing non-target tissue. Genes were ranked using a margin-first rule: descending margin, followed by higher $T\_med$, lower $O\_max$, and lower $O\_q95$. This ranking defined the primary ordering of tissue-dominant candidate genes.

After target-tissue-specific candidate genes had been identified for a given target tissue, their expression patterns were compared across all tissues using a grouped tissue-level view. For each selected gene, the median expression within each tissue was computed on the $\log2(TPM + 1)$ scale and displayed side by side across tissues. This comparison was used to confirm that expression in the target tissue exceeded that in non-target tissues.

For analyses restricted to protein-coding genes, Ensembl gene identifiers were matched to GENCODE v47 gene annotations after removing version suffixes. The protein-coding filter was applied after score calculation. Optional gene-symbol annotation was added separately for readability and did not affect scoring or ranking.

The resulting ranked list was used to prioritize candidate genes for downstream biomarker design and experimental evaluation. To verify the tissue-specificity of the top candidates, their expression profiles were visualized using normalized nTPM data from the Human Protein Atlas (v24), which integrates curated GTEx datasets.

The code used for tissue-dominance analysis, candidate ranking, and group-view visualization of tissue-level expression is available at https://github.com/mizuno-group/tissue-dominance.

**Fluorescence microscopy**

Fluorescence images were acquired using a fluorescence microscope (Ti2, Nikon) equipped with a 20× dry objective lens (Plan Apo 20×), sCMOS camera (ORCA-Fusion C14440, Hamamatsu Photonics), white LED illumination unit (X-Cite Xylis, Opto Science), a moving stage and a rotating stage. The rotating stage was custom-made to hold a Simoa Disc (Quanterix Corporation, MA, USA) and rotate it using a motorized rotation stage (OSMS-60YAW, Opto Sigma). For continuous monitoring of multiple device images, the rotating stage rotated the disc by 15° for each acquisition. Images were acquired in tile scan mode of 3×4 (with overlap of 1%) with perfect focus. The assay was performed using a solution containing IR-dye 800 (30 μM) as the internal standard, and the focus was adjusted using its fluorescence. The excitation and emission filters used were FITC (mirror = 505 nm, Ex. = 465-495 nm, Em. = 512-558 nm), mCherry (mirror = 600 nm, Ex. = 550-590 nm, Em. = 608-683 nm), Cy5 (mirror = 660 nm, Ex. = 590-650 nm, Em. = 663-738 nm) and Cy7 (mirror = 743 nm, Ex. = 743 nm, Em. = 767 nm), respectively. Fluorescence signals of sHMRG- and sTG-based probes were acquired with FITC filter, those of sSiR-, dsSiR-, sTM-, and resorufin-based probes were acquired with mCherry or Cy5 filter, and those of IR-dye 800 were acquired with Cy7 filter.

**Single-molecule enzyme activity assay**

Loading of biological samples to microdevice (Simoa Disc, Quanterix Corporation, MA, USA) was conducted using either the SR-X system (Quanterix Corporation)[2] or manual pipetting[3,4], as specified for each target enzyme (**Table S4**).

For loading conducted with the SR-X system, biological samples were prepared at 2× the final working concentration in 96-well plates (Quanterix Corporation), and probe solutions were prepared at 2× concentration in Narrow-Mouth Bottles (Thermo Scientific). The biological sample and the probe solution were mixed and introduced into a Simoa Disc followed by sealing with Simoa Sealing Oil (Quanterix Corporation) via the SR-X system using the standard assay protocol of SR-X. After automated processing by the SR-X system, an additional volume of Simoa Sealing Oil was manually applied to the lane inlets of the disc, and the lane outlets were sealed with adhesive stickers to prevent evaporation of the reaction mixture.

For manual loading, both the biological samples and the probe solutions were prepared at 2× concentration in microcentrifuge tubes (Eppendorf, 30108116). A total of 17.5 µL of each solution was mixed, and 30 µL of the resulting mixture was dispensed into a Simoa Disc using a P-100 or P-200 micropipette equipped with wide-bore pipette tips (Thermo Scientific [QSP, T118R-Q]). Subsequently, 65 µL of FC-70 (Tokyo Chemical Industry) was added to the disc to seal the device. After completion of these steps, additional FC-70 was manually applied to the lane inlets, and the lane outlets were sealed with adhesive stickers to prevent evaporation of the reaction mixture.

The device was incubated in an incubator (AS ONE Corporation, FCI-280G) at 25°C for the indicated time. Fluorescence images of the microdevice were acquired using a fluorescence microscope.

**Image processing**

Images were processed using the GA3 module of NIS Elements software (Nikon). First, all fluorescence images were background-corrected using a rolling ball correction (diameter = 3 µm). Then, ROIs were selected by bright spot detection using either FITC and mCherry filters or FITC and Cy5 filters (diameter = 3 µm). To exclude irregular fluorescent spots derived from fluorescent debris or air bubbles, the detected ROIs were dilated, and ROIs that overlapped after dilation were removed. The remaining ROIs were further filtered by size and shape, and only ROIs with fill area <40 and circularity >0.8 were retained for subsequent intensity quantification. The fluorescence signals were calculated from a 3×3 pixel area centered on each ROI as the sum of all nine pixels, excluding the maximum and minimum values. All analyzable chambers were chosen by bright spot detection using Cy5 or Cy7 filter (diameter = 4 µm) to calculate lambda (signal numbers per all analyzed chambers) values. For elastase measurements, additional ROIs were generated to calculate fluorescence intensities from chambers adjacent to each primary ROI. Specifically, the original ROI was dilated and intersected with the set of all chambers, and the original ROI was then subtracted to yield adjacent chamber ROIs. Their fluorescence intensities were calculated in the same manner as for the primary ROIs.

**Cluster analysis**

Detailed procedures are provided below for the markers used to build the prediction model. For the remaining enzyme assays, the cluster analysis approaches are summarized in **Table S4.**

**DPP4/FAPα**

Fluorescence intensity distributions from the FITC and mCherry channels were transformed into histograms on a logarithmic scale, and the histograms were smoothed prior to parametric modeling. Events with mCherry intensity < 10,000, corresponding to FAPα-related signals, were fitted with a single Gaussian component, which defined the boundary separating the FAPα homodimer population from the other populations. For the remaining events, the FITC channel was fitted with a two-component Gaussian mixture. The FITC-low and FITC-high components were assigned to DPP4 homodimer- and DPP4-FAP$\alpha$ heterodimer related populations, respectively, and vertical boundaries were defined accordingly. For these DPP4-selected events, the mCherry channel was further fitted with a two-component Gaussian mixture. The separation between the two fitted peaks was evaluated using the normalized peak-to-peak distance, defined as follows:

$$D_m = \frac{|\mu_1 - \mu_2|}{\sqrt{{\sigma_1}^2 + {\sigma_2}^2}}$$

where $\mu_1$ and $\mu_2$ are the centers of the two fitted Gaussian components, and ${\sigma_1}^2$ and ${\sigma_2}^2$ are their variances, respectively. When $D_m < 1$, the two mCherry components were considered insufficiently separated. In this case, populations related to DPP4 homodimers and DPP4-FAP$\alpha$ heterodimers defined by the FITC channel were each fitted separately with a single Gaussian component in the mCherry direction, and horizontal boundaries were determined from these individual fits. When $D_m \geq 1$, the horizontal boundaries were determined from the two-component Gaussian mixture fitted to the mCherry distribution.

The vertical and horizontal boundaries were then combined to generate rectangular gates on the two-dimensional scatter plot, corresponding to FAPα homodimers, DPP4 homodimers, and DPP4–FAPα heterodimers. DPP4/FAPα was defined as the ratio of DPP4 homodimer events to the sum of DPP4 homodimer and DPP4–FAPα heterodimer events.

**Chymotrypsinogen**

Fluorescence intensity data from the FITC and mCherry channels were analyzed as two-dimensional scatter plots. Because clustering based on individual measurements was unstable owing to the limited number of events per measurement, all analyzable events from the training and validation sets (**Table S1**) were pooled into a single dataset for clustering. This unsupervised clustering step was used only to define fluorescence-intensity-based event populations and was independent of the diagnostic labels used for classifier training. The two-channel coordinates were transformed into polar features, consisting of angle and radius, in the two-dimensional fluorescence intensity space. The polar features were standardized to zero mean and unit variance and clustered with k-means using five clusters.

Each cluster was assigned to populations derived from either CTRB1 or CTRB2 based on recombinant

CTRB1 and CTRB2 measurements (**Figure S5c**). Among the five clusters, the cluster with the smallest angle, located in the upper-left region of the FITC–mCherry scatter plot, was assigned to the CTRB1-derived population. Three clusters with intermediate angles were assigned to CTRB2-derived populations. The cluster with the largest angle, located in the lower-right region of the FITC–mCherry scatter plot, was also assigned to a CTRB2-derived population, denoted as CTRB2-B, based on its consistency with recombinant CTRB2 measurements (**Figure S5c**).

For data that were not included in the original pooled dataset, including newly acquired measurements, the corresponding events were added to the original pooled dataset, and k-means clustering was repeated using the combined dataset. This approach was used because the number of events in the pooled dataset was substantially larger than that in each added dataset, allowing stable cluster assignment while minimizing changes in cluster centers and boundaries. The cluster labels obtained from the combined clustering were then extracted for the added data and used for sample-level quantification.

Quantification was performed for each sample individually. After cluster assignment in the pooled or combined dataset, the number of events assigned to each CTRB1- or CTRB2-derived population was counted separately for each sample. The CTRB2 value was calculated as the number of signals assigned to CTRB2-derived populations divided by the total number of analyzed chambers. The CTRB1/CTRB2 ratio, representing the fraction of CTRB1-assigned events among chymotrypsinogen-derived fluorescent events, was defined as the number of CTRB1-assigned signals divided by the sum of CTRB1- and CTRB2-assigned signals, including CTRB2-B.

**Elastase**

Fluorescence intensity data from the FITC and Cy5 channels were analyzed as two-dimensional scatter plots. FITC and Cy5 intensities were extracted from the primary ROIs defined during image processing. Additional ROIs were generated from chambers adjacent to each primary ROI to estimate the local Cy5 background intensity around each event. Events were retained when the difference between the Cy5 intensity of the primary ROI and the local Cy5 background intensity estimated from adjacent chambers was <300.

For the retained events, the FITC intensity distribution was analyzed to identify CELA3A-derived signals. A fixed FITC intensity threshold of 12,578 was used as the boundary, and events with FITC intensity higher than this threshold were assigned to the CELA3A population. The CELA3A value was calculated as the number of CELA3A-assigned signals divided by the total number of analyzed chambers.

**Prediction model construction**

The prediction model was built using four single-molecule enzyme activity measurements (DPP4/FAPα, CTRB2, CTRB1/CTRB2, and CELA3A) as input variables. No normalization or scaling was applied, and raw values were used directly as classifier inputs. Cohort definitions and sample allocation to training and validation sets are provided in the **Study design and cohorts** section. A random forest classifier was trained using scikit-learn, with hyperparameters optimized by grid search and five-fold cross-validation using the area under the receiver operating characteristic curve (AUROC) as the objective function. The final model was constructed with the following hyperparameters: n_estimators = 100, max_depth = 3, max_features = 'sqrt', criterion = 'entropy', and ccp_alpha = 0.01, with random_state = 42 used for the main analysis. Model performance was evaluated in the validation cohort using AUROC and using sensitivity and specificity at an operating cutoff selected in the validation cohort to meet the prespecified specificity target of ≥93% among control subjects. This cutoff was 0.369 and was subsequently applied without modification to the independent early-stage test cohort. To evaluate robustness to algorithmic randomness, the classifier was retrained using multiple random seeds while keeping the training and validation cohorts fixed. ROC curves and summary performance metrics showed only minor variation across seeds (**Figure S27**), indicating that the model performance was stable with respect to seed selection.

**Measurement of CA19-9 values of plasma samples**

CA19-9 levels were measured using commercially available ELISA kits for CA19-9 (Lumipulse Prestro CA19-9; Fujirebio Inc.)[5,6].

**Measurement of Elastase-1 values of plasma samples**

Elastase-1 concentrations were measured by latex agglutination turbidimetric immunoassay using the Iatro IRE1® II reagent kit on a JCA-BM8040 automated clinical chemistry analyzer (JEOL Ltd., Tokyo, Japan), according to the manufacturer's instructions.

**Statistics**

No statistical method was used to predetermine sample size. Sample sizes were based on sample availability and are indicated in the corresponding figure legends and tables. Unless otherwise stated, all statistical tests were two-sided, and $P < 0.05$ was considered statistically significant.

Binomial proportions, including sensitivity, specificity, detection rates and marker-positive rates, are reported with exact (Clopper–Pearson) 95% confidence intervals. This approach was used for performance estimates of the prediction model and CA19-9 reported in the main text and supplementary tables.

AUROC values and their 95% confidence intervals were estimated using DeLong's method. Pairwise comparison of AUROC values between the prediction model and CA19-9 was performed using DeLong's test.

Correlations were assessed using Spearman's rank correlation coefficient ($\rho$). Spearman's $\rho$ was used to evaluate associations between enzyme activity–related measurements and between ordinal UICC disease stage and continuous marker or classifier scores, because these variables were not assumed to follow a normal distribution and included skewed distributions and potential outliers.

For paired two-group comparisons, including paired measurements of the same samples under different assay conditions, paired t-tests were used.

Details of the statistical tests used for individual figures and tables are provided in the corresponding legends where applicable.

**Implementation**

All analyses were implemented in Python 3.13 using standard scientific libraries. Prediction model construction was performed using scikit-learn 1.8.0, and data processing was performed using pandas 3.0.3. The complete code for **Cluster analysis** and **Prediction model construction** is available at https://github.com/cosomil/early-detection-PDAC.

### Supplementary tables and figures

**Table S1.** Distribution of study samples by contributing institution and cohort.

| Institution | Cohort | Training/ Validation/ Test set | PDAC | Non-cancer control | Healthy subjects | Other cancers | Other pancreatic diseases |
|---|---|---|---|---|---|---|---|
| Kobe University Hospital | JP bank | Training | 50 | 0 | 0 | 0 | 0 |
| Asahara Clinic | JP bank | Training | 0 | 0 | 29 | 0 | 0 |
| Kita-Harima Medical Center | JP bank | Training | 0 | 0 | 16 | 0 | 0 |
| Japanese Red Cross Kyoto Daini Hospital | JP bank | Training | 0 | 0 | 6 | 0 | 0 |
| Hotel Okura Kobe Clinic | JP bank | Training | 0 | 0 | 96 | 0 | 0 |
| Yodogawa Christian Hospital | JP bank | Training | 0 | 0 | 3 | 0 | 0 |
| Biobank of Japan Institute for Health Security | JP bank/ Screening | Validation | 13 | 96 | 22 | 157 | 75 |
| Okayama University Hospital Biobank | Screening | Validation | 19 | 0 | 0 | 0 | 5 |
| PrecisionMed, LLC. | US bank | Training | 4 | 62 | 0 | 0 | 0 |
| REPROCELL, Inc. | US bank | Training | 6 | 0 | 0 | 0 | 0 |
| Dx Biosamples, LLC. | US bank | Validation | 16 | 0 | 0 | 0 | 0 |
| BioIVT, LLC. | US bank | Validation | 0 | 15 | 0 | 0 | 0 |
| JA Onomichi General Hospital | Early-stage PDAC | Test | 33 | 0 | 0 | 0 | 0 |
| Hiroshima University Hospital | Early-stage PDAC | Test | 19 | 0 | 0 | 0 | 0 |

**Table S2.** Identification of pancreas-specific enzymes based on transcriptomic analysis. A list of enzymes demonstrating pancreas-specific expression profiles was generated using GTEx transcriptome databases. Top 50 protein-coding genes with high pancreatic specificity are shown. Yellow highlighted proteins are enzymes and enzyme activity-regulating proteins that are potentially covered in our screening.

| | Gene | Name | Score† |
|---|---|---|---|
| 1 | CLPS | Colipase | 12.6 |
| 2 | PNLIP | Pancreatic lipase | 12.6 |
| 3 | CPA1 | Carboxypeptidase A1 | 12.3 |
| 4 | CTRB2 | Chymotrypsin B2 | 12.0 |
| 5 | CTRB1 | Chymotrypsin B1 | 11.7 |
| 6 | PRSS1 | Serine protease 1 | 11.5 |
| 7 | PNLIPRP1 | Pancreatic lipase related protein 1 | 10.9 |
| 8 | INS | Insulin | 10.9 |
| 9 | CELA3A | Chymotrypsin like elastase 3A | 10.9 |
| 10 | AMY2A | Amylase alpha 2A | 10.9 |
| 11 | CTRC | Chymotrypsin C | 10.8 |
| 12 | CELA2A | Chymotrypsin like elastase 2A | 10.6 |
| 13 | CELA3B | Chymotrypsin like elastase 3B | 10.6 |
| 14 | PLA2G1B | Phospholipase A2 group 1B | 10.5 |
| 15 | GP2 | Glycoprotein 2 | 9.4 |
| 16 | CELA2B | Chymotrypsin like elastase 2B | 8.7 |
| 17 | CUZD1 | CUB and zona pellucida like domains 1 | 8.4 |
| 18 | PRSS2 | Serine protease 2 | 8.3 |
| 19 | PNLIPRP2 | Pancreatic lipase related protein 2 | 8.2 |
| 20 | CPB1 | Carboxypeptidase B1 | 7.8 |
| 21 | SYCN | Syncollin | 7.2 |
| 22 | SERPINI2 | Serpin I2 | 7.0 |
| 23 | AQP8 | Aquaporin 8 | 6.8 |
| 24 | CPA2 | Carboxypeptidase A2 | 6.7 |
| 25 | RBPJL | Recombination signal binding protein for immunoglobulin kappa J region like | 6.5 |
| 26 | CTRL | Chymotrypsin like | 6.4 |
| 27 | CEL | Carboxyl ester lipase | 6.3 |
| 28 | REG1B | Regenerating family member 1 beta | 5.9 |
| 29 | PRSS3 | Serine protease 3 | 5.7 |
| 30 | CLPSL1 | Colipase like 1 | 5.6 |
| 31 | KLK1 | Kallikrein 1 | 5.3 |
| 32 | AQP12B | Aquaporin 12B | 5.1 |
| 33 | REG1A | Regenerating family member 1 alpha | 4.5 |
| 34 | AQP12A | Aquaporin 12A | 4.5 |
| 35 | SPINK1 | Serine peptidase inhibitor Kazal type 1 | 4.0 |
| 36 | KIRREL2 | Kirre like nephrin family adhesion molecule 2 | 4.0 |
| 37 | REG3G | Regenerating family member 3 gamma | 3.9 |
| 38 | ERP27 | Endoplasmic reticulum protein 27 | 3.8 |
| 39 | AMY2B | Amylase alpha 2B | 3.7 |
| 40 | PM20D1 | Peptidase M20 domain containing 1 | 3.7 |
| 41 | IAPP | Islet amyloid polypeptide | 3.3 |
| 42 | PTF1A | Pancreas associated transcription factor 1a | 3.3 |
| 43 | PDX1 | Pancreatic and duodenal homeobox 1 | 3.1 |
| 44 | AMY1B | Amylase alpha 1B | 3.0 |
| 45 | SCTR | Secretin receptor | 3.0 |
| 46 | PPY | Pancreatic polypeptide | 3.0 |
| 47 | LEFTY1 | Left-right determination factor 1 | 2.8 |
| 48 | PDIA2 | Protein disulfide isomerase family A member 2 | 2.7 |
| 49 | BHLHA15 | Basic helix-loop-helix family member a15 | 2.6 |
| 50 | GUCA1C | Guanylate cyclase activator 1C | 2.5 |

†Score = $median_{pancreas}$ - $max(median_{other\_tissues})$, where $median_{pancreas}$ represents the $log_2(TPM+1)$ transformed median expression level in pancreatic tissue, and $max(median_{other\_tissues})$ denotes the maximum median expression level observed among all other analyzed tissues.

**Table S3.** Summary of SEAP assays performed in this study. Assays conducted under distinct pH conditions (acidic, neutral, or basic) were categorized as separate entries to account for the functional diversity of enzymes, such as those with neutral pH optima versus lysosomal enzymes that exhibit acidic pH optima. For the initial screening, activities were examined using either pooled blood samples or a set of 128 independent blood samples.

| Category | No. of assays tested |
|---|---|
| Rhodamine†-based protease probes[2,7] | 108 |
| ProTide-based carboxypeptidase probes[8] | 12 |
| Glycosidase probes | 5 |
| Esterase/phosphatase probes[3,9] | 4 |
| FRET-based exopeptidase probes[10] | 28 |
| Oxidoreductases/coupled assays[4] | 15 |
| ABP-based single-molecule assays[11] | 10 |
| Total | 182 |

†Probes are the amide-modified probes of sHMRG, sSiR, or dsSiR.

**Table S4.** Assay conditions used to measure the target enzymes. The filter wavelength is provided in the **Fluorescence microscopy** section.

| Enzyme | Probe | Conc. (μM) | Sample dilution | Reaction buffer | Load | Incubation Time | Filter | Exposure | Cluster analysis | Ref. |
|---|---|---|---|---|---|---|---|---|---|---|
| Dipeptidyl peptidase IV/ Fibroblast activation protein α | Ac-GP-sHMRG | 30 | x5,000 | HEPES-Na buffer (100 mM, pH 7.4) containing $MgCl_2$ (1 mM), $CaCl_2$ (1 mM), DTT (100 μM), BSA (1% w/v), and Triton X-100 (150 μM) | SR-X | 4 h | FITC | 20%, 1 s | Gaussian Fitting (See Methods) | 2 |
| | EP-sSiR | 30 | | | | | mCherry | 5%, 1 s | | |
| Aminopeptidase | Y-sHMRG | 30 | x5,000 | HEPES-Na buffer (100 mM, pH 7.4) containing $MgCl_2$ (1 mM), $CaCl_2$ (1 mM), DTT (100 μM), and Triton X-100 (150 μM) | SR-X | 2 h | FITC | 5%, 1 s | Manual gating | 2 |
| | A-sSiR | 30 | | | | | mCherry | 5%, 1 s | | |
| γ-glutamyl transferase | γGlu-sHMRG | 30 | x500 | MES-Na buffer (100 mM, pH 5.5) containing $MgCl_2$ (1 mM), $CaCl_2$ (1 mM), DTT (100 μM), and Triton X-100 (150 μM) | Manual | 20 h | FITC | 20%, 1 s | Manual gating | - |
| | γGlu-sSiR | 30 | | | | | mCherry | 20%, 1 s | | |
| Alkaline phosphatase | sTG-mPhos | 25 | x1,000 | DEA-HCl buffer (1 M, pH 9.3) containing $MgCl_2$ (1 mM), $CaCl_2$ (1 mM), $ZnCl_2$ (20 μM), and Triton X-100 (150 μM) | SR-X | 50 min | FITC | 5%, 500 ms | Manual gating | 9 |
| | sTM-Phos | 25 | | | | | mCherry | 5%, 300 ms | | |
| Ectonucleotide pyrophosphatase/ phosphodiesterase 1/3 | sTG-mADP | 30 | x200 | Tris-HCl buffer (100 mM, pH 9.0) containing HEPES-Na (25 mM), $MgCl_2$ (1 mM), $CaCl_2$ (1 mM), NaCl (75 mM), $ZnCl_2$ (250 μM), ALP (80 ng/mL), CHAPS (0.05% w/v), and Triton X-100 (150 μM) | SR-X | 1 h | FITC | 5%, 500 ms | Manual gating | - |
| | sTM-dCMP | 30 | | | | | mCherry | 5%, 200 ms | | |
| Ectonucleotide pyrophosphatase/ phosphodiesterase 2 | sTG-diPrAMP | 30 | x5,000 | Tris-HCl buffer (100 mM, pH 9.0) containing $MgCl_2$ (5 mM), NaCl (150 mM), BSA (1% w/v), and Triton X-100 (150 μM) | Manual | 2 h | FITC | 20%, 500 ms | Manual gating | - |
| Chymotrypsinogen | Suc-RPY-sHMRG | 30 | x2,500 | HEPES-Na buffer (100 mM, pH 7.4) containing $MgCl_2$ (1 mM), $CaCl_2$ (5 mM), NaCl (1 M), DTT (100 μM), Trypsin (25 μg/mL), and Triton X-100 (150 μM) | Manual | 20 h | FITC | 20%, 1 s | K-means clustering (See Methods) | - |
| | Suc-AAVY-dsSiR | 30 | | | | | mCherry | 20%, 1 s | | |
| Chymotrypsin | Suc-RPY-sHMRG | 30 | x100 | HEPES-Na buffer (125 mM, pH 7.4) containing $MgCl_2$ (1 mM), $CaCl_2$ (10 mM), NaCl (1.075 M), DTT (100 μM), CHAPS (0.05% w/v), and Triton X-100 (150 μM) | Manual | 20 h | FITC | 50%, 1 s | Manual gating | - |
| | Suc-AAVY-dsSiR | 30 | | | | | Cy5 | 100%, 1 s | | |
| Trypsinogen | Ac-RSPK-sHMRG | 30 | x2,500 | HEPES-Na buffer (100 mM, pH 7.4) containing $MgCl_2$ (1 mM), $CaCl_2$ (5 mM), NaCl (50 mM), DTT (100 μM), bestatin (100 μM), enterokinase (10 ng/mL), TLCK-treated chymotrypsin (16 μg/mL), and Triton X-100 (150 μM) | Manual | 20 h | FITC | 20%, 1 s | Manual gating | - |
| | Ac-IGGR-PABC-dsSiR | 10 | | | | | mCherry | 10%, 1 s | | |
| Trypsin | Ac-RSPK-sHMRG | 30 | x250 | HEPES-Na buffer (90 mM, pH 7.4) containing $MgCl_2$ (0.8 mM), $CaCl_2$ (4 mM), NaCl (70 mM), DTT (80 μM), CHAPS (0.02% w/v), and Triton X-100 (120 μM) | Manual | 20 h | FITC | 20%, 1 s | Manual gating/ Local background ratio thresholding | - |
| | Ac-IGGR-PABC-dsSiR | 10 | | | | | mCherry | 5%, 400 ms | | |

| Enzyme | Probe | Conc. (μM) | Sample dilution | Reaction buffer | Load | Incubation Time | Filter | Exposure | Cluster analysis | Ref. |
|---|---|---|---|---|---|---|---|---|---|---|
| Carboxypeptidase A | sTG-PHBA-P(Et)-AY | 10 | x2,500 | Tris-HCl buffer (100 mM, pH 8.5) containing $CaCl_2$ (5 mM), $ZnCl_2$ (10 μM), DTT (100 μM), Trypsin (1.67 μg/mL), and Triton X-100 (150 μM) | Manual | 24 h | FITC | 20%, 1 s | Manual gating | 8 |
| | Res-PHBA-P(Et)-GL | 10 | | | | | mCherry | 20%, 1 s | | |
| Carboxypeptidase B | sTG-PHBA-P(Et)-GR | 10 | x20,000 | HEPES-Na buffer (100 mM, pH 7.4) containing NaCl (100 mM), $ZnCl_2$ (100 μM), DTT (100 μM), Trypsin (0.21 μg/mL), and Triton X-100 (150 μM) | Manual | 20 h | FITC | 10%, 1 s | Manual gating | - |
| | Res-PHBA-P(Et)-AK | 10 | | | | | mCherry | 10%, 1 s | | |
| Amylase | sTG-Amy | 1 | x1,000 | HEPES-Na buffer (100 mM, pH 7.4) containing $MgCl_2$ (1 mM), $CaCl_2$ (5 mM), NaCl (100 mM), and Triton X-100 (150 μM) | Manual | 24 h | FITC | 20%, 500 ms | Local background ratio thresholding | - |
| Elastase | Suc-AAPAbu-PABC-sHMRG | 10 | x100 | HEPES-Na buffer (125 mM, pH 7.4) containing $MgCl_2$ (1 mM), $CaCl_2$ (10 mM), NaCl (75 mM), DTT (100 μM), CHAPS (0.05% w/v), and Triton X-100 (150 μM) | Manual | 24 h | FITC | 30%, 1 s | Local background difference thresholding/ fixed FITC thresholding (See Methods) | - |
| | Suc-AAPV-PABC-dsSiR | 15 | | | | | Cy5 | 100%, 3 s | | |
| Proelastase | Suc-AAPAbu-PABC-sHMRG | 10 | x1,000 | HEPES-Na buffer (125 mM, pH 7.4) containing $MgCl_2$ (1 mM), $CaCl_2$ (10 mM), NaCl (75 mM), DTT (100 μM), Trypsin (4 μg/mL), and Triton X-100 (150 μM) | Manual | 24 h | FITC | 30%, 1 s | Manual gating | - |
| | Suc-AAPV-PABC-dsSiR | 15 | | | | | Cy5 | 100%, 3 s | | |

**Table S5.** Clinical characteristics of plasma samples in this study for the training and validation sets.

| | Training set | | Validation set | |
|---|---|---|---|---|
| | PDAC (N=60) | Control (N=212) | PDAC (N=48) | Control (N=133) |
| Sex | | | | |
| Male | 34 | 127 | 29 | 60 |
| Female | 26 | 85 | 19 | 73 |
| Age (y) | | | | |
| Mean (s.d.) | 67.4 (11.0) | 66.4 (9.5) | 70.0 (8.3) | 59.4 (14.3) |
| Median | 67 | 66 | 70.5 | 57 |
| Min, Max | 37, 86 | 46, 87 | 45, 86 | 28, 92 |
| Race | | | | |
| Asian | 50 | 150 | 32 | 118 |
| White | 10 | 44 | 12 | 8 |
| Black or African American | 0 | 8 | 4 | 5 |
| Unknown | 0 | 10 | 0 | 2 |
| Hispanic or Latino ethnic group | | | | |
| Yes | 0 | 12 | 0 | 2 |
| No | 50 | 150 | 48 | 118 |
| Unknown | 10 | 50 | 0 | 13 |
| Stage | | | | |
| I | 15 | - | 13 | - |
| IA | 7 | - | 8 | - |
| IB | 8 | - | 5 | - |
| II | 10 | - | 18 | - |
| IIA | 0 | - | 3 | - |
| IIB | 10 | - | 15 | - |
| III | 15 | - | 7 | - |
| IV | 20 | - | 10 | - |
| Location | | | | |
| Pancreatic head | 31 | - | 26 | - |
| Pancreatic body | 14 | - | 11 | - |
| Pancreatic tail | 5 | - | 6 | - |
| Unknown | 10 | - | 5 | - |
| Institutions | - Kobe University Hospital<br>- Asahara Clinic<br>- Kita-Harima Medical Center<br>- Japanese Red Cross Kyoto Daini Hospital<br>- Hotel Okura Kobe Clinic<br>- Yodogawa Christian Hospital<br>- PrecisionMed, LLC.<br>- REPROCELL, Inc. | | - Biobank of Japan Institute for Health Security<br>- Okayama University Hospital Biobank<br>- Dx Biosamples, LLC.<br>- BioIVT, LLC. | |

**Table S6.** Sensitivity and specificity of the prediction model-derived score, CA19-9, and the combined rule in the validation set. Sensitivity is shown overall and stratified by disease stage, and specificity is shown for control subjects. The prediction model-derived score was considered positive at >0.369, and CA19-9 was considered positive at >37 U/mL. The combined rule was defined as positive when either the prediction score or CA19-9 was positive. The column "Prediction score in CA19-9-negative PDAC cases" shows the proportion of CA19-9-negative PDAC cases with a positive prediction score. Values are percentages with exact (Clopper–Pearson) 95% confidence intervals in brackets.

| | | | Sensitivity, % (95% CI) | | | |
|---|---|---|---|---|---|---|
| | Stage | N | Prediction score (> 0.369) | CA19-9 (> 37 U/mL) | Combined (Either positive) | Sensitivity among CA19-9-negative PDAC cases |
| PDAC | All | 48 | 75.0 [60.4, 86.4] | 58.3 [43.2, 72.4] | 91.7 [80.0, 97.7] | 80.0 [56.3, 94.3] |
| | I | 13 | 61.5 [31.6, 86.1] | 38.5 [13.9, 68.4] | 84.6 [54.6, 98.1] | 75.0 [34.9, 96.8] |
| | IA | 8 | 75.0 [34.9, 96.8] | 25.0 [3.2, 65.1] | 75.0 [34.9, 96.8] | 66.7 [22.3, 95.7] |
| | IB | 5 | 40.0 [5.3, 85.3] | 60.0 [14.7, 94.7] | 100 [47.8, 100] | 100 [15.8, 100] |
| | II | 18 | 83.3 [58.6, 96.4] | 61.1 [35.7, 82.7] | 94.4 [72.7, 99.9] | 85.7 [42.1, 99.6] |
| | IIA | 3 | 100 [29.2, 100] | 66.7 [9.4, 99.2] | 100 [29.2, 100] | 100 [2.5, 100] |
| | IIB | 15 | 80.0 [51.9, 95.7] | 60.0 [32.3, 83.7] | 93.3 [68.1, 99.8] | 83.3 [35.9, 99.6] |
| | I-II | 31 | 74.2 [55.4, 88.1] | 51.6 [33.1, 69.8] | 90.3 [74.2, 98.0] | 80.0 [51.9, 95.7] |
| | III | 7 | 71.4 [29.0, 96.3] | 57.1 [18.4, 90.1] | 85.7 [42.1, 99.6] | 66.7 [9.4, 99.2] |
| | IV | 10 | 80.0 [44.4, 97.5] | 80.0 [44.4, 97.5] | 100 [69.2, 100] | 100 [15.8, 100] |
| | | | Specificity, % (95% CI) | | | |
| Control | | 133 | 95.5 [90.4, 98.3] | 94.7 [89.5, 97.9] | 90.2 [83.9, 94.7] | |

**Table S7.** Positive rates of the score (>0.369) and CA19-9 (>37 U/mL) in samples from individuals with other pancreatic diseases and cancers evaluated for specificity. Values are shown as percentages with exact (Clopper–Pearson) 95% confidence intervals in brackets. **Abbreviations:** IPMN-LGD, intraductal papillary mucinous neoplasm with low-grade dysplasia; IPMN-HGD, IPMN with high-grade dysplasia; IPMN-INV, IPMN with an associated invasive carcinoma; P-NET, pancreatic neuroendocrine tumor; P-NEC, pancreatic neuroendocrine carcinoma.

| | | Positivity rate, % (95% CI) | |
|---|---|---|---|
| | N | Prediction score (> 0.369) | CA19-9 (> 37 U/mL) |
| Other pancreatic diseases | 80 | | |
| IPMN-HGD | 2 | 50.0 [1.3, 98.7] | 50.0 [1.3, 98.7] |
| IPMN-INV | 4 | 50.0 [6.8, 93.2] | 50.0 [6.8, 93.2] |
| IPMN-LGD | 26 | 11.5 [2.4, 30.2] | 15.4 [4.4, 34.9] |
| Diabetes mellitus | 30 | 3.3 [0.1, 17.2] | 6.7 [0.8, 22.1] |
| Chronic pancreatitis | 9 | 22.2 [2.8, 60.0] | 11.1 [0.3, 48.2] |
| P-NET | 8 | 0.0 [0.0, 36.9] | 0.0 [0.0, 36.9] |
| P-NEC | 1 | 100 [2.5, 100] | 0.0 [0.0, 97.5] |
| Other cancers | 157 | 12.7 [8.0, 19.0] | 20.4 [14.4, 27.5] |
| Esophageal cancer | 14 | 28.6 [8.4, 58.1] | 21.4 [4.7, 50.8] |
| Gastric cancer | 19 | 15.8 [3.4, 39.6] | 26.3 [9.1, 51.2] |
| Colorectal cancer | 30 | 16.7 [5.6, 34.7] | 3.3 [0.1, 17.2] |
| Lung cancer | 18 | 5.6 [0.1, 27.3] | 16.7 [3.6, 41.4] |
| Breast cancer | 31 | 3.2 [0.1, 16.7] | 12.9 [3.6, 29.8] |
| Ovarian cancer | 13 | 30.8 [9.1, 61.4] | 46.2 [19.2, 74.9] |
| Uterine cancer | 32 | 6.3 [0.8, 20.8] | 31.3 [16.1, 50.0] |

**Table S8.** Details of patients with PDAC from JA Onomichi General Hospital and Hiroshima University Hospital. -, not available. CA19-9 values below the lower limit of quantification were recorded as 1 U/mL. **Abbreviation:** MPD, main pancreatic duct.

| No. | Sex | Age | Stage | Prediction score | CA19-9 (U/mL) | Tumor size (mm) | Location | MPD Dilatation | MPD Stenosis | Pancreatic atrophy | DPP4/FAPα (%) | CTRB2 (λ value, x$10^{-3}$) | CTRB1/CTRB2 (%) | CELA3A (λ value, x$10^{-4}$) |
|---|---|---|---|---|---|---|---|---|---|---|---|---|---|---|
| 1 | F | 60 | 0 | 0.035 | 7.2 | - | Body | Yes | Yes | Yes | 57.2 | 4.00 | 32.6 | 3.54 |
| 2 | F | 79 | 0 | 0.066 | 27.7 | - | Body | Yes | Yes | Yes | 48.1 | 2.19 | 29.2 | 5.02 |
| 3 | F | 75 | 0 | 0.042 | 5.2 | - | Body | No | Yes | Yes | 52.6 | 1.65 | 33.0 | 3.15 |
| 4 | F | 66 | 0 | 0.445 | 1 | - | Tail | Yes | Yes | Yes | 60.3 | 8.34 | 21.3 | 3.25 |
| 5 | M | 75 | 0 | 0.236 | 27.5 | - | Tail | Yes | Yes | Yes | 61.2 | 2.43 | 33.3 | 5.40 |
| 6 | M | 76 | 0 | 0.909 | 5.9 | - | Tail | Yes | Yes | Yes | 55.2 | 17.3 | 12.1 | 3.83 |
| 7 | F | 73 | 0 | 0.237 | 3.4 | - | Body | Yes | Yes | Yes | 59.7 | 3.42 | 29.8 | 1.70 |
| 8 | F | 76 | 0 | 0.237 | 1 | - | Body | Yes | Yes | Yes | 52.5 | 3.30 | 19.9 | 3.41 |
| 9 | M | 73 | 0 | 0.088 | 4.9 | - | Body | Yes | Yes | Yes | 60.3 | 2.86 | 30.2 | 7.53 |
| 10 | F | 83 | 0 | 0.057 | 9.5 | - | Body | Yes | Yes | Yes | 50.9 | 2.87 | 30.2 | 8.42 |
| 11 | F | 68 | 0 | 0.071 | 6.9 | - | Body | Yes | Yes | Yes | 53.9 | 3.33 | 26.9 | 8.63 |
| 12 | F | 82 | 0 | 0.363 | 4 | - | Tail | Yes | Yes | Yes | 60.4 | 3.45 | 30.9 | 1.48 |
| 13 | M | 84 | 0 | 0.711 | 1 | - | Tail | Yes | Yes | Yes | 67.6 | 9.56 | 22.9 | 1.74 |
| 14 | M | 81 | 0 | 0.239 | 41.7 | 18 | Tail | Yes | Yes | Yes | 61.4 | 4.22 | 31.2 | 4.25 |
| 15 | F | 78 | IA | 0.102 | 12.9 | 10 | Body | Yes | Yes | No | 54.6 | 3.07 | 24.3 | 2.21 |
| 16 | F | 68 | IA | 0.903 | 103 | 11 | Head | Yes | Yes | No | 64.1 | 164 | 7.42 | 14.1 |
| 17 | F | 79 | IA | 0.619 | 51.4 | 10 | Head | Yes | Yes | Yes | 66.3 | 50.1 | 37.2 | 940 |
| 18 | F | 79 | IA | 0.199 | 4.5 | 12 | Body | Yes | Yes | Yes | 54.9 | 7.44 | 16.0 | 5.69 |
| 19 | M | 82 | IA | 0.348 | 69.7 | 20 | Body | Yes | Yes | Yes | 58.0 | 7.47 | 16.3 | 3.00 |
| 20 | F | 76 | IA | 0.722 | 40.9 | 8 | Head | Yes | Yes | Yes | 64.9 | 9.38 | 20.3 | 4.04 |
| 21 | F | 80 | IA | 0.932 | 9.5 | 13 | Body | Yes | Yes | Yes | 65.4 | 16.6 | 9.61 | 5.86 |
| 22 | F | 78 | IA | 0.680 | 30.7 | 6 | Tail | Yes | Yes | Yes | 72.2 | 2.27 | 31.7 | 1.42 |
| 23 | M | 71 | IA | 0.059 | 20.9 | 16 | Tail | Yes | Yes | Yes | 46.9 | 2.29 | 28.7 | 8.37 |
| 24 | F | 70 | IA | 0.861 | 7.5 | 16 | Body | Yes | Yes | Yes | 76.0 | 12.9 | 20.7 | 3.32 |
| 25 | M | 80 | IA | 0.057 | 14.1 | 15 | Tail | Yes | Yes | Yes | 54.2 | 4.52 | 29.4 | 5.85 |
| 26 | M | 75 | IA | 0.265 | 1 | 20 | Head | Yes | Yes | Yes | 51.0 | 4.21 | 16.5 | 1.65 |
| 27 | M | 83 | IA | 0.246 | 48.3 | 12 | Head | No | Yes | No | 59.9 | 10.8 | 26.2 | 8.53 |
| 28 | M | 83 | IA | 0.220 | 17.2 | 8 | Body | Yes | Yes | Yes | 59.3 | 4.51 | 25.6 | 4.40 |
| 29 | M | 67 | IA | 0.066 | 2.1 | 13 | Body | Yes | Yes | Yes | 53.3 | 9.80 | 28.6 | 15.0 |
| 30 | F | 56 | IA | 0.991 | 19 | 12 | Body | No | Yes | No | 70.7 | 43.2 | 13.1 | 1.79 |
| 31 | M | 62 | IA | 0.634 | 18.8 | 10 | Head | Yes | Yes | Yes | 58.6 | 17.2 | 25.9 | 76.9 |
| 32 | F | 66 | IA | 0.646 | 1 | 15 | Tail | No | No | Yes | 72.0 | 1.09 | 45.5 | 1.63 |
| 33 | F | 68 | IA | 0.327 | 11.8 | 4 | Body | Yes | Yes | Yes | 57.1 | 11.3 | 19.2 | 6.61 |
| 34 | F | 81 | IA | 0.604 | 10.9 | 13 | Body | Yes | Yes | Yes | 68.1 | 9.66 | 13.0 | 35.8 |
| 35 | F | 81 | IA | 0.053 | 15.4 | 18 | Body | No | No | Yes | 57.5 | 3.89 | 31.1 | 9.28 |
| 36 | M | 77 | IA | 0.744 | 7.1 | 15 | Head | Yes | Yes | Yes | 64.1 | 4.70 | 14.5 | 2.36 |
| 37 | M | 87 | IA | 0.951 | 207 | 7 | Head | Yes | Yes | No | 64.1 | 34.3 | 14.8 | 7.36 |
| 38 | F | 85 | IA | 0.787 | 22.4 | 19 | Body | Yes | Yes | Yes | 75.1 | 33.8 | 22.8 | 29.7 |
| 39 | F | 65 | IB | 0.081 | 43.5 | 28 | Tail | Yes | No | No | 57.4 | 3.03 | 27.9 | 6.44 |
| 40 | M | 79 | IB | 0.431 | 105 | 21 | Body | Yes | Yes | Yes | 61.5 | 3.27 | 45.2 | 1.43 |
| 41 | M | 58 | IB | 0.348 | 69.5 | 30 | Head | Yes | Yes | No | 61.2 | 5.13 | 26.4 | 1.77 |
| 42 | M | 77 | IB | 0.882 | 14.2 | 20 | Head | Yes | Yes | No | 57.6 | 100 | 8.27 | 7.70 |
| 43 | F | 56 | IB | 0.906 | 2.2 | 24 | Body | Yes | Yes | No | 55.2 | 27.8 | 12.0 | 5.87 |
| 44 | M | 71 | IB | 0.925 | 15.1 | 21 | Body | Yes | Yes | Yes | 65.9 | 57.1 | 6.96 | 8.38 |
| 45 | F | 76 | IB | 0.879 | 115 | 23 | Head | Yes | Yes | No | 55.7 | 18.9 | 15.1 | 7.64 |
| 46 | M | 67 | IB | 0.206 | 9.7 | 21 | Body | Yes | Yes | Yes | 64.0 | 5.52 | 36.1 | 8.53 |
| 47 | M | 70 | IB | 0.616 | 59 | 25 | Head | Yes | - | No | 90.6 | 7.13 | 24.6 | 2.18 |
| 48 | F | 76 | IB | 0.149 | 49.6 | 25 | Tail | No | No | No | 44.4 | 4.60 | 17.2 | 4.94 |
| 49 | M | 87 | IB | 0.518 | 144 | 26 | Head | Yes | Yes | No | 69.9 | 4.35 | 25.5 | 3.93 |
| 50 | M | 73 | IB | 0.276 | 86 | 40 | Head | Yes | Yes | No | 55.3 | 2.11 | 26.7 | 1.02 |
| 51 | F | 87 | IB | 0.529 | 15 | 21 | Head | No | No | No | 65.6 | 6.04 | 23.4 | 4.00 |
| 52 | M | 81 | IB | 0.602 | 1 | 37 | Tail | No | No | Yes | 66.4 | 11.8 | 28.7 | 3.92 |

**Table S9.** DPP4 inhibitors and maximum plasma concentrations used for the spike-in interference assay shown in **Figure S25**. The inhibitor panel was selected to cover all nine DPP4 inhibitors approved in Japan. $C_{max}$ values were obtained from the corresponding package inserts. When multiple dose levels were reported, the highest dose was used. When $C_{max}$ was reported as mean ± s.d., the mean value was used to calculate the final spike-in concentration. $C_{max}$, maximum plasma concentration; s.d., standard deviation.

| DPP4 inhibitor | Molecular weight (g/mol) | Reported $C_{max}$ in package insert | Final spike-in concentration (nM) | Ref. |
|---|---|---|---|---|
| Alogliptin | 339.39 | 193.3±32.5 ng/mL | 570 | [12] |
| Anagliptin | 383.45 | 1040±291 ng/mL | 2712 | [13] |
| Linagliptin | 472.54 | 23.1 nM | 23 | [14] |
| Omarigliptin | 398.43 | 750 nM | 750 | [15] |
| Saxagliptin | 333.43 | 18.7±3.4 ng/mL | 56 | [16] |
| Sitagliptin | 407.31 | 944±307 nM | 944 | [17] |
| Teneligliptin | 426.58 | 382.40 ± 89.83 ng/mL | 896 | [18] |
| Trelagliptin | 475.47 | 619.4±77.3 ng/mL | 1303 | [19] |
| Vildagliptin | 303.40 | 538±149 ng/mL | 1773 | [20] |

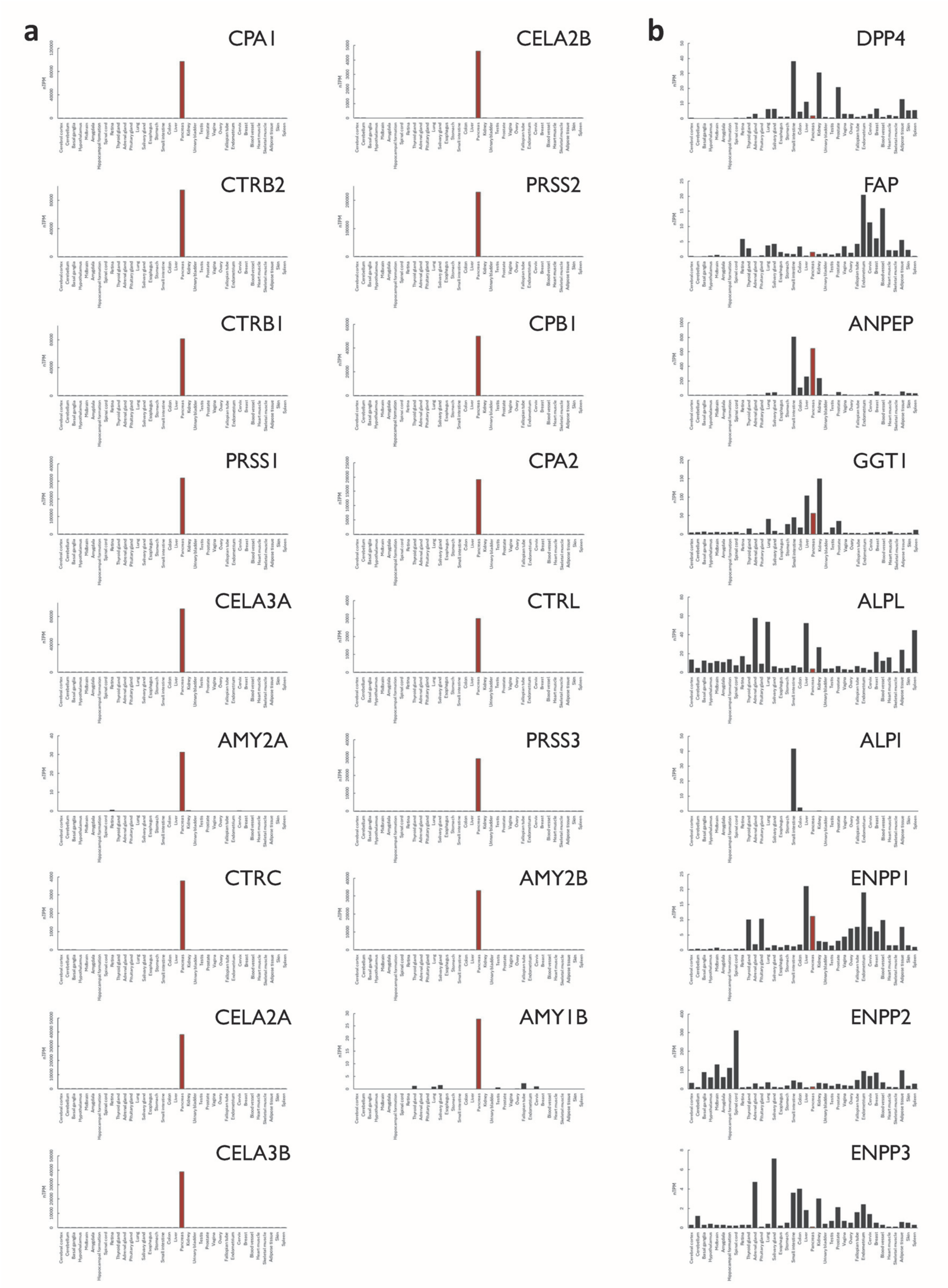


**Figure S1.** Tissue expression patterns of enzymes evaluated in this study. (a) Tissue expression patterns of enzymes categorized as “pancreas-specific” based on the list of **Table S2**. (b) Tissue expression patterns of representative enzymes in random screening using panels of SEAP assays. Red bar = pancreas.

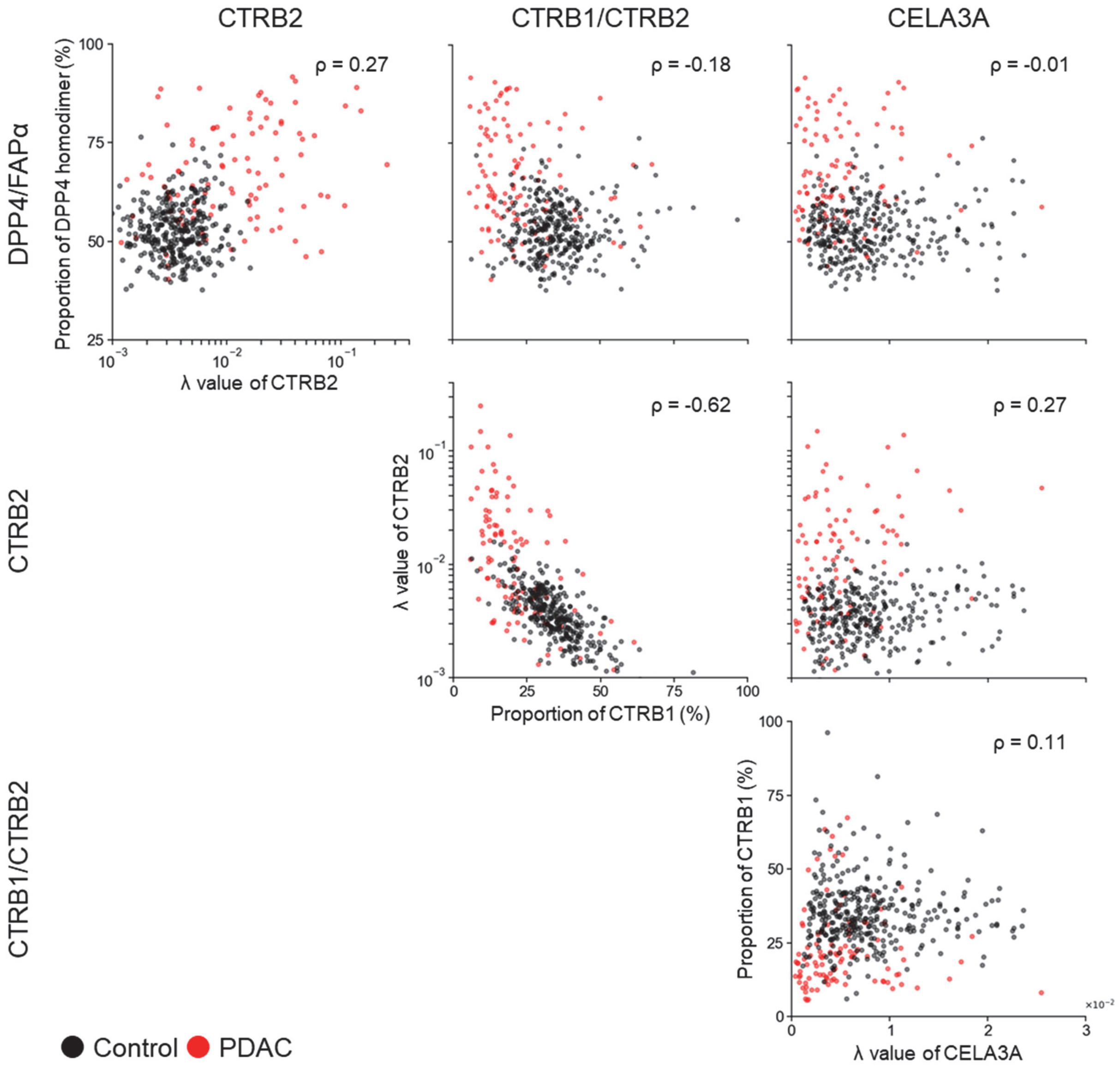


**Figure S2.** Correlations of the four markers (DPP4/FAPα, CTRB2, CTRB1/CTRB2, and CELA3A). Scatter plots were generated using patients with PDAC and control subjects from the combined training and validation sets (PDAC, N = 108; Control, N = 345). Black dots represent control subjects and red dots represent PDAC cases. ρ, Spearman's rank correlation coefficient calculated using PDAC samples only.

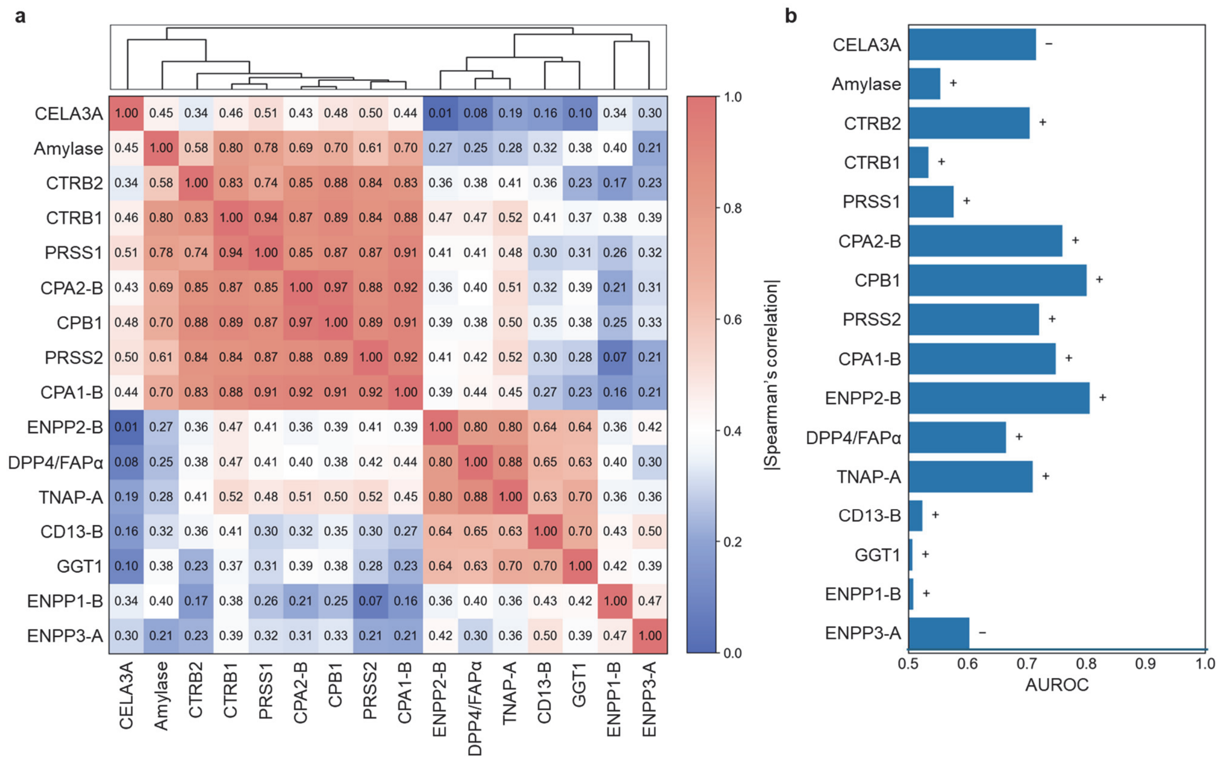


**Figure S3.** Pairwise correlations among all tested biomarkers and their univariate diagnostic performance. (a) Heatmap of absolute Spearman rank correlation coefficients (|ρ|) among candidate biomarker features, calculated using PDAC samples in the screening cohort (N = 32). Rows and columns are displayed in the same order, determined by hierarchical clustering (top dendrogram; average linkage) using a distance metric defined as $d = 1 - |\rho|$. Numeric values in each cell indicate the corresponding |ρ|. (b) Univariate diagnostic performance of each biomarker feature for discriminating PDAC from controls in the screening cohort (PDAC, N = 32; control, N = 96), expressed as the AUROC. Symbols next to the bars indicate the direction of change in PDAC relative to controls: "+" indicates higher values in PDAC, and "–" indicates lower values in PDAC. The clusters corresponding to each biomarker feature are defined in the relevant **Supplementary Figures S4, S5, S9, S11, S12, S13, S14, S15, S16, S17, S18, and S20**.

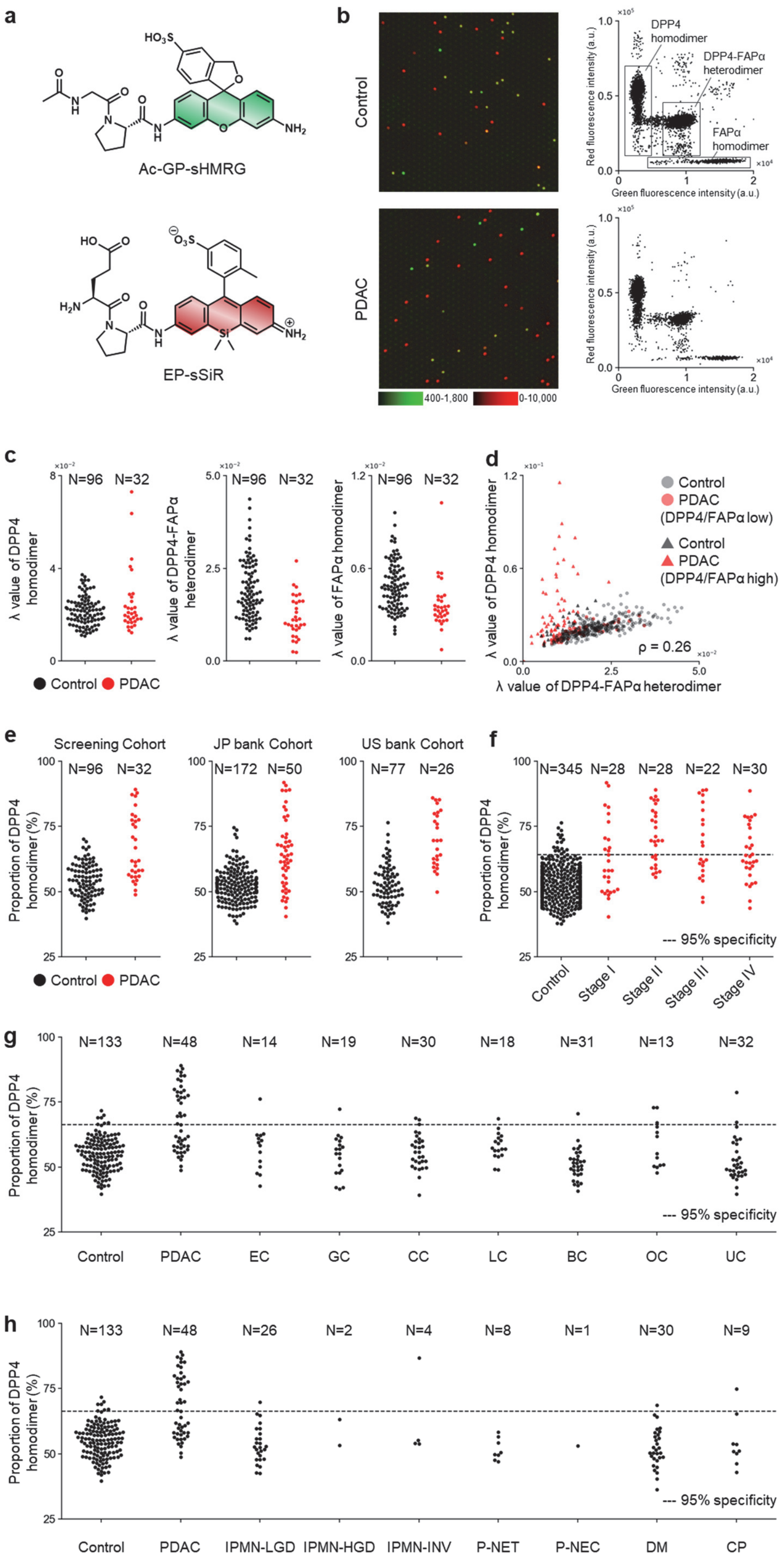

a
Ac-GP-sHMRG
EP-sSiR
b
Control
PDAC
400-1,800
0-10,000
DPP4 homodimer
DPP4-FAPα heterodimer
FAPα homodimer
Red fluorescence intensity (a.u.)
Green fluorescence intensity (a.u.)
c
N=96 N=32
λ value of DPP4 homodimer
λ value of DPP4-FAPα heterodimer
λ value of FAPα homodimer
Control PDAC
d
Control
PDAC (DPP4/FAPα low)
Control
PDAC (DPP4/FAPα high)
ρ = 0.26
λ value of DPP4 homodimer
λ value of DPP4-FAPα heterodimer
e
Screening Cohort
JP bank Cohort
US bank Cohort
N=96 N=32
N=172 N=50
N=77 N=26
Proportion of DPP4 homodimer (%)
Control PDAC
f
N=345 N=28 N=28 N=22 N=30
--- 95% specificity
Control
Stage I
Stage II
Stage III
Stage IV
g
N=133 N=48 N=14 N=19 N=30 N=18 N=31 N=13 N=32
Control PDAC EC GC CC LC BC OC UC
h
N=133 N=48 N=26 N=2 N=4 N=8 N=1 N=30 N=9
Control PDAC IPMN-LGD IPMN-HGD IPMN-INV P-NET P-NEC DM CP

**Figure S4.** Characterization and measurement of DPP4/FAPα activity. (a) Structures of Ac-GP-sHMRG and EP-sSiR. (b) Representative fluorescence images of the microdevice and corresponding scatter plots obtained after loading Ac-GP-sHMRG (30 μM), EP-sSiR (30 μM), IR-dye 800 (30 μM), and plasma samples (1:5,000 diluted) from control subjects or patients with PDAC in HEPES-Na buffer (100 mM, pH 7.4) containing $MgCl_2$ (1 mM), $CaCl_2$ (1 mM), DTT (100 μM), BSA (1% w/v), and Triton X-100 (150 μM), followed by incubation at 25°C for 4 h. (c) Distribution of lambda (signal numbers per all analyzed chambers) values for each cluster in the screening cohort. (d) Correlation between DPP4 homodimer and DPP4-FAPα heterodimer in patients with PDAC and control subjects from the combined training and validation sets (PDAC, N = 108, red; Control, N = 345, black). Triangles indicate samples with a high DPP4–DPP4/DPP4–FAPα ratio, defined as values over the 95th percentile of the control distribution. (e) Distribution of DPP4–DPP4/DPP4–FAPα ratio across sample cohorts, including Screening, JP bank, and US bank cohort (**Table S1**). (f) Distribution of DPP4–DPP4/DPP4–FAPα ratio in control subjects and PDAC patients in the validation set, stratified by disease stage. (g) Distribution of DPP4–DPP4/DPP4–FAPα ratio in control subjects and patients with cancers from various tissues. (h) Distribution of DPP4–DPP4/DPP4–FAPα ratio in control subjects and patients with PDAC or other diseases related to pancreatic dysfunction. Numbers above the plots indicate sample size. **Abbreviations:** EC, esophageal cancer; GC, gastric cancer; CC, colorectal cancer; LC, lung cancer; BC, breast cancer; OC, ovarian cancer; UC, uterine cancer; IPMN-LGD, intraductal papillary mucinous neoplasm with low-grade dysplasia; IPMN-HGD, IPMN with high-grade dysplasia; IPMN-INV, IPMN with an associated invasive carcinoma; P-NET, pancreatic neuroendocrine tumor; P-NEC, pancreatic neuroendocrine carcinoma.

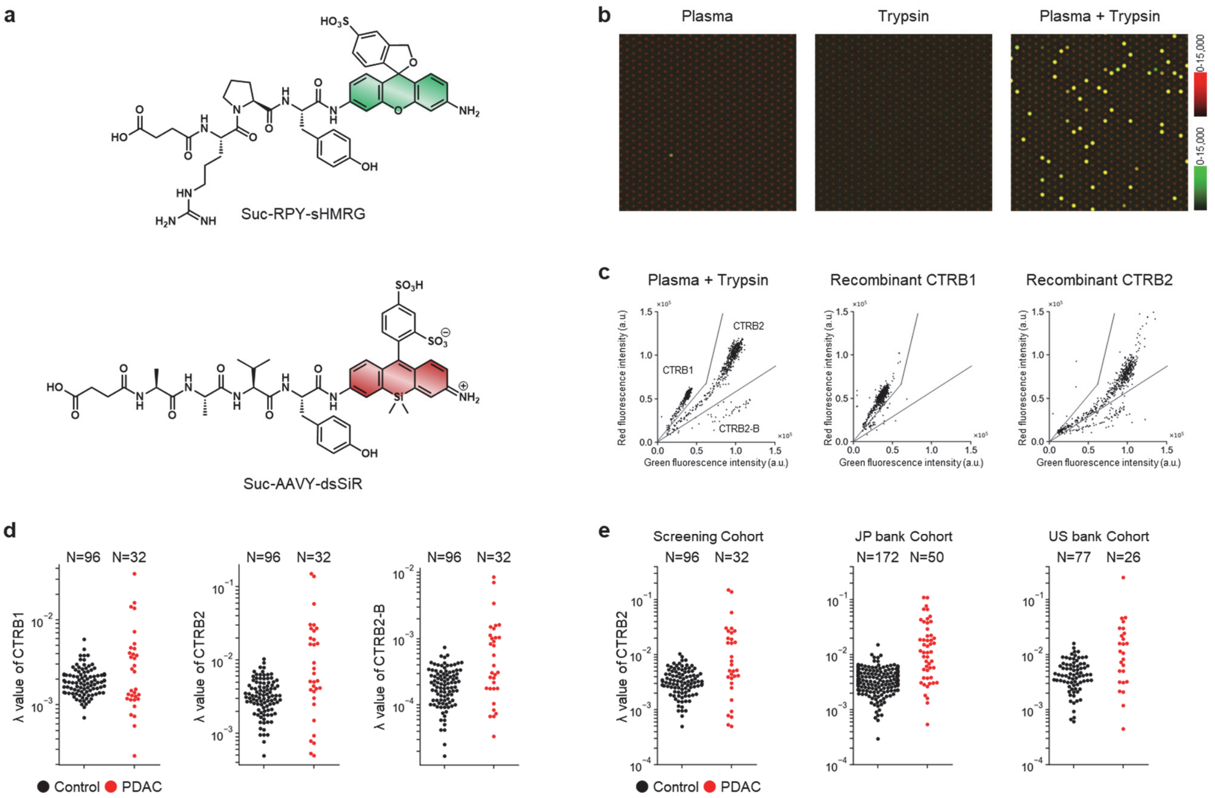


**Figure S5.** Characterization and measurement of chymotrypsin activity with trypsin activation. (a) Structures of Suc-RPY-sHMRG and Suc-AAVY-dsSiR. (b) Representative fluorescence images of the microdevice obtained after loading Suc-RPY-sHMRG (30 μM), Suc-AAVY-dsSiR (30 μM), IR-dye 800 (30 μM), and plasma sample (1:2,500 diluted), trypsin (25 μg/mL) or both into HEPES-Na buffer (100 mM, pH 7.4) containing $MgCl_2$ (1 mM), $CaCl_2$ (5 mM), NaCl (1 M), DTT (100 μM), and Triton X-100 (150 μM), followed by incubation at 25°C for 20 h. (c) Scatter plots of the fluorescence intensities of microdevice obtained after loading plasma sample and trypsin (1:2,500 diluted and 25 μg/mL), recombinant human CTRB1 (12.5 pg/mL), or recombinant human CTRB2 (12.5 pg/mL). Other conditions were the same as in (b). (d) Distribution of lambda values for each cluster in the screening cohort. (e) Distribution of lambda values for CTRB2 across sample cohorts, including Screening, JP bank, and US bank cohort (**Table S1**). Numbers above plots indicate sample size.

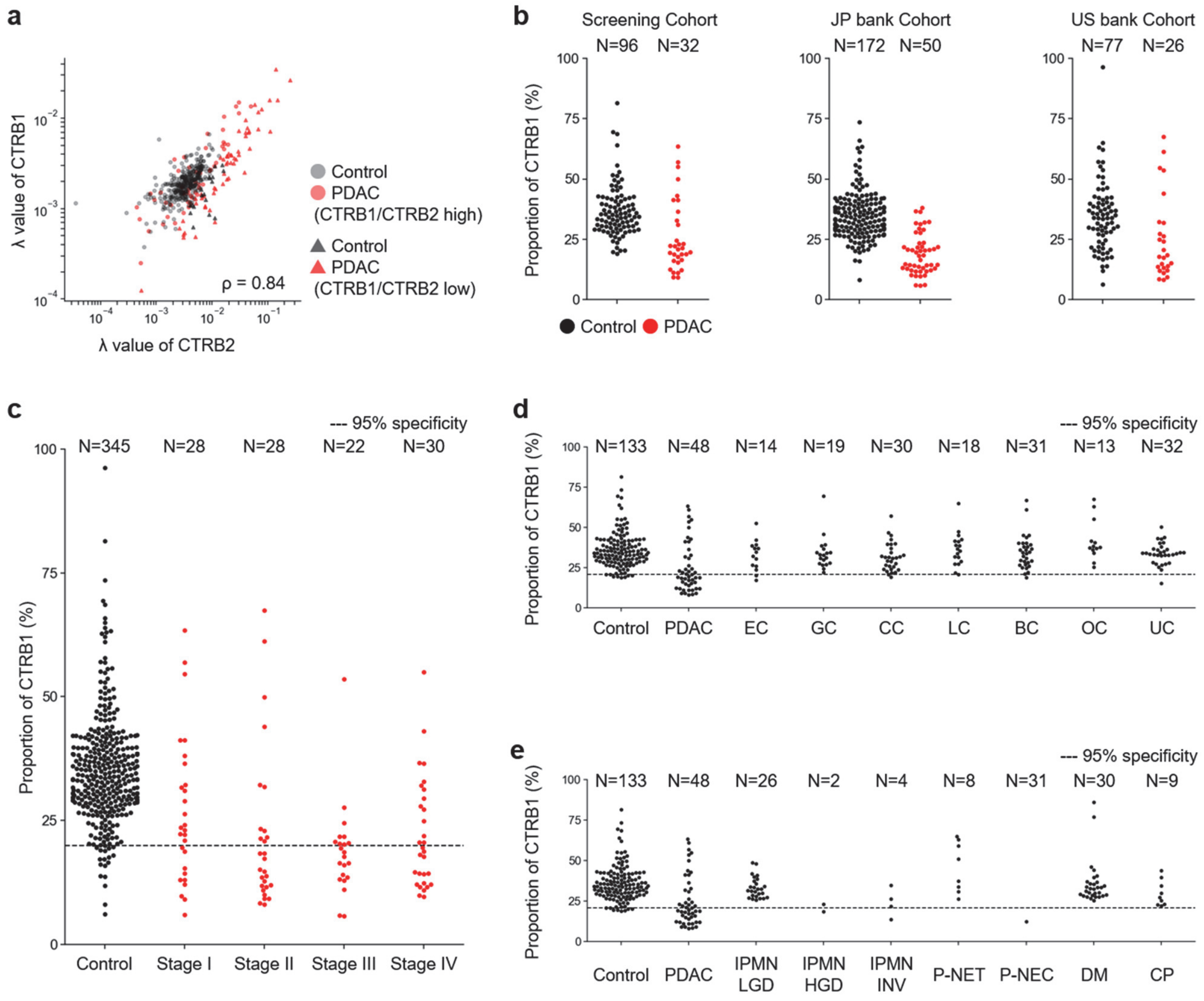


**Figure S6.** Profiles of CTRB1/CTRB2 ratio. (a) Correlation plot between CTRB1 and CTRB2 of patients with PDAC and control subjects from the combined training and validation sets (PDAC, N = 108, red; Control, N = 345, black). Triangles indicate samples with a low CTRB1/CTRB2 ratio, defined as values below the 5th percentile of the control distribution. (b) Distribution of CTRB1/CTRB2 ratio across sample cohorts, including Screening, JP bank, and US bank cohort (**Table S1**). (c) Distribution of CTRB1/CTRB2 ratio in control subjects and PDAC patients in the validation set, stratified by disease stage. (d) Distribution of CTRB1/CTRB2 ratio in control subjects and patients with PDAC or cancers from other tissues. (e) Distribution of CTRB1/CTRB2 ratio in control subjects and patients with PDAC or other pancreatic diseases. Numbers above plots indicate sample size. **Abbreviations:** EC, esophageal cancer; GC, gastric cancer; CC, colorectal cancer; LC, lung cancer; BC, breast cancer; OC, ovarian cancer; UC, uterine cancer; IPMN-LGD, intraductal papillary mucinous neoplasm with low-grade dysplasia; IPMN-HGD, IPMN with high-grade dysplasia; IPMN-INV, IPMN with an associated invasive carcinoma; P-NET, pancreatic neuroendocrine tumor; P-NEC, pancreatic neuroendocrine carcinoma.

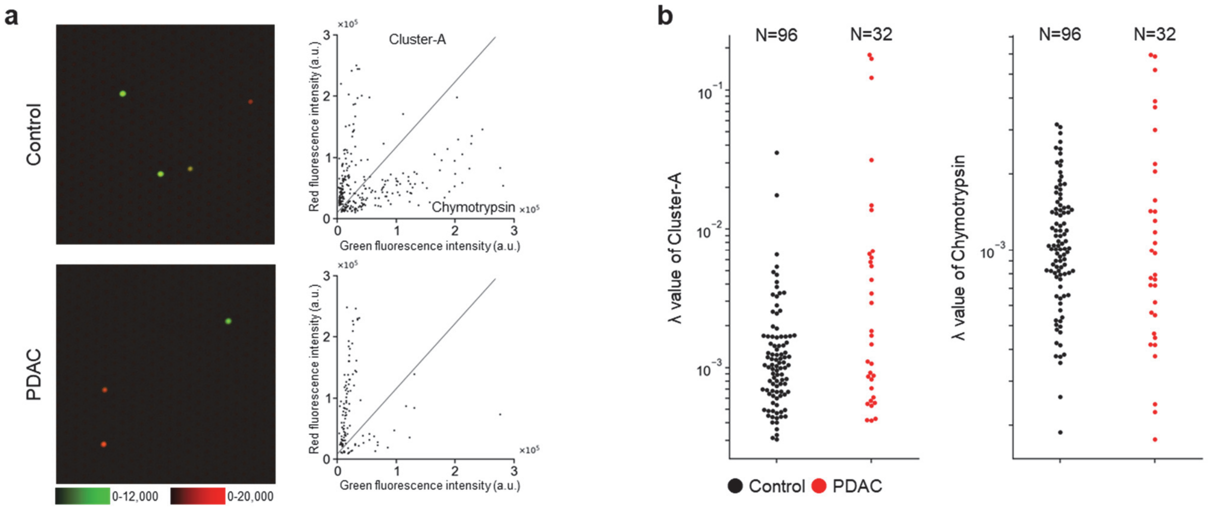


**Figure S7.** Characterization and measurement of chymotrypsin activity without trypsin activation. (a) Representative fluorescence images of the microdevice and corresponding scatter plots obtained after loading Suc-RPY-sHMRG (30 μM), Suc-AAVY-dsSiR (30 μM), IR-dye 800 (30 μM), and plasma samples (1:100 diluted) from control subjects or patients with PDAC into HEPES-Na buffer (125 mM, pH 7.4) containing $MgCl_2$ (1 mM), $CaCl_2$ (10 mM), NaCl (1.075 M), DTT (100 μM), CHAPS (0.05% w/v), and Triton X-100 (150 μM), followed by incubation at 25°C for 20 h. (b) Distribution of lambda values for each cluster in the screening cohort. Numbers above plots indicate sample sizes.

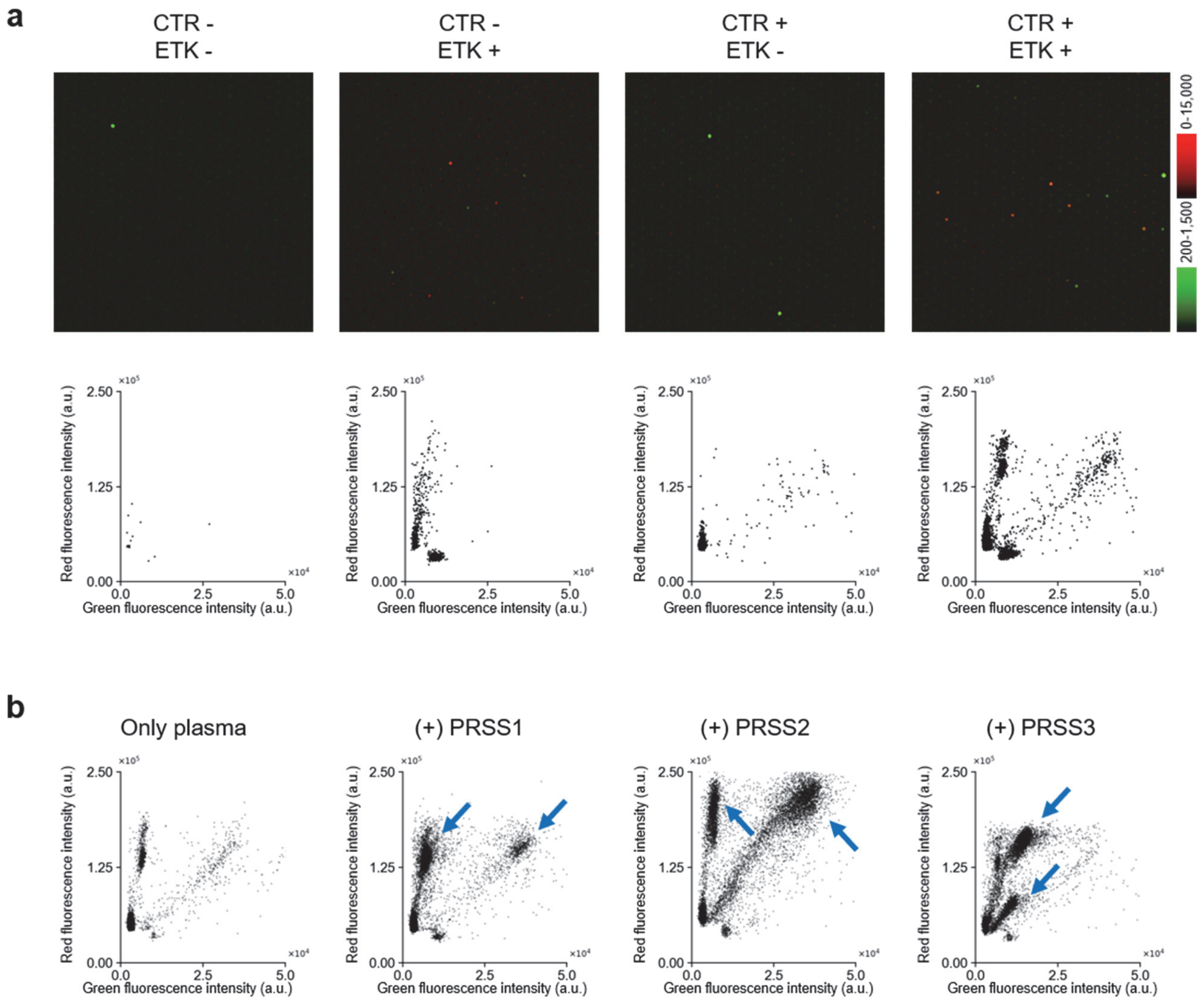


**Figure S8.** Optimization of activation conditions and characterization of the trypsin activity assay. (a) Representative fluorescence images of the microdevice and corresponding scatter plots obtained after loading Ac-RSPK-sHMRG (30 μM), Ac-IGGR-PABC-dsSiR (10 μM), IR-dye 800 (30 μM), and plasma samples (1:2,500 diluted) in the presence or absence of TLCK-treated chymotrypsin (labeled "CTR" in the figure, 16 μg/mL, pre-incubated for 3 h) and enterokinase (labeled "ETK" in the figure, 10 ng/mL) in HEPES-Na buffer (100 mM, pH 7.4) containing $MgCl_2$ (1 mM), $CaCl_2$ (5 mM), NaCl (50 mM), DTT (100 μM), bestatin (100 μM), and Triton X-100 (150 μM), followed by incubation at 25°C for 20 h. (b) Scatter plots obtained after loading Ac-RSPK-sHMRG (30 μM), Ac-IGGR-PABC-dsSiR (10 μM), IR-dye 800 (30 μM), and plasma samples (1:2,500 diluted, pre-treated with TLCK-treated chymotrypsin [16 μg/mL] for 3 h at 25°C) with or without recombinant PRSS1, PRSS2, or PRSS3 (0.1 ng/mL each) in HEPES-Na buffer (100 mM, pH 7.4) containing $MgCl_2$ (1 mM), $CaCl_2$ (5 mM), NaCl (50 mM), DTT (100 μM), bestatin (100 μM), enterokinase (10 ng/mL), and Triton X-100 (150 μM), followed by incubation at 25°C for 20 h. Arrows indicate clusters that appeared after the addition of recombinant protein to plasma samples.

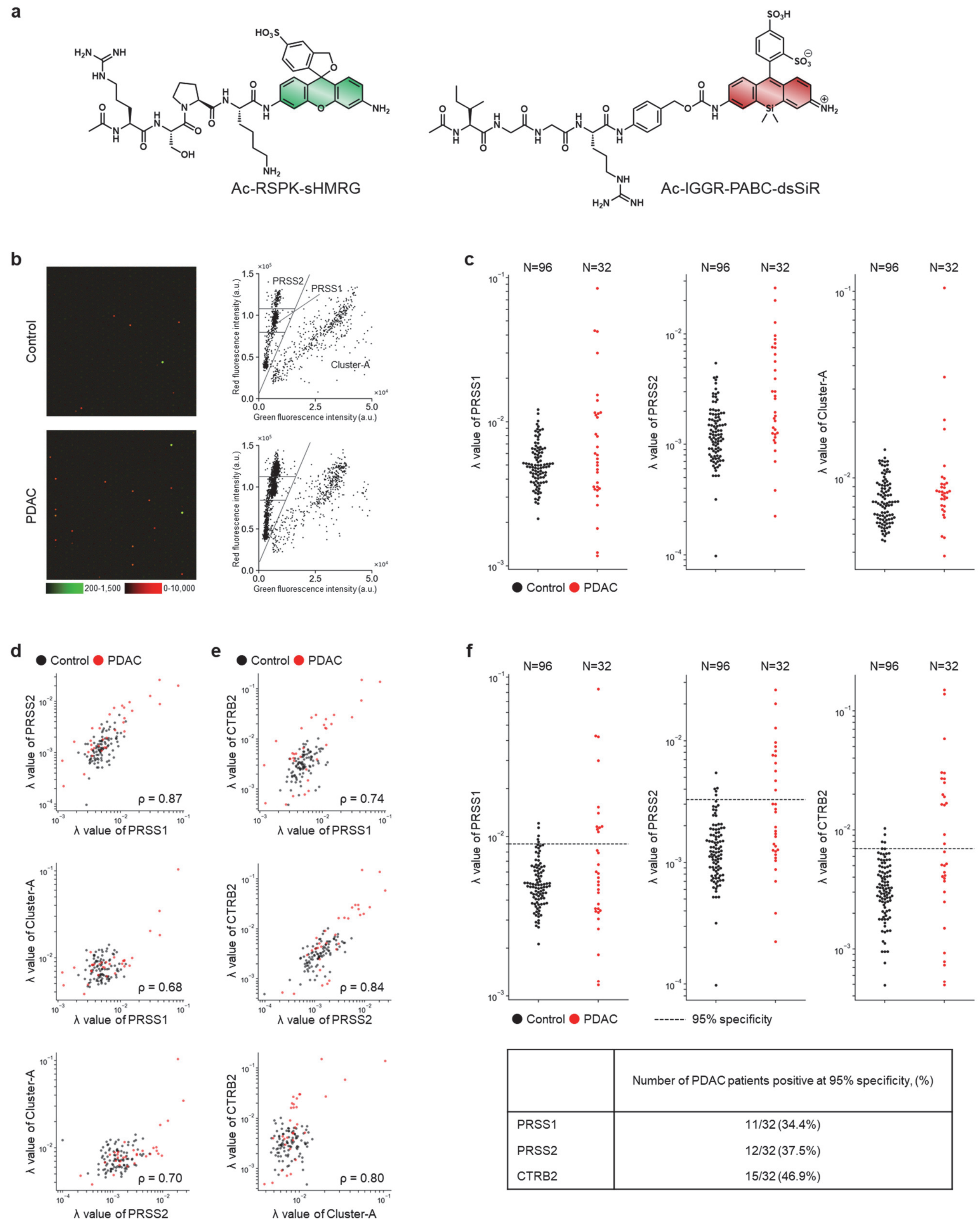


| | Number of PDAC patients positive at 95% specificity, (%) |
|---|---|
| PRSS1 | 11/32 (34.4%) |
| PRSS2 | 12/32 (37.5%) |
| CTRB2 | 15/32 (46.9%) |

**Figure S9.** Measurement of trypsin activity with enterokinase activation. (a) Structures of Ac-RSPK-sHMRG and Ac-IGGR-PABC-dsSiR. (b) Representative fluorescence images of the microdevice and corresponding scatter plots obtained after loading Ac-RSPK-sHMRG (30 μM), Ac-IGGR-PABC-dsSiR (10 μM), IR-dye 800 (30 μM), and plasma samples (1:2,500 diluted, pre-treated with TLCK-treated chymotrypsin (16 µg/mL) for 3 h at 25°C) from control subjects or patients with PDAC into HEPES-Na buffer (100 mM, pH 7.4) containing $MgCl_2$ (1 mM), $CaCl_2$ (5 mM), NaCl (50 mM), DTT (100 µM), bestatin (100 µM), enterokinase (10 ng/mL), and Triton X-100 (150 µM), followed by incubation at 25°C for 20 h. (c) Distribution of lambda values for each cluster in the screening cohort. (d) Correlations between PRSS1 and PRSS2, between PRSS1 and Cluster-A, and between PRSS2 and Cluster-A in the screening cohort (PDAC, N = 32, red; control, N = 96, black). ρ, Spearman's rank correlation coefficient calculated using PDAC samples only. (e) Correlation between each trypsin cluster and lambda value of CTRB2 in the screening cohort (PDAC, N = 32, red; control, N = 96, black). ρ, Spearman's rank correlation coefficient calculated using PDAC samples only. (f) Comparison of PDAC detection based on PRSS1, PRSS2, and CTRB2 lambda values at 95% specificity in the screening cohort. Dashed lines indicate cutoffs corresponding to 95% specificity among control subjects. Numbers above plots indicate sample sizes. The table shows the number and percentage of PDAC patients classified as positive using each marker.

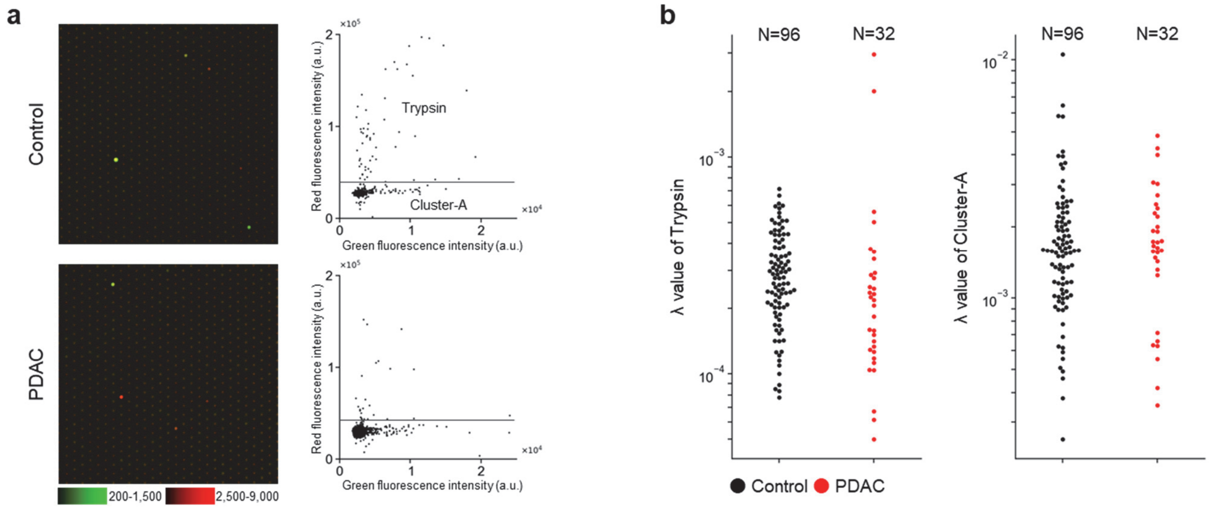


**Figure S10.** Characterization and measurement of trypsin activity without enterokinase activation. (a) Representative fluorescence images of the microdevice and corresponding scatter plots obtained after loading Ac-RSPK-sHMRG (30 μM), Ac-IGGR-PABC-dsSiR (10 μM), IR-dye 800 (30 μM), and plasma samples (1:250 diluted) from control subjects or patients with PDAC into HEPES-Na buffer (90 mM, pH 7.4) containing $MgCl_2$ (0.8 mM), $CaCl_2$ (4 mM), NaCl (70 mM), DTT (80 μM), CHAPS (0.02% w/v), and Triton X-100 (120 μM), followed by incubation at 25°C for 20 h. (b) Distribution of lambda values for each cluster in the screening cohort. Numbers above plots indicate sample sizes.

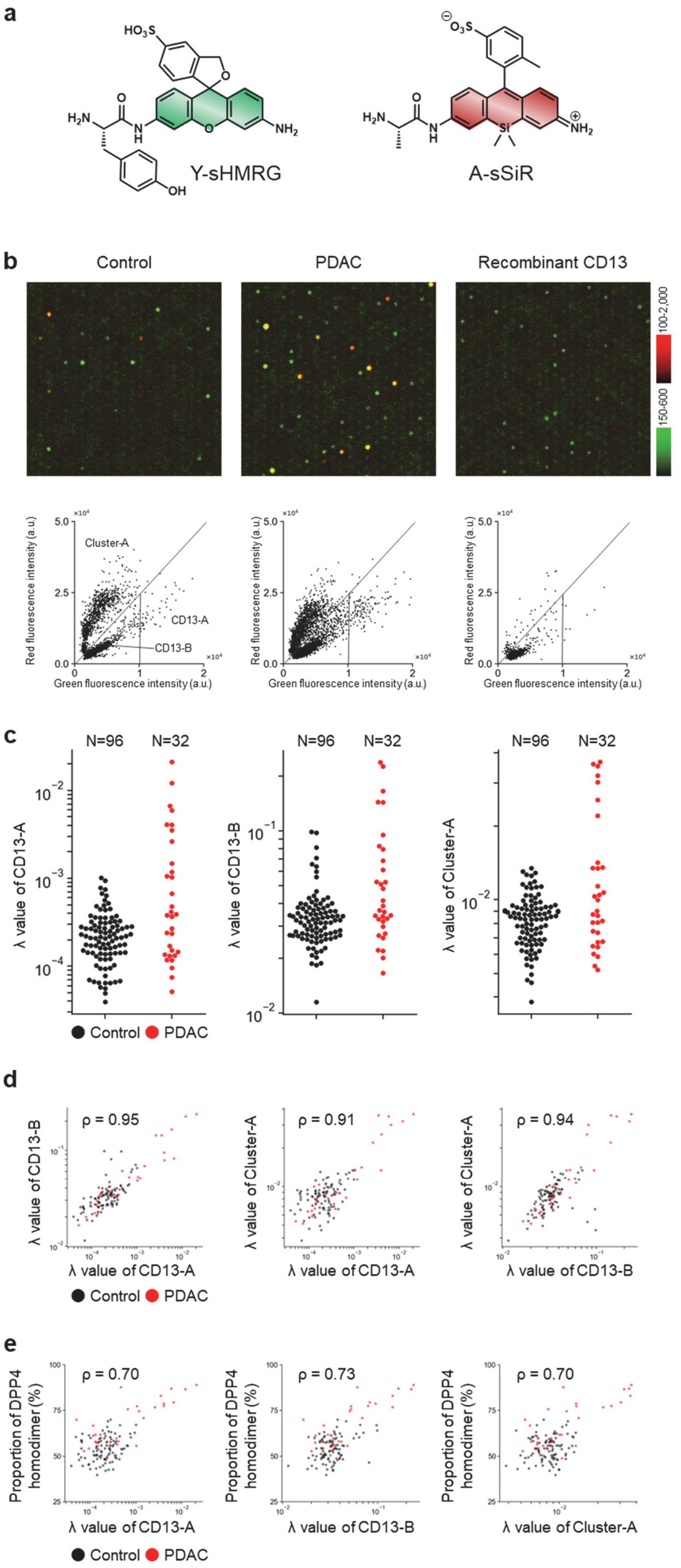
a
Y-sHMRG
A-sSiR
b
Control
PDAC
Recombinant CD13
100-2,000
150-600
Cluster-A
CD13-A
CD13-B
Red fluorescence intensity (a.u.)
Green fluorescence intensity (a.u.)
c
N=96
N=32
λ value of CD13-A
λ value of CD13-B
λ value of Cluster-A
Control
PDAC
d
ρ = 0.95
ρ = 0.91
ρ = 0.94
e
ρ = 0.70
ρ = 0.73
ρ = 0.70
Proportion of DPP4 homodimer (%)

**Figure S11.** Characterization and measurement of aminopeptidase activity. (a) Structures of Y-sHMRG and A-sSiR. (b) Representative fluorescence images of the microdevice and corresponding scatter plots obtained after loading Y-sHMRG (30 μM), A-sSiR (30 μM), IR-dye 800 (30 μM), and plasma samples (1:5,000 diluted) from control subjects or patients with PDAC, or recombinant CD13 (0.2 ng/mL) into HEPES-Na buffer (100 mM, pH 7.4) containing $MgCl_2$ (1 mM), $CaCl_2$ (1 mM), DTT (100 μM), and Triton X-100 (150 μM), followed by incubation at 25°C for 2 h. (c) Distribution of lambda values for each cluster in the screening cohort. Numbers above plots indicate sample sizes. (d) Correlations between CD13-A and CD13-B, between CD13-A and Cluster-A, and between CD13-B and Cluster-A in the screening cohort (PDAC, N = 32, red; control, N = 96, black). ρ, Spearman's rank correlation coefficient calculated using PDAC samples only. (e) Correlation between each aminopeptidase cluster and DPP4–DPP4/DPP4–FAPα ratio in the screening cohort (PDAC, N = 32, red; control, N = 96, black). ρ, Spearman's rank correlation coefficient calculated using PDAC samples only.

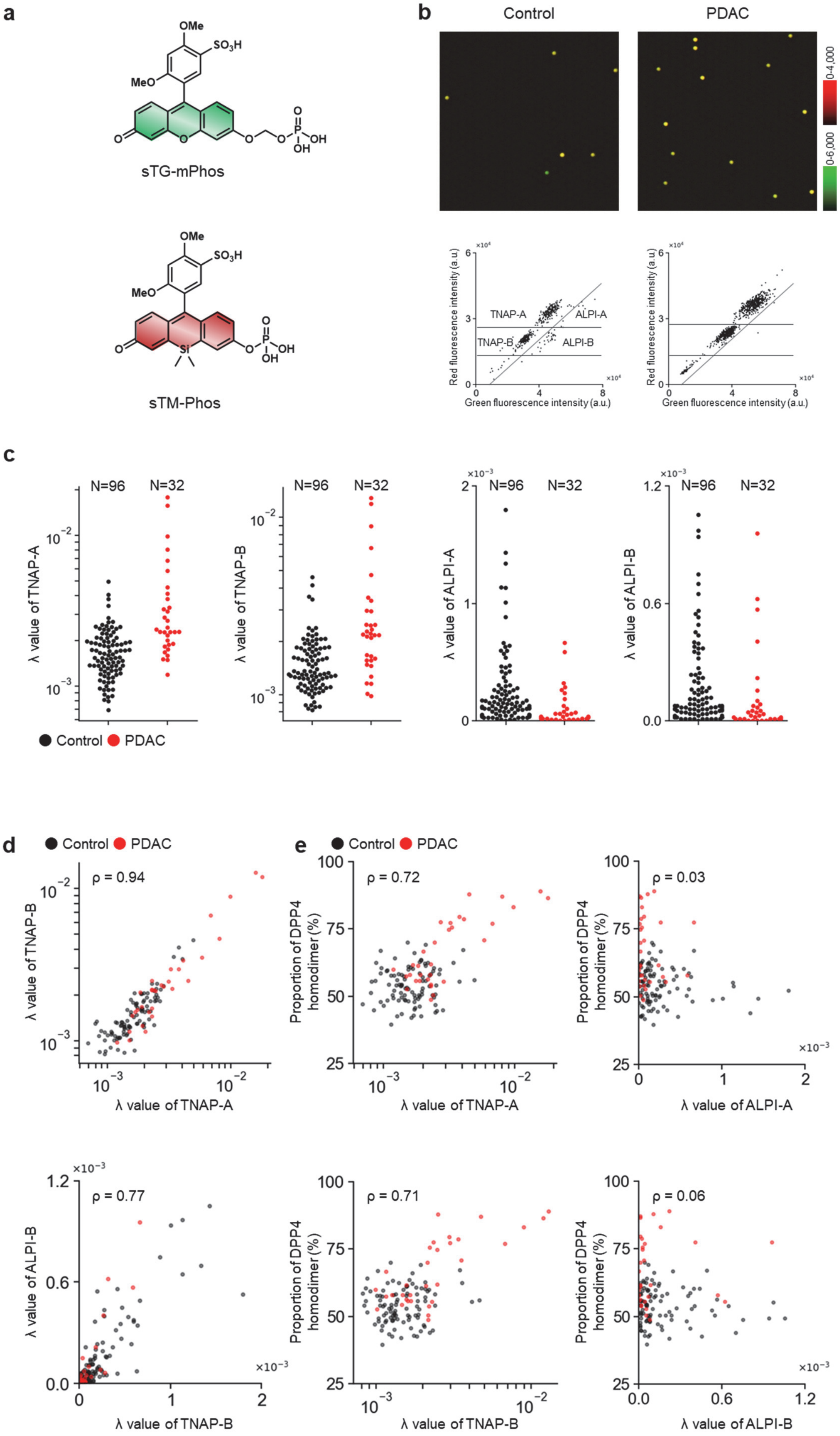
a
OMe
SO3H
MeO
sTG-mPhos
sTM-Phos
b
Control
PDAC
0-4,000
0-6,000
Red fluorescence intensity (a.u.)
Green fluorescence intensity (a.u.)
TNAP-A
ALPI-A
TNAP-B
ALPI-B
c
N=96
N=32
λ value of TNAP-A
λ value of TNAP-B
λ value of ALPI-A
λ value of ALPI-B
Control
PDAC
d
Control
PDAC
ρ = 0.94
ρ = 0.77
λ value of TNAP-A
λ value of TNAP-B
λ value of ALPI-B
e
ρ = 0.72
ρ = 0.03
ρ = 0.71
ρ = 0.06
Proportion of DPP4 homodimer (%)
λ value of ALPI-A
λ value of ALPI-B

**Figure S12.** Characterization and measurement of ALP activity. (a) Structures of sTG-mPhos and sTM-Phos. (b) Representative fluorescence images of the microdevice and corresponding scatter plots obtained after loading sTG-mPhos (25 μM), sTM-Phos (25 μM), IR-dye 800 (30 μM), and plasma samples (1:1,000 diluted) from control subjects or patients with PDAC into DEA-HCl buffer (1 M, pH 9.3) containing $MgCl_2$ (1 mM), $CaCl_2$ (1 mM), $ZnCl_2$ (20 μM), and Triton X-100 (150 μM), followed by incubation at 25°C for 50 min. Cluster identification was performed according to a previous report[9]. (c) Distribution of lambda values for each cluster in the screening cohort. Numbers above plots indicate sample sizes. (d) Correlations between TNAP-A and TNAP-B, and between ALPI-A and ALPI-B in the screening cohort (PDAC, N = 32, red; control, N = 96, black). ρ, Spearman's rank correlation coefficient calculated using PDAC samples only. (e) Correlations between each ALP cluster and DPP4–DPP4/DPP4–FAPα ratio in the screening cohort (PDAC, N = 32, red; control, N = 96, black). ρ, Spearman's rank correlation coefficient calculated using PDAC samples only.

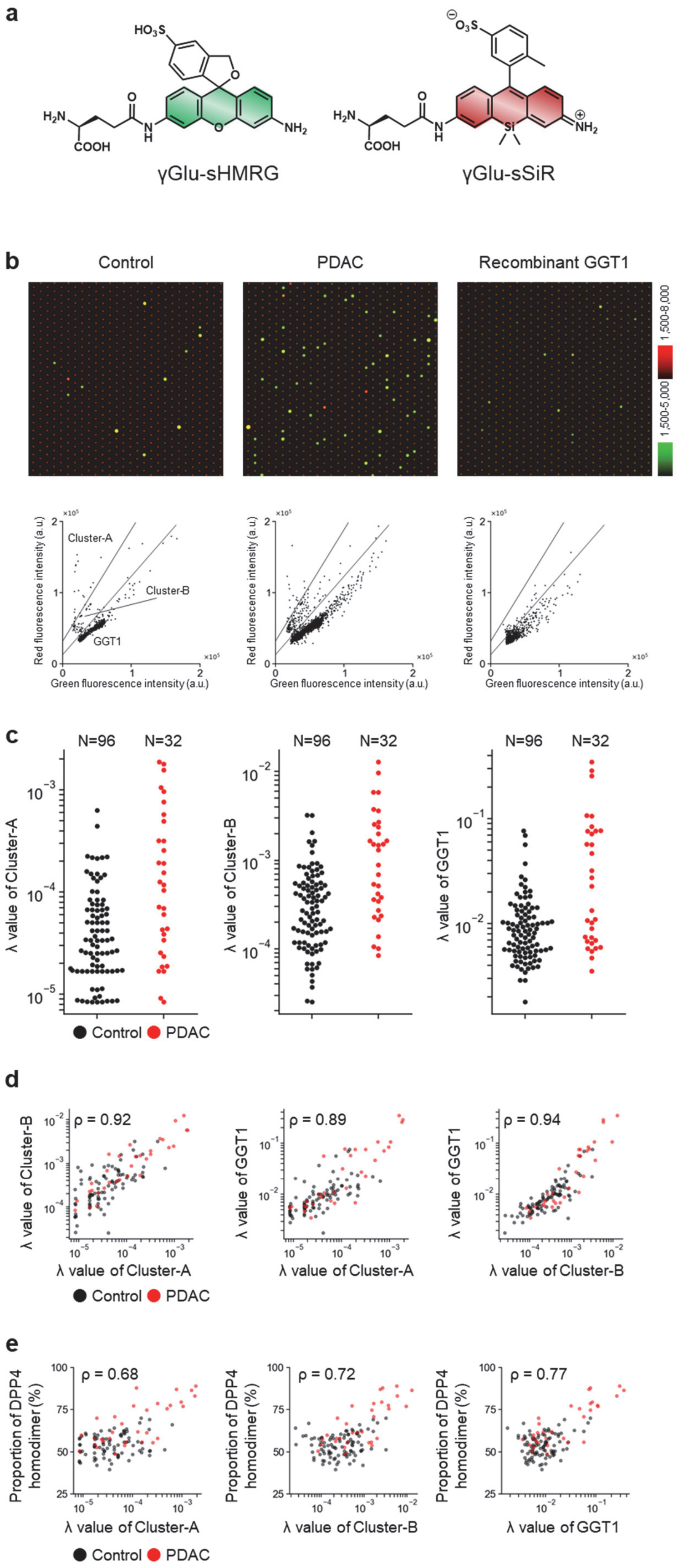
a
γGlu-sHMRG
γGlu-sSiR
b
Control
PDAC
Recombinant GGT1
1,500-8,000
1,500-5,000
Cluster-A
Cluster-B
GGT1
Red fluorescence intensity (a.u.)
Green fluorescence intensity (a.u.)
c
N=96
N=32
λ value of Cluster-A
λ value of Cluster-B
λ value of GGT1
Control
PDAC
d
ρ = 0.92
ρ = 0.89
ρ = 0.94
λ value of Cluster-B
λ value of GGT1
λ value of Cluster-A
λ value of Cluster-B
Control
PDAC
e
ρ = 0.68
ρ = 0.72
ρ = 0.77
Proportion of DPP4 homodimer (%)
λ value of Cluster-A
λ value of Cluster-B
λ value of GGT1
Control
PDAC

**Figure S13.** Characterization and measurement of GGT activity. (a) Structures of γGlu-sHMRG and γGlu-sSiR. (b) Representative fluorescence images of the microdevice and corresponding scatter plots obtained after loading γGlu-sHMRG (30 μM), γGlu-sSiR (30 μM), IR-dye 800 (30 μM), and plasma samples (1:500 diluted) from control subjects or patients with PDAC, or recombinant GGT1 (0.2 ng/mL) into MES-Na buffer (100 mM, pH 5.5) containing $MgCl_2$ (1 mM), $CaCl_2$ (1 mM), DTT (100 μM), and Triton X-100 (150 μM), followed by incubation at 25°C for 20 h. (c) Distribution of lambda values for each cluster in the screening cohort. Numbers above plots indicate sample sizes. (d) Correlations between Cluster-A and Cluster-B, between Cluster-A and GGT1, and between Cluster-B and GGT1 in the screening cohort (PDAC, N = 32, red; control, N = 96, black). ρ, Spearman's rank correlation coefficient calculated using PDAC samples only. (e) Correlations between each GGT cluster and DPP4–DPP4/DPP4–FAPα ratio in the screening cohort (PDAC, N = 32, red; control, N = 96, black). ρ, Spearman's rank correlation coefficient calculated using PDAC samples only.

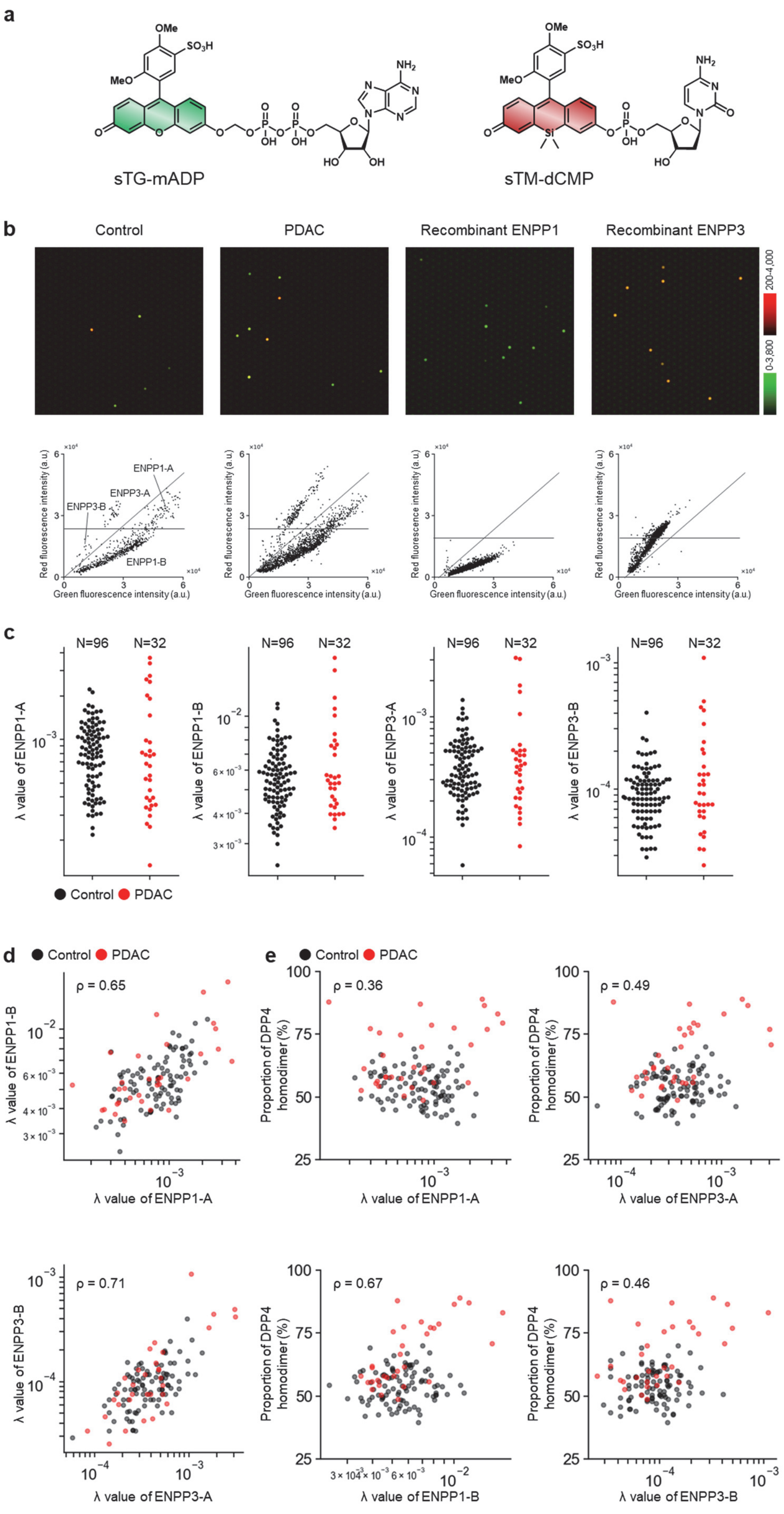

a
sTG-mADP
sTM-dCMP
b
Control
PDAC
Recombinant ENPP1
Recombinant ENPP3
ENPP1-A
ENPP3-A
ENPP3-B
ENPP1-B
Red fluorescence intensity (a.u.)
Green fluorescence intensity (a.u.)
c
N=96
N=32
λ value of ENPP1-A
λ value of ENPP1-B
λ value of ENPP3-A
λ value of ENPP3-B
Control
PDAC
d
e
ρ = 0.65
ρ = 0.36
ρ = 0.49
ρ = 0.71
ρ = 0.67
ρ = 0.46
Proportion of DPP4 homodimer (%)

**Figure S14.** Characterization and measurement of ENPP1/3 activity. (a) Structures of sTG-mADP and sTM-dCMP. (b) Representative fluorescence images of the microdevice and corresponding scatter plots obtained after loading sTG-mADP (30 μM), sTM-dCMP (30 μM), IR-dye 800 (60 μM), and plasma samples (1:200 diluted) from control subjects or patients with PDAC, or recombinant ENPP1 (2.5 ng/mL) or ENPP3 (250 ng/mL) into Tris-HCl buffer (100 mM, pH 9.0) containing HEPES-Na (25 mM), $MgCl_2$ (1 mM), $CaCl_2$ (1 mM), NaCl (75 mM), $ZnCl_2$ (250 μM), recombinant ALP (80 ng/mL), CHAPS (0.05% w/v), and Triton X-100 (150 μM), followed by incubation at 25°C for 1 h. (c) Distribution of lambda values for each cluster in the screening cohort. Numbers above plots indicate sample sizes. (d) Correlations between ENPP1-A cluster and ENPP1-B cluster, and between ENPP3-A cluster and ENPP3-B cluster in the screening cohort (PDAC, N = 32, red; control, N = 96, black). ρ, Spearman's rank correlation coefficient calculated using PDAC samples only. (e) Correlations between each ENPP1/3 cluster and DPP4–DPP4/DPP4–FAPα ratio in the screening cohort (PDAC, N = 32, red; control, N = 96, black). ρ, Spearman's rank correlation coefficient calculated using PDAC samples only.

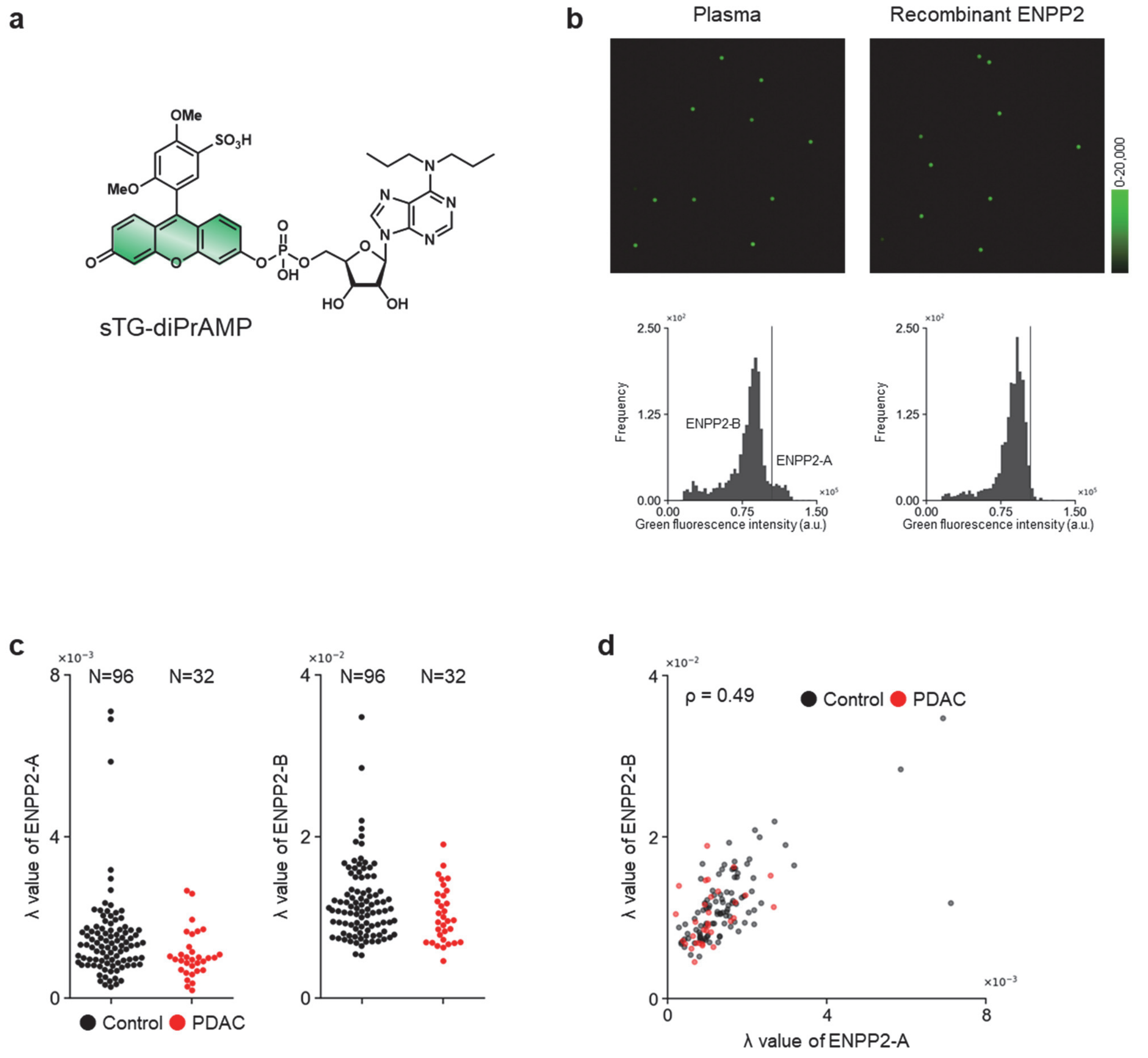


**Figure S15.** Characterization and measurement of ENPP2 activity. (a) Structure of sTG-diPrAMP. (b) Representative fluorescence images of the microdevice and corresponding histograms obtained after loading sTG-diPrAMP (30 μM), sCy5-SE (10 μM), and plasma samples (1:5,000 diluted) from control subjects, or recombinant ENPP2 (0.1 ng/mL) into Tris-HCl buffer (100 mM, pH 9.0) containing $MgCl_2$ (5 mM), NaCl (150 mM), BSA (1% w/v), and Triton X-100 (150 μM), followed by incubation at 25°C for 2 h. (c) Distribution of lambda values for each cluster in the screening cohort. Numbers above plots indicate sample sizes. (d) Correlation between ENPP2-A cluster and ENPP2-B cluster in the screening cohort (PDAC, N = 32, red; control, N = 96, black). ρ, Spearman's rank correlation coefficient calculated using PDAC samples only.

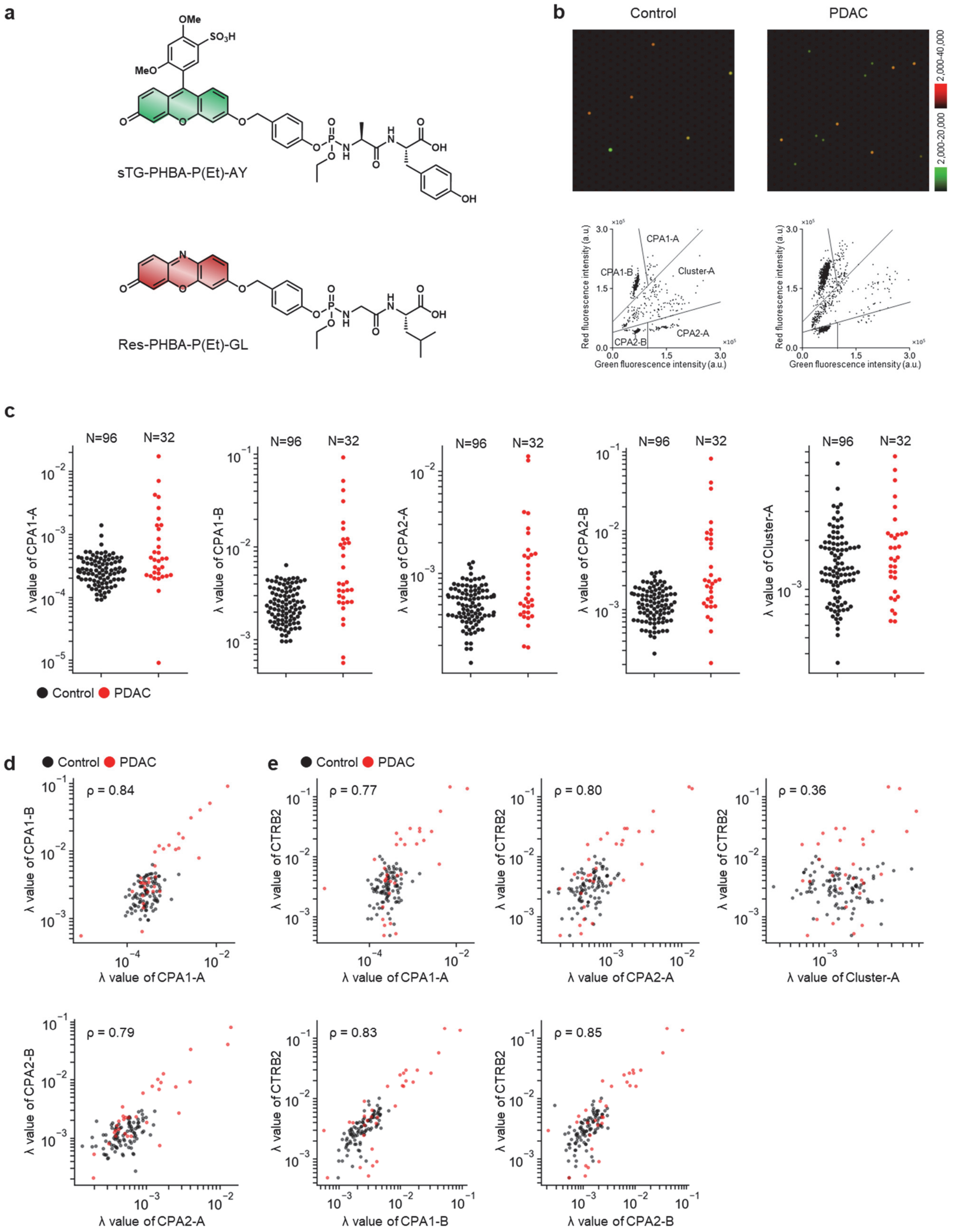


**Figure S16.** Characterization and measurement of CPA activity. (a) Structures of sTG-PHBA-P(Et)-AY and

Res-PHBA-P(Et)-GL. (b) Representative fluorescence images of the microdevice and corresponding scatter plots obtained after loading sTG-PHBA-P(Et)-AY (10 μM), Res-PHBA-P(Et)-GL (10 μM), IR-dye 800 (30 μM), and plasma samples (1:2,500 diluted) from control subjects or patients with PDAC into Tris-HCl buffer (100 mM, pH 8.5) containing $CaCl_2$ (5 mM), $ZnCl_2$ (10 μM), DTT (100 μM), Trypsin (1.67 μg/mL), and Triton X-100 (150 μM), followed by incubation at 25°C for 24 h. Cluster identification was performed according to a previous report[8]. (c) Distribution of lambda values of CPA1 and CPA2 in the screening cohort. Numbers above plots indicate sample size. (d) Correlations between CPA1-A cluster and CPA1-B cluster, and between CPA2-A cluster and CPA2-B cluster in the screening cohort (PDAC, N = 32, red; control, N = 96, black). ρ, Spearman's rank correlation coefficient calculated using PDAC samples only. (e) Correlations between each CPA cluster and lambda value of CTRB2 in the screening cohort (PDAC, N = 32, red; control, N = 96, black). ρ, Spearman's rank correlation coefficient calculated using PDAC samples only.

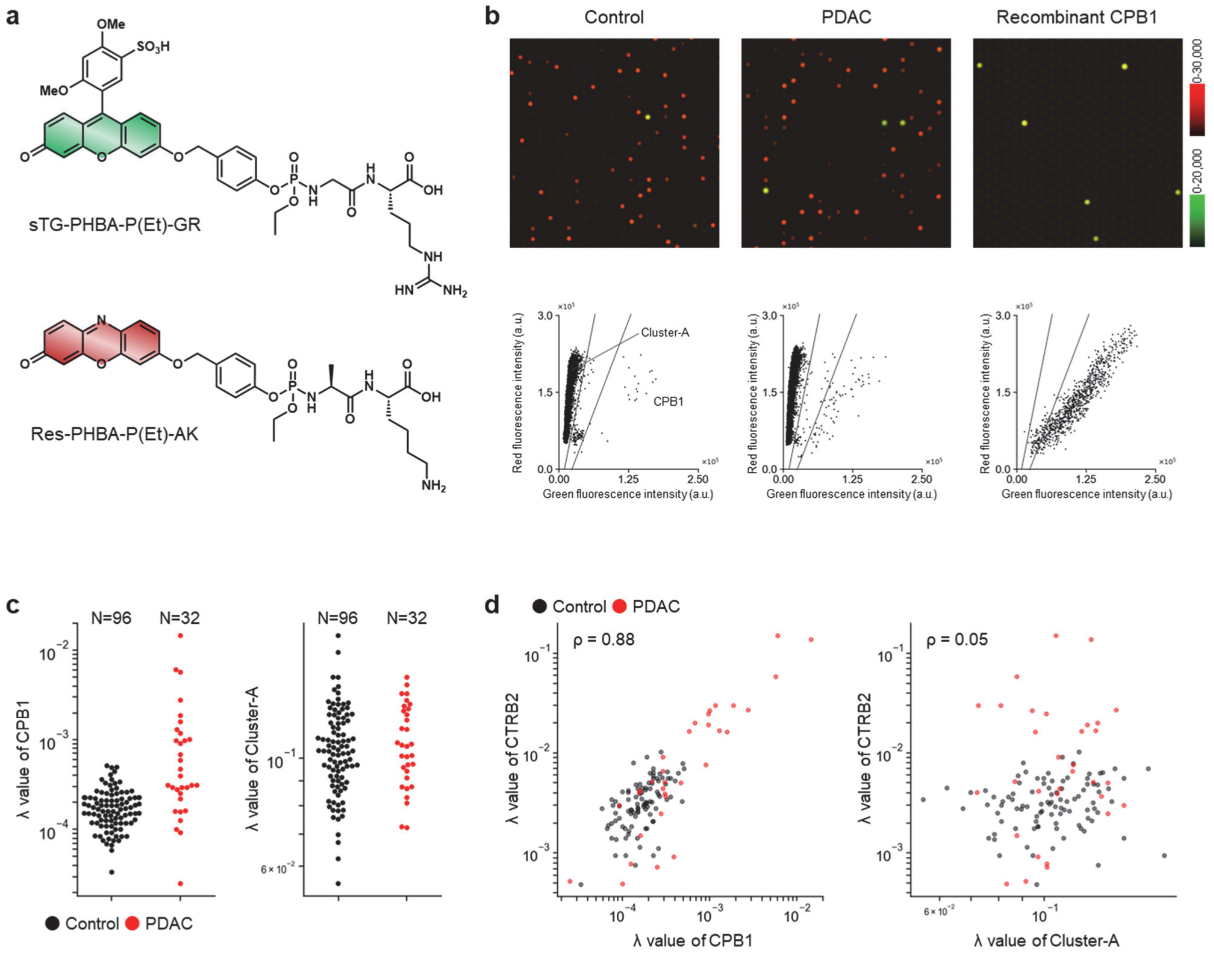


**Figure S17.** Characterization and measurement of CPB activity. (a) Structures of sTG-PHBA-P(Et)-GR and Res-PHBA-P(Et)-AK. (b) Representative fluorescence images of the microdevice and corresponding scatter plots obtained after loading sTG-PHBA-P(Et)-GR (10 μM), Res-PHBA-P(Et)-AK (10 μM), IR-dye 800 (30 μM), and plasma samples (1:20,000 diluted) from control subjects or patients with PDAC, or recombinant CPB1 (0.1 ng/mL) into HEPES-Na buffer (100 mM, pH 7.4) containing NaCl (100 mM), $ZnCl_2$ (100 μM), DTT (100 μM), Trypsin (0.21 μg/mL), and Triton X-100 (150 μM), followed by incubation at 25°C for 20 h. (c) Distribution of lambda values for each cluster in the screening cohort. Numbers above plots indicate sample sizes. (d) Correlations between each CPB cluster and lambda value of CTRB2 in the screening cohort (PDAC, N = 32, red; control, N = 96, black). ρ, Spearman's rank correlation coefficient calculated using PDAC samples only.

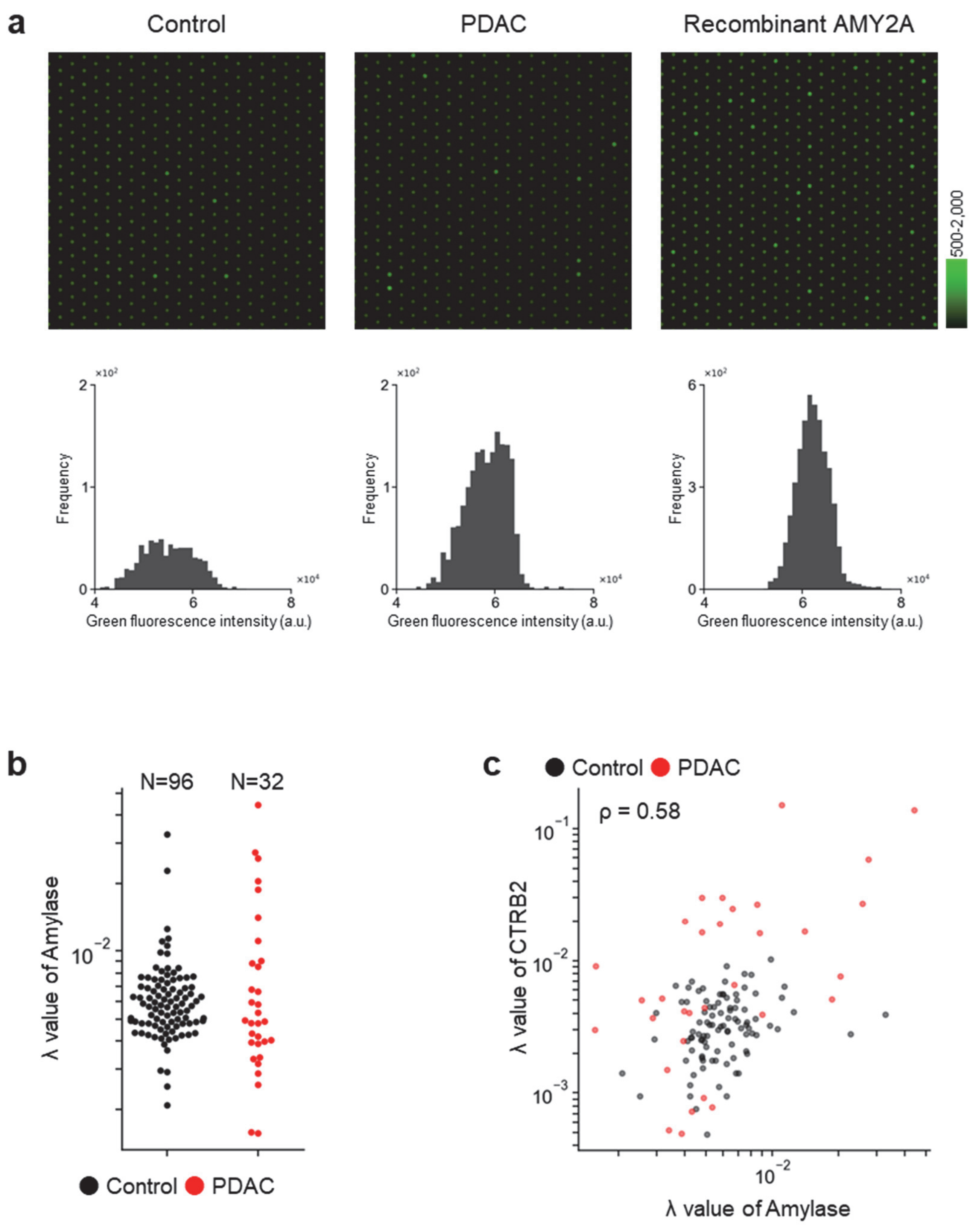


**Figure S18.** Characterization and measurement of amylase activity. (a) Representative fluorescence images of the microdevice and corresponding histograms obtained after loading sTG-Amy (1 μM), IR-dye 800 (30 μM), and plasma samples (1:1,000 diluted) from control subjects or patients with PDAC, or recombinant AMY2A (1 ng/mL) into HEPES-Na buffer (100 mM, pH 7.4) containing $MgCl_2$ (1 mM), $CaCl_2$ (5 mM), NaCl (100 mM), and Triton X-100 (150 μM), followed by incubation at 25°C for 24 h. (b) Distribution of lambda value of amylase in the screening cohort. Numbers above plots indicate sample sizes. (c) Correlation between lambda value of amylase and lambda value of CTRB2 in the screening cohort (PDAC, N = 32, red; control, N = 96, black). ρ, Spearman's rank correlation coefficient calculated using PDAC samples only.

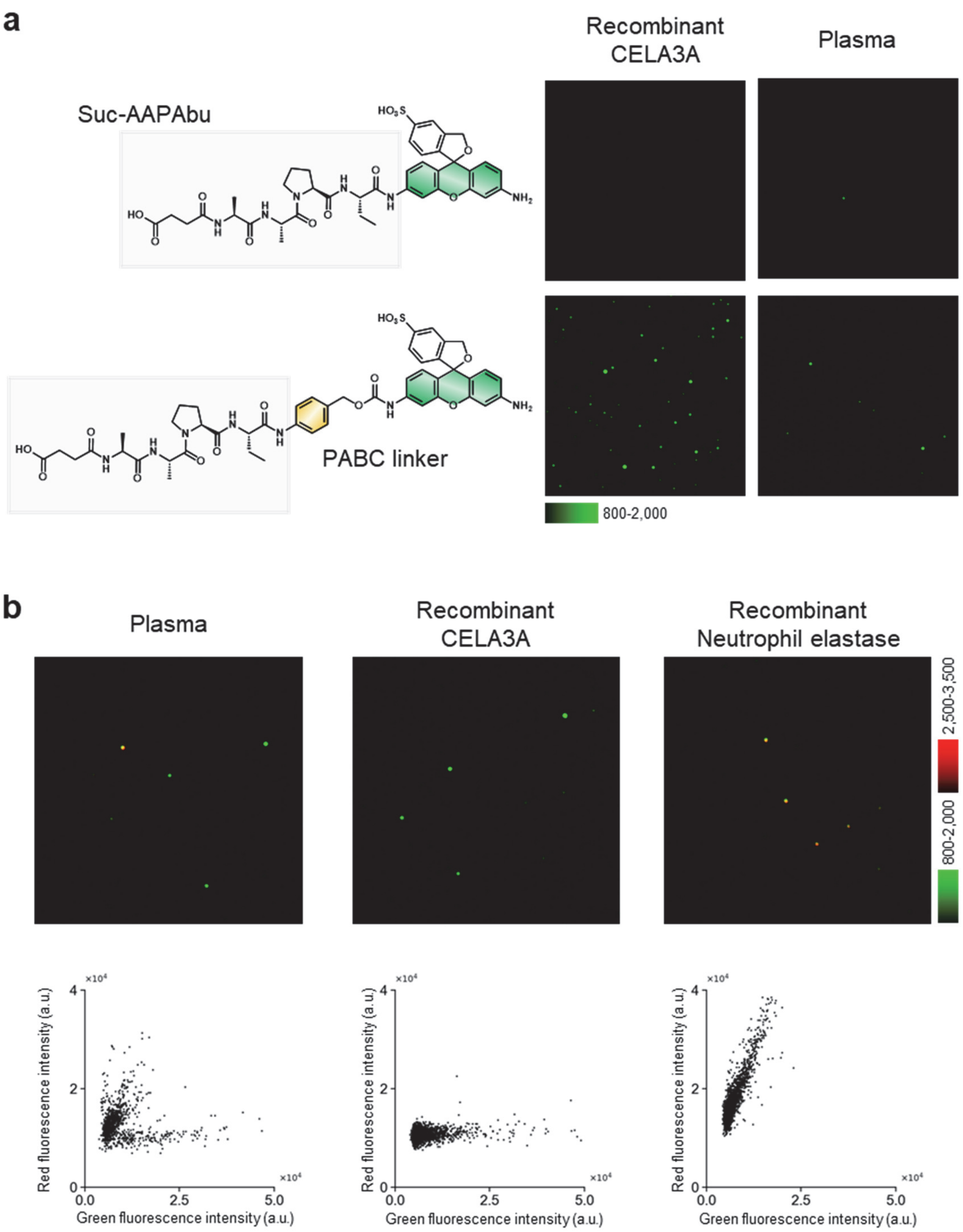


**Figure S19.** Characterization of elastase. (a) Structures of Suc-AAPAbu-sHMRG and Suc-AAPAbu-PABC-sHMRG, and representative fluorescence images of the microdevice obtained after loading Suc-AAPAbu-sHMRG (10 μM) or Suc-AAPAbu-PABC-sHMRG (10 μM), IR-dye 800 (30 μM), and plasma samples (1:100 diluted), or recombinant CELA3A (50 ng/mL) into HEPES-Na buffer (125 mM, pH 7.4) containing $MgCl_2$ (1 mM), $CaCl_2$ (10 mM), NaCl (75 mM), DTT (100 μM), CHAPS (0.05% w/v), and Triton X-100 (150 μM), followed by incubation at 25°C for 24 h. (b) Representative fluorescence images of the microdevice and corresponding scatter plots obtained after loading Suc-AAPAbu-PABC-sHMRG (10 μM), Suc-AAPV-PABC-dsSiR (15 μM), IR-dye 800 (30 μM), and plasma sample (1:100 diluted), or recombinant CELA3A (10 ng/mL) or purified native neutrophil elastase (10 pg/mL) into HEPES-Na buffer (125 mM, pH 7.4) containing $MgCl_2$ (1 mM), $CaCl_2$ (10 mM), NaCl (75 mM), DTT (100 μM), CHAPS (0.05% w/v), and Triton X-100 (150 μM), followed by incubation at 25°C for 24 h.

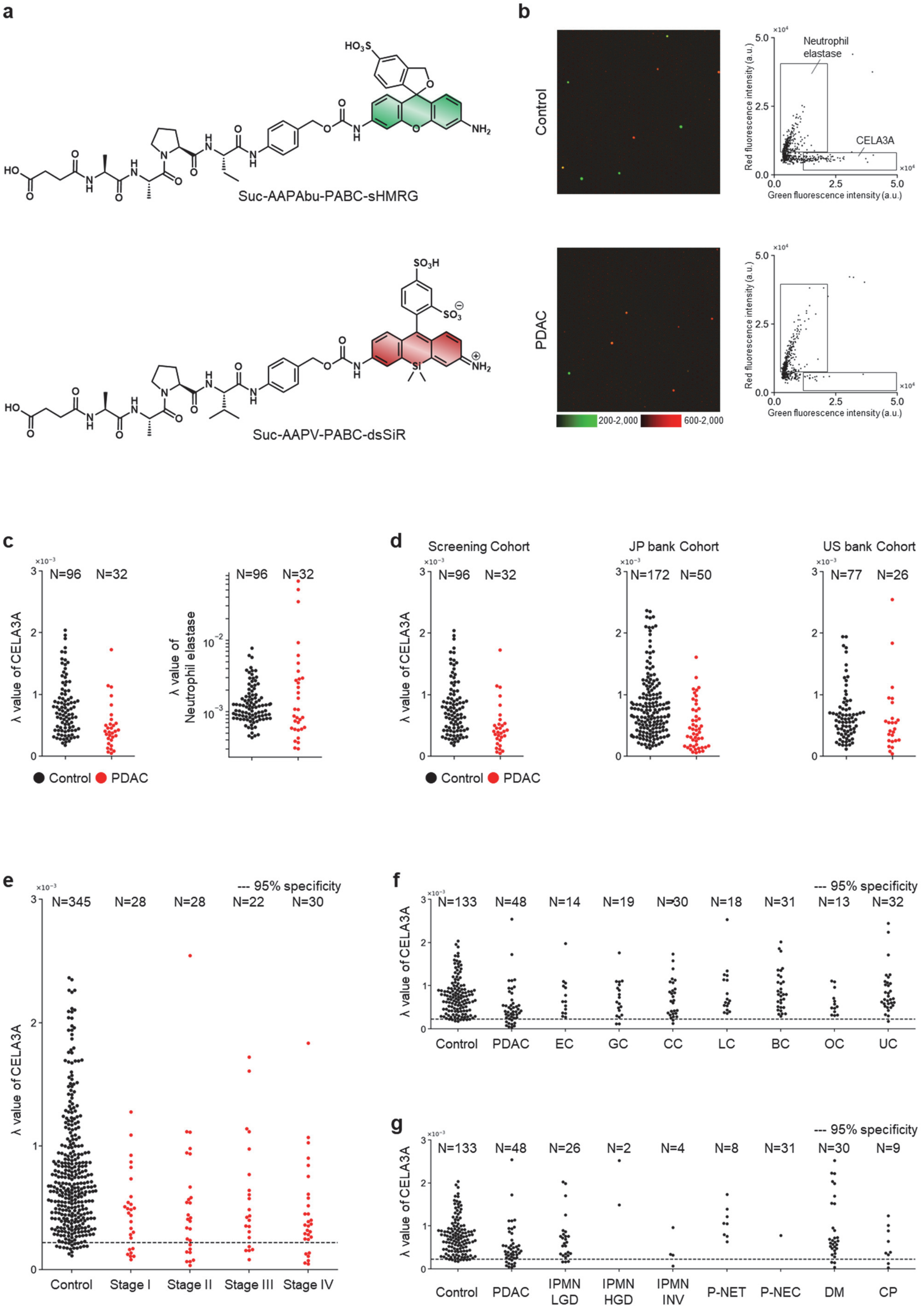
a
Suc-AAPAbu-PABC-sHMRG
Suc-AAPV-PABC-dsSiR
b
Control
PDAC
Neutrophil elastase
CELA3A
Red fluorescence intensity (a.u.)
Green fluorescence intensity (a.u.)
200-2,000
600-2,000
c
N=96 N=32
λ value of CELA3A
λ value of Neutrophil elastase
Control PDAC
d
Screening Cohort
JP bank Cohort
US bank Cohort
N=96 N=32
N=172 N=50
N=77 N=26
e
--- 95% specificity
N=345 N=28 N=28 N=22 N=30
Control Stage I Stage II Stage III Stage IV
f
N=133 N=48 N=14 N=19 N=30 N=18 N=31 N=13 N=32
Control PDAC EC GC CC LC BC OC UC
g
N=133 N=48 N=26 N=2 N=4 N=8 N=31 N=30 N=9
Control PDAC IPMN LGD IPMN HGD IPMN INV P-NET P-NEC DM CP

**Figure S20.** Measurement of elastase activity. (a) Structures of Suc-AAPAbu-PABC-sHMRG and Suc-AAPV-PABC-dsSiR. (b) Representative fluorescence images of the microdevice and corresponding scatter plots obtained after loading Suc-AAPAbu-PABC-sHMRG (10 μM), Suc-AAPV-PABC-dsSiR (15 μM), IR-dye 800 (30 μM), and plasma samples (1:100 diluted) from control subjects or patients with PDAC into HEPES-Na buffer (125 mM, pH 7.4) containing $MgCl_2$ (1 mM), $CaCl_2$ (10 mM), NaCl (75 mM), DTT (100 μM), CHAPS (0.05% w/v), and Triton X-100 (150 μM), followed by incubation at 25°C for 24 h. (c) Distribution of lambda values for each cluster in the screening cohort. (d) Distribution of lambda value of CELA3A across sample cohorts, including Screening, JP bank, and US bank cohort (**Table S1**). (e) Distribution of lambda values of CELA3A in control subjects and PDAC patients in the validation set, stratified by disease stage. (f) Distribution of lambda values of CELA3A in control subjects and patients with cancers from various tissues. (g) Distribution of lambda values of CELA3A in control subjects and patients with PDAC or other diseases related to pancreatic dysfunction. Numbers above the plots indicate sample sizes. **Abbreviations:** EC, esophageal cancer; GC, gastric cancer; CC, colorectal cancer; LC, lung cancer; BC, breast cancer; OC, ovarian cancer; UC, uterine cancer; IPMN-LGD, intraductal papillary mucinous neoplasm with low-grade dysplasia; IPMN-HGD, IPMN with high-grade dysplasia; IPMN-INV, IPMN with an associated invasive carcinoma; P-NET, pancreatic neuroendocrine tumor; P-NEC, pancreatic neuroendocrine carcinoma.

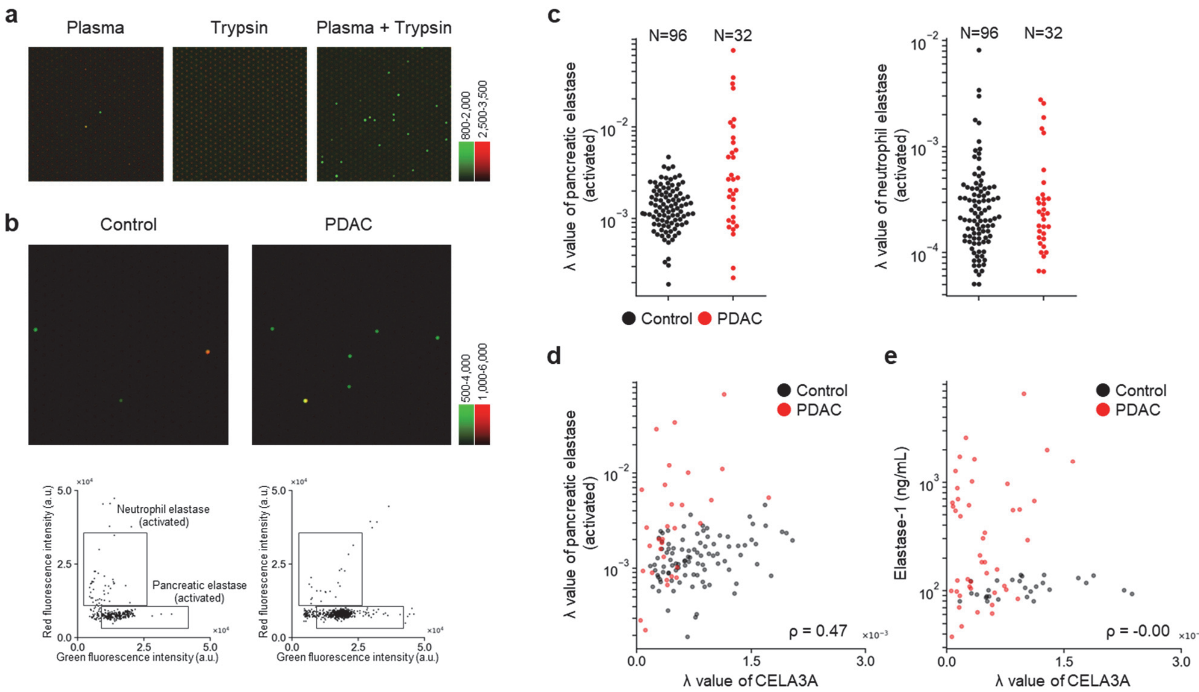


**Figure S21.** Characterization and measurement of elastase activity with trypsin activation. (a) Representative fluorescence images of the microdevice obtained after loading Suc-AAPAbu-PABC-sHMRG (10 μM), Suc-AAPV-PABC-dsSiR (15 μM), IR-dye 800 (30 μM), and plasma sample (1:100 diluted), trypsin (4 μg/mL) or both into HEPES-Na buffer (125 mM, pH 7.4) containing $MgCl_2$ (1 mM), $CaCl_2$ (10 mM), NaCl (75 mM), DTT (100 μM), and Triton X-100 (150 μM), followed by incubation at 25°C for 24 h. (b) Representative fluorescence images of the microdevice and corresponding scatter plots obtained after loading Suc-AAPAbu-PABC-sHMRG (10 μM), Suc-AAPV-PABC-dsSiR (15 μM), IR-dye 800 (30 μM), and plasma samples (1:1,000 diluted) from control subjects or patients with PDAC into the same buffer as in (a), followed by incubation at 25°C for 24 h. (c) Distribution of lambda values for each cluster in the screening cohort. Numbers above plots indicate sample sizes. (d) Correlation between lambda value of activated pancreatic elastase and lambda value of CELA3A in the screening cohort (PDAC, N = 32, red; control, N = 96, black). ρ, Spearman's rank correlation coefficient calculated using PDAC samples only. (e) Correlation between lambda value of CELA3A and clinical elastase-1 values in samples with available elastase-1 data (PDAC, N = 46, red; control, N = 29, black). ρ, Spearman's rank correlation coefficient calculated across all samples included in this analysis.

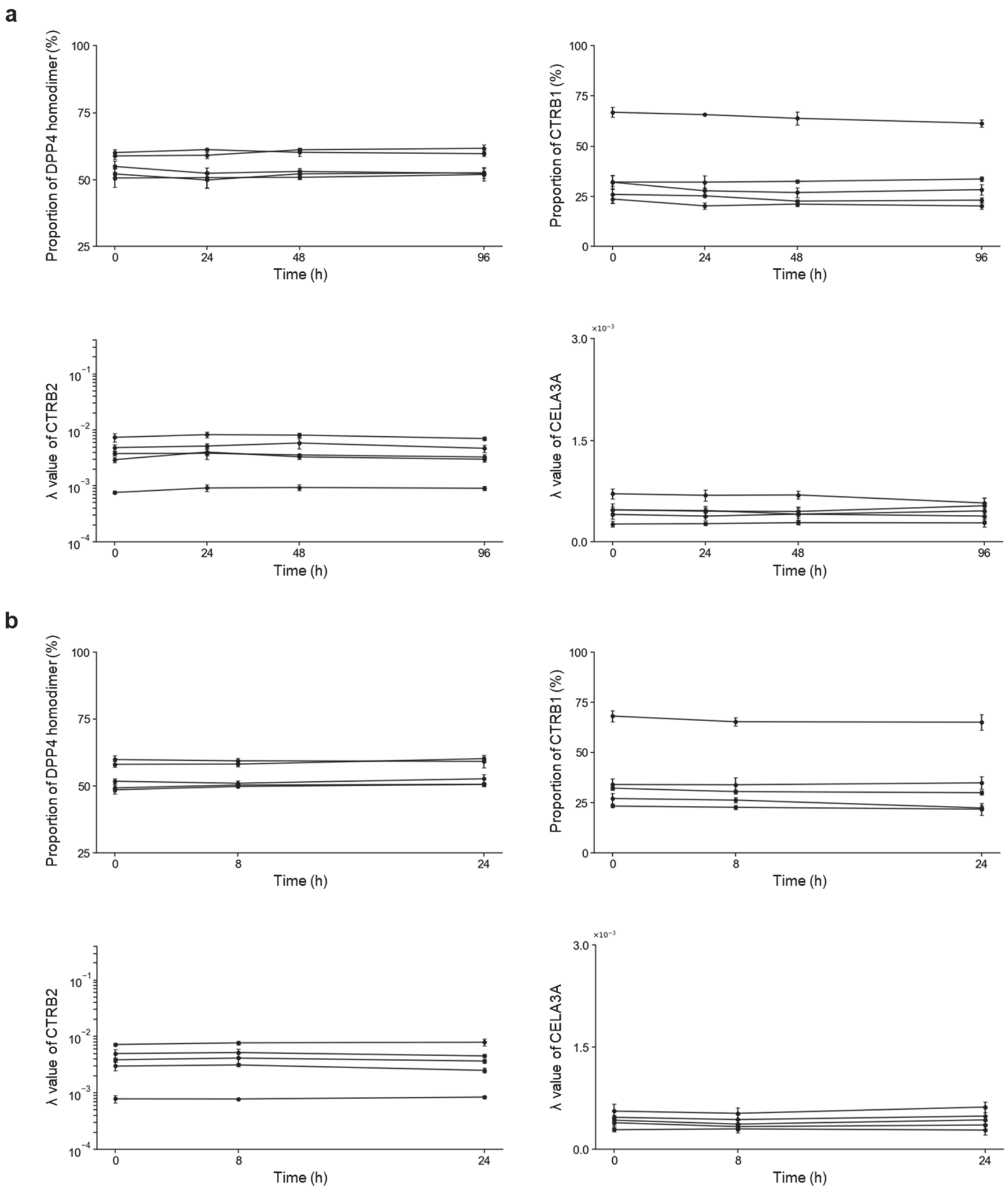


**Figure S22.** Stability of selected biomarkers in whole blood and plasma at 25°C. (a) Stability in whole blood. DPP4/FAPα, CTRB1/CTRB2, CTRB2, and CELA3A were measured in plasma samples obtained from whole blood stored at 25°C for 0, 24, 48, and 96 h. Measurements were conducted in five independent donors (N = 5), and each data point represents the mean ± s.d. of technical triplicates (n = 3). (b) Stability in plasma. DPP4/FAPα, CTRB1/CTRB2, CTRB2, and CELA3A were measured in plasma samples obtained from plasma stored at 25°C for 0, 8, and 24 h. Measurements were conducted in five independent donors (N = 5), and each data point and the error bars represent the mean ± s.d. of technical triplicates (n = 3).

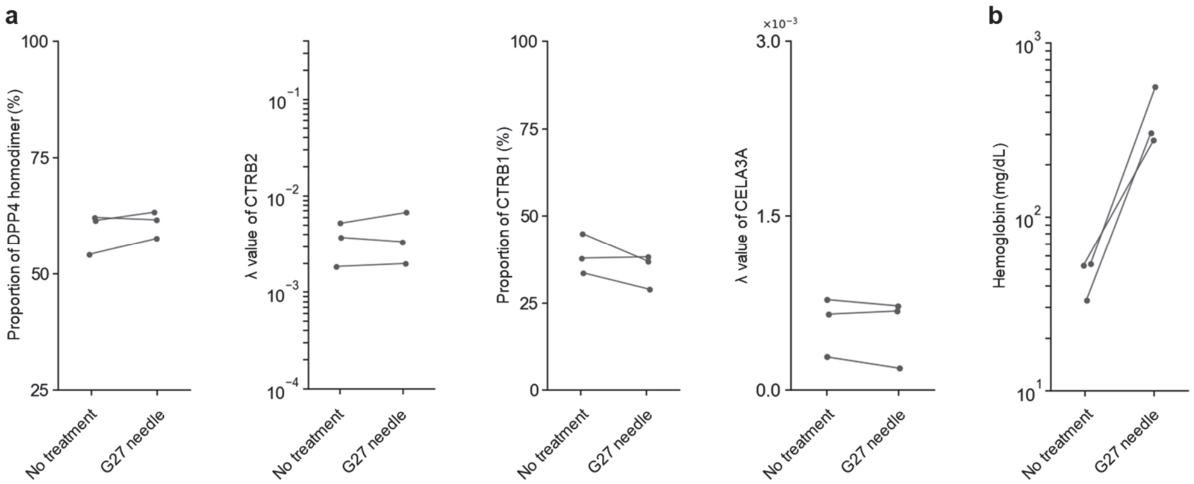


**Figure S23.** Effect of hemolysis. (a) DPP4/FAPα, CTRB2, CTRB1/CTRB2 and CELA3A marker values were measured in plasma samples obtained from whole blood without treatment or after induction of hemolysis by passage through a 27G needle. (b) Hemoglobin concentrations were measured in the same paired samples to confirm induction of hemolysis. Measurements were conducted in three independent donors (N = 3). *P* values were calculated using two-sided paired t-tests. No statistically significant changes were observed in the enzymatic marker values after 27G needle treatment (DPP4/FAPα, $P$ = 0.306; CTRB2, $P$ = 0.523; CTRB1/CTRB2, $P$ = 0.232; CELA3A, $P$ = 0.371), whereas hemoglobin levels increased in all three paired samples after treatment ($P$ = 0.061).

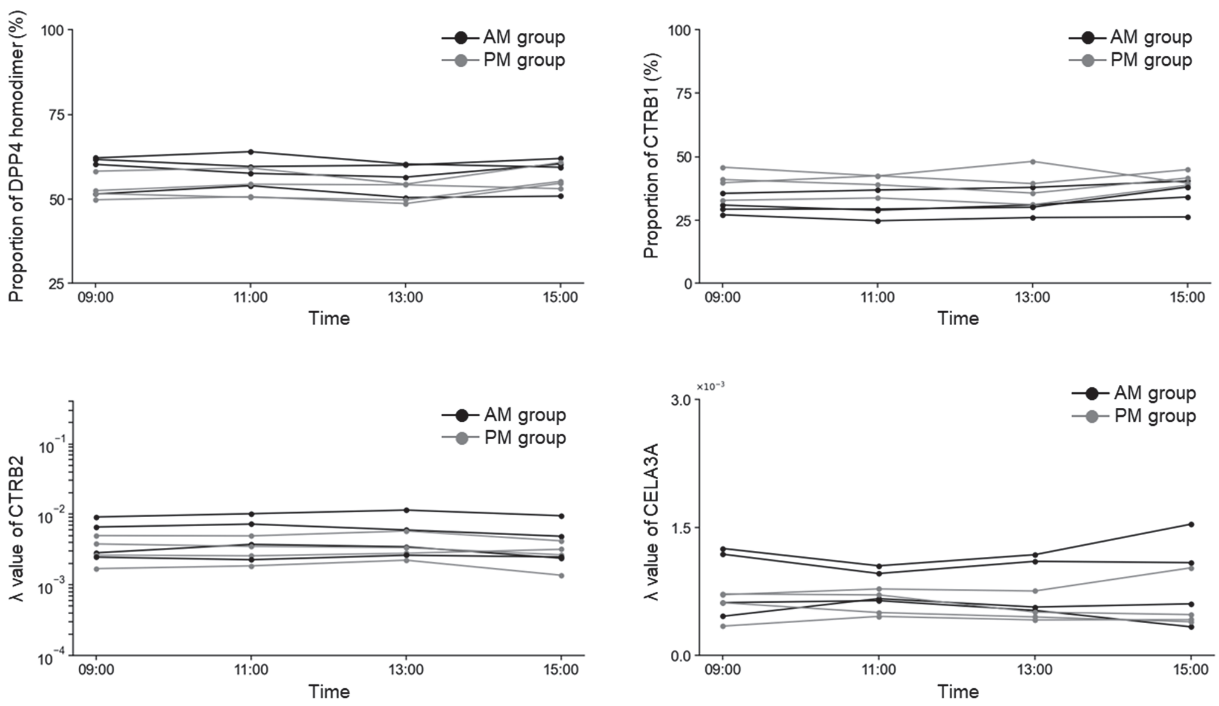


**Figure S24.** Diurnal variation. DPP4/FAPα, CTRB1/CTRB2, CTRB2, and CELA3A were measured in plasma samples collected at 09:00, 11:00, 13:00, and 15:00. Measurements were conducted in eight independent donors (N = 8), separated into AM and PM groups.

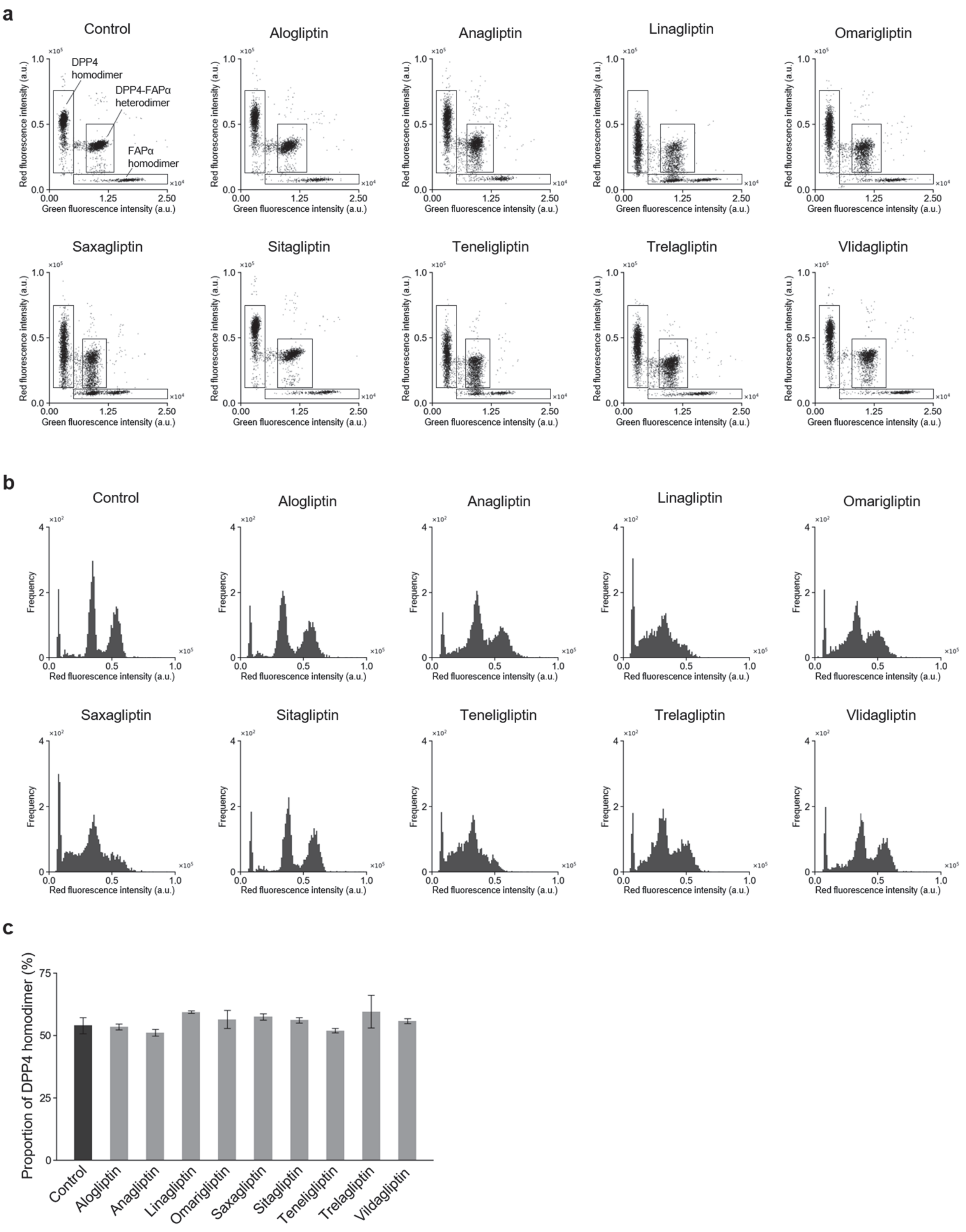


**Figure S25.** Effect of DPP4 inhibitors on DPP4/FAPα measurements. The DPP4 inhibitor panel covered all nine inhibitors approved in Japan. (a) Representative scatter plots of green and red fluorescence intensities obtained from plasma samples treated with DPP4 inhibitors. Undiluted plasma was supplemented with each DPP4 inhibitor at its reported maximum plasma concentration ($C_{max}$; **Table S9**) and incubated at 25°C for 30 min before dilution. Control plasma was supplemented with water instead of inhibitor. After pretreatment, samples were diluted 1:5,000 and analyzed with Ac-GP-sHMRG (30 μM), EP-sSiR (30 μM), and IR-dye 800 (30 μM) in HEPES-Na buffer (100 mM, pH 7.4) containing $MgCl_2$ (1 mM), $CaCl_2$ (1 mM), DTT (100 μM), BSA (1% w/v), and Triton X-100 (150 μM), followed by incubation at 25°C for 4 h. Rectangular gates indicate populations corresponding to FAPα homodimers, DPP4 homodimers, and DPP4–FAPα heterodimers. (b) Histograms of red fluorescence intensity for the corresponding measurements. (c) DPP4/FAPα values for plasma samples treated with each DPP4 inhibitor. Measurements were performed in technical triplicate, and error bars represent s.d.

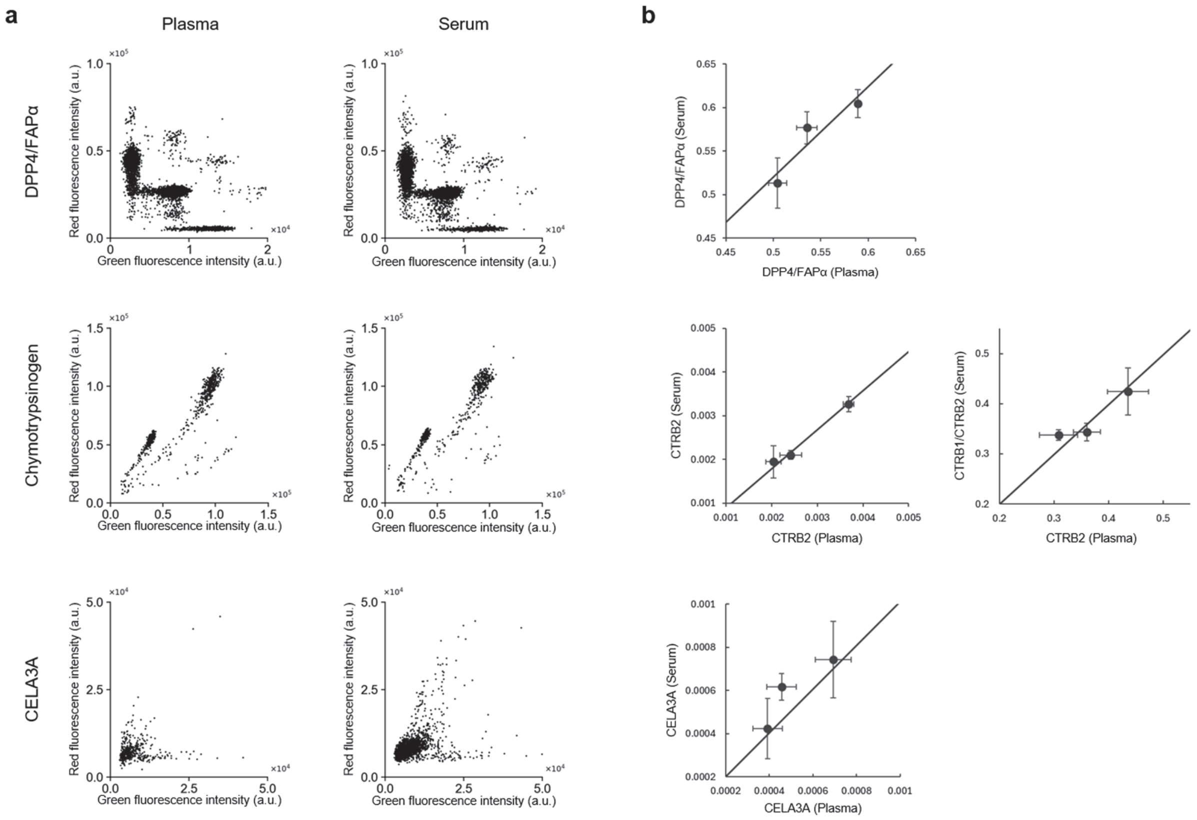


**Figure S26.** Comparison of plasma and serum measurements. (a) Representative scatter plots of green and red fluorescence intensities obtained from paired plasma and serum samples. DPP4/FAPα, chymotrypsinogen, and elastase activities were measured under the assay conditions described in **Table S4**. (b) Comparison of DPP4/FAPα, CTRB2, CTRB1/CTRB2, and CELA3A values between paired plasma and serum samples. Three biological replicates were analyzed, with each plasma and serum sample measured in technical triplicate. Data points represent the mean of technical triplicates, and error bars represent s.d. The solid line represents a linear fit to the three paired measurements constrained to pass through the origin.

a

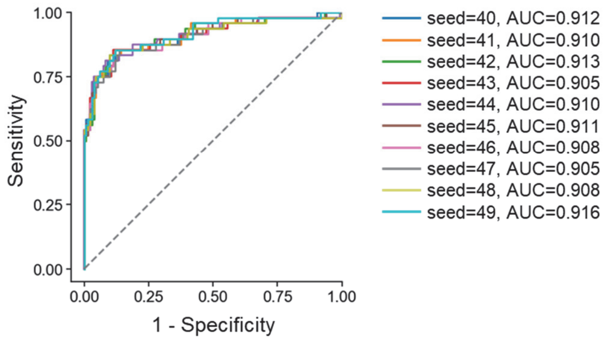


b

| | AUC | Sensitivity | Specificity |
|---|---|---|---|
| Median | 0.910 | 0.750 | 0.951 |
| Mean | 0.910 | 0.752 | 0.950 |
| s.d. | 0.003 | 0.012 | 0.009 |
| Min | 0.905 | 0.729 | 0.932 |
| Max | 0.916 | 0.771 | 0.962 |
| 25$^{th}$ percentile | 0.908 | 0.750 | 0.947 |
| 75$^{th}$ percentile | 0.912 | 0.750 | 0.955 |

**Figure S27.** Robustness of random forest prediction performance to random seed selection. (a) Receiver operating characteristic (ROC) curves in the validation cohort for random forest models trained with fixed training and validation cohorts while varying only the random seed (random_state = 40–49). (b) Summary statistics of AUROC, sensitivity, and specificity across seeds. For each seed, the operating cutoff was selected in the validation cohort to achieve a specificity of ≥93% among control subjects, using the same rule as in the main analysis. Sensitivity was then calculated at the corresponding seed-specific cutoff. s.d. indicates standard deviation across 10 random seeds.

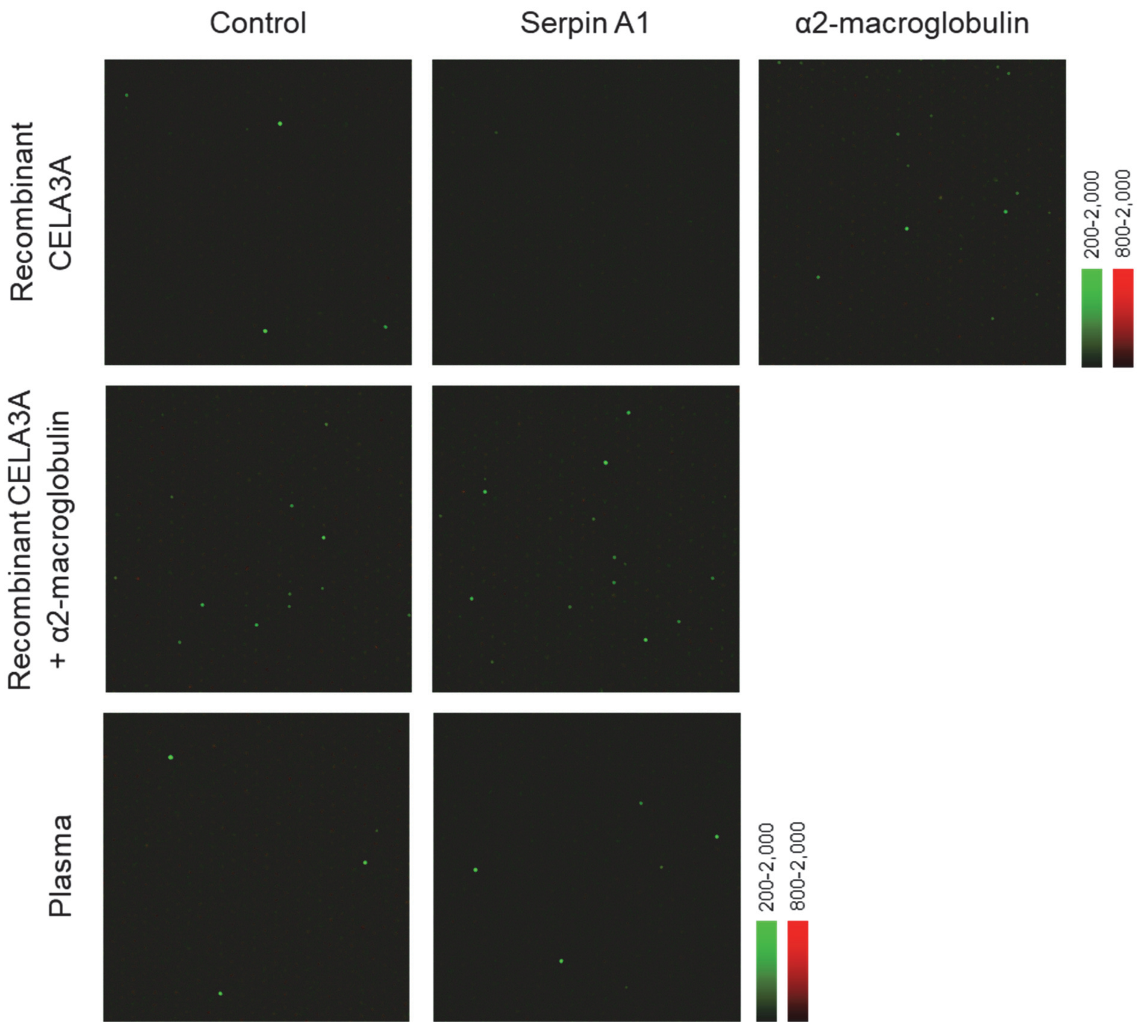


**Figure S28.** Effects of Serpin A1 and α2-macroglobulin. (Top row) Representative fluorescence images of the microdevice obtained after loading Suc-AAPAbu-PABC-sHMRG (10 μM), Suc-AAPV-PABC-dsSiR (15 μM), IR-dye 800 (30 μM), and recombinant CELA3A (50 ng/mL) pre-incubated for 1 h without inhibitor as a control (left), with Serpin A1 (5 μg/mL; center) or with α2-macroglobulin (5 μg/mL; right) into HEPES–Na buffer (125 mM, pH 7.4) containing $MgCl_2$ (1 mM), $CaCl_2$ (10 mM), NaCl (75 mM), DTT (100 μM), and Triton X-100 (150 μM), followed by incubation at 25 °C for 24 h. (Middle and bottom rows) Representative fluorescence images obtained after loading Suc-AAPAbu-PABC-sHMRG (10 μM), Suc-AAPV-PABC-dsSiR (15 μM), IR-dye 800 (30 μM), and recombinant CELA3A (50 ng/mL, pre-treated with α2-macroglobulin [5 μg/mL]) or plasma samples (1:100 diluted) pre-incubated for 1 h with or without Serpin A1 (5 μg/mL), in HEPES–Na buffer (125 mM, pH 7.4) containing $MgCl_2$ (1 mM), $CaCl_2$ (10 mM), NaCl (75 mM), DTT (100 μM), and Triton X-100 (150 μM), followed by incubation at 25 °C for 24 h.

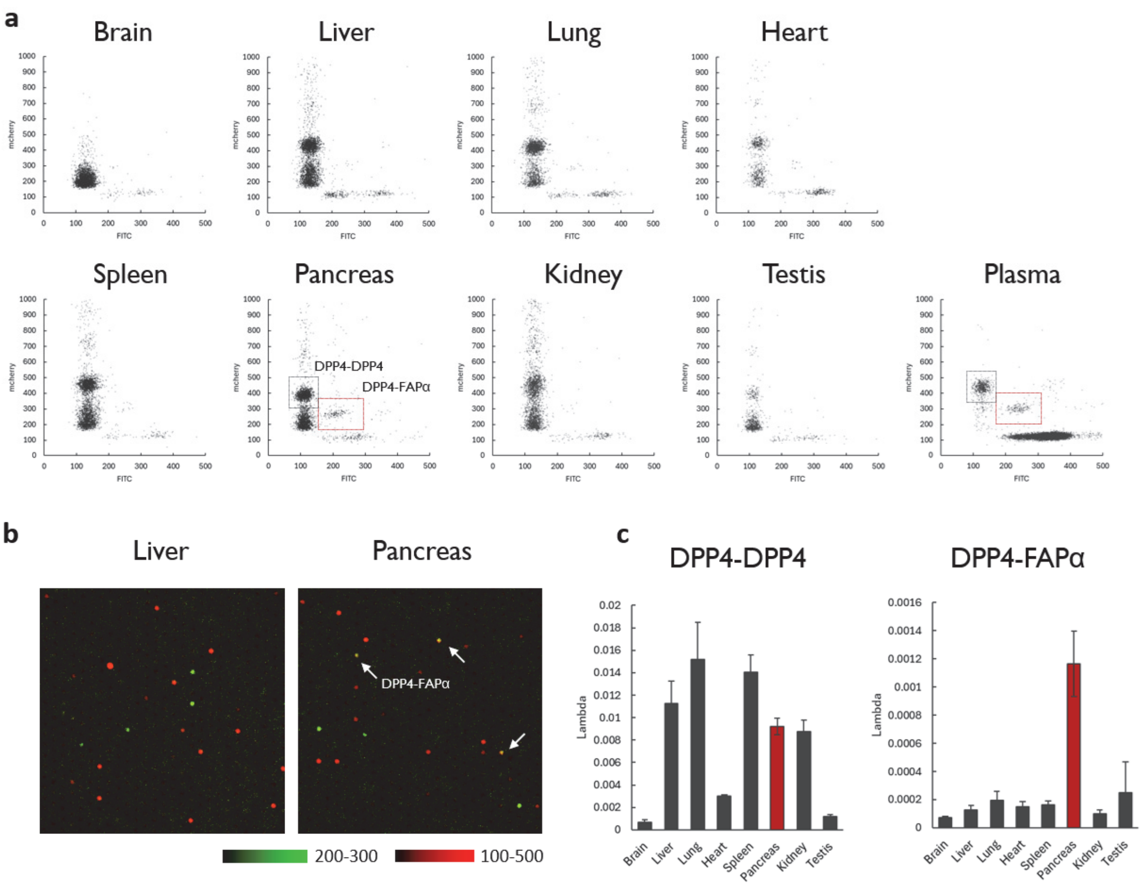


**Figure S29.** Analysis of tissue distribution of DPP4-FAPα heterodimer. (a) Mouse tissues (100 μg/mL) were mixed with Ac-GP-sHMRG (30 μM) and EP-sSiR (30 μM) in HEPES-Na buffer (pH 7.4) containing $CaCl_2$ (1 mM), $MgCl_2$ (1 mM), DTT (100 μM), BSA (1% w/v), and Triton X-100 (150 μM) and loaded into microdevice. Fluorescence images were acquired after incubating at 25°C for 2 h. (b) Fluorescence images of microdevice from (a) in the analysis of liver and pancreas. (c) Lambda values of clusters assigned as DPP4 homodimer and DPP4-FAPα heterodimer in different tissues. Error bars represent s.d. (n = 3).

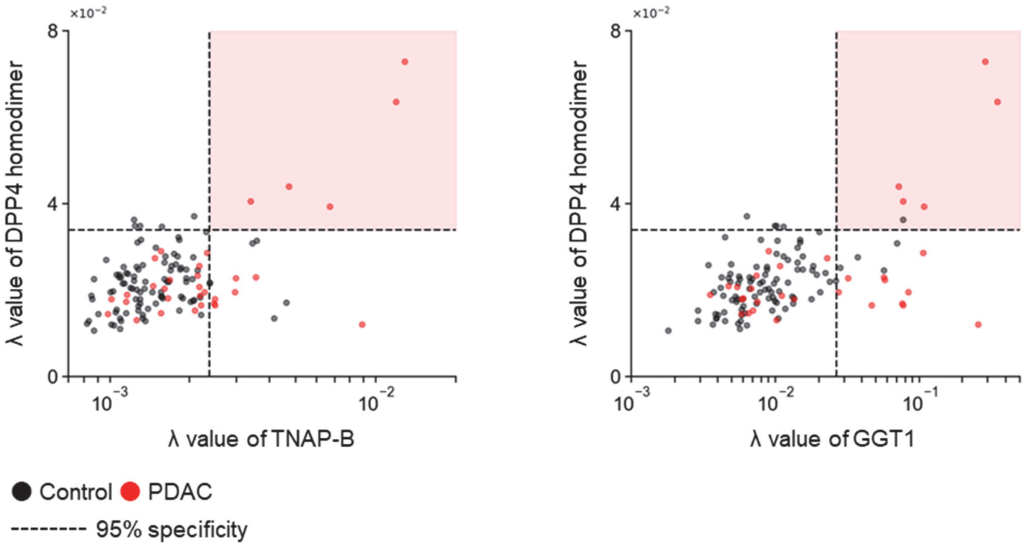


**Figure S30.** Association between DPP4 homodimer and biliary obstruction–associated markers. Correlation plots of DPP4 homodimer lambda values with TNAP-B and GGT1 lambda values in the screening cohort (PDAC, N = 32, red; control, N = 96, black). TNAP-B and GGT1 clusters are defined in **Figures S12** and **S13**, respectively. Dashed lines indicate the cutoffs corresponding to 95% specificity among control subjects.

**Supplementary data for synthesis and characterization of compounds**

**Chemical reagents and solvents**

Reagents and solvents were of the best grade available, supplied by Tokyo Chemical Industry, Wako Pure Chemical, Merck, and Watanabe Chemical Industries, and were used without further purification.

**Instruments for organic synthesis**

$^{1}H$ NMR spectra were recorded on Bruker AVANCE NEO 400 or JEOL JNM-LA400 spectrometer at 400 MHz and on JEOL JNM-ECZ500R spectrometer at 500 MHz. $^{13}C$ NMR spectra were recorded on Varian VNMRS 500 spectrometer at 125 MHz. High resolution mass spectra (HRMS) were obtained using JEOL JMS-T100LP AccuTOF LC-plus 4G (ESI). Preparative HPLC was performed on an Inertsil ODS-3 (10.0 × 250 mm) column (GL Sciences Inc.) using an HPLC system composed of a pump (PU-2080, JASCO) and a detector (MD-2015). Preparative MPLC was performed on an Isolera One purification system (Biotage) equipped with a Biotage SNAP Ultra C18 column (for reverse phase separation) or on an Isolera Prime purification system (Biotage) equipped with a silica gel column (silica gel 40 μm or Amino 40 μm, Yamazen) (for normal phase separation). LC-MS analysis was performed on an Acquity UPLC H-Class system (Waters) equipped with an Acquity UPLC BEH C18 1.7 μm (2.1 × 50 mm) column (Waters) and an MS detector (QDa or Xevo TQD, Waters).

**Synthesis of sHMRG-based and dsSiR-based peptidase probes**

sHMRG-based and dsSiR-based peptidase probes were synthesized following the scheme of synthesis based on capture and release (SCCR) previously reported[7] with minor modifications. For the conjugation step, 1 μmol of sHMRG-Cpz was used per reaction. For the capture and peptide synthesis steps, 1.5 μmol of sHMRG-Cpz or leuco dsSiR-Ert and 150 mg of azide resin were used per reactor. For large-scale preparation, the same procedure was performed in parallel across multiple reactors.

**Ac-RSPK-sHMRG**

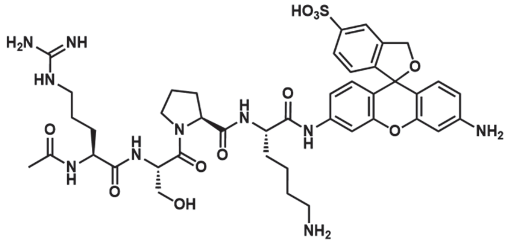


Prepared from sHMRG-Cpz[7] (2.0 μmol). Yield 65%.

LC chromatogram (A/B = 99:1 to 0:100 over 3.5 min; A = $H_2O$ + 0.1% TFA; B = $MeCN/H_2O$ (80:20, v/v) + 0.1% TFA; λ = 500 nm).

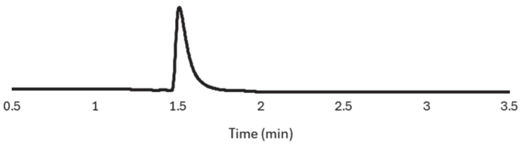


HRMS (ESI+): *m/z* Calcd. for [M+H]+ 907.3767, Found: 907.3741.

**Suc-RPY-sHMRG**

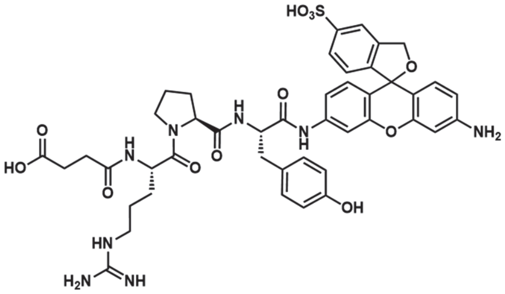

Prepared from sHMRG-Cpz (41 μmol). Yield 46%.

LC chromatogram (A/B = 99:1 to 0:100 over 3.5 min; A = $H_2O$ + 0.1% TFA; B = MeCN/$H_2O$ (80:20, v/v) + 0.1% TFA; λ = 500 nm).

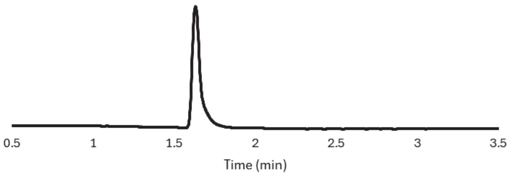


$^1$H NMR (400 MHz, DMSO-$d_6$ + 2 drops of 1M NaOD in $D_2O$): δ = 7.70-7.57 (m, 2H), 7.50 (d, *J* = 7.6 Hz, 1H), 7.12-7.03 (m, 1H), 6.89 (d, *J* = 7.9 Hz, 2H), 6.80 (d, *J* = 7.7 Hz, 1H), 6.69 (d, *J* = 6.0 Hz, 1H), 6.56 (d, *J* = 8.3 Hz, 1H), 6.51 (d, *J* = 7.9 Hz, 2H), 6.37 (s, 1H), 6.32 (d, *J* = 8.6 Hz, 1H), 5.19 (s, 2H), 4.49-4.37 (m, 2H), 4.34-4.24 (m, 1H), 3.62-3.51 (m, 1H), 3.49-3.38 (m, 1H), 3.01-2.77 (m, 4H), 2.41-2.29 (m, 1H), 2.29-2.17 (m, 1H), 2.17-2.06 (m, 2H), 2.06-1.95 (m, 1H), 1.90-1.70 (m, 3H), 1.56-1.36 (m, 4H).

HRMS (ESI-): *m/z* Calcd. for [M-H]- 911.3040, Found: 911.3019.

**Suc-AAVY-dsSiR**

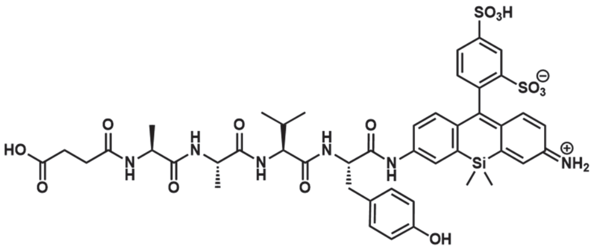

Prepared from leuco dsSiR-Ert[7] (32 μmol). Yield 28%.

LC chromatogram (A/B = 99:1 to 0:100 over 3.5 min; A = $H_2O$ + 0.1% TFA; B = MeCN/$H_2O$ (80:20, v/v) + 0.1% TFA; λ = 500 nm).

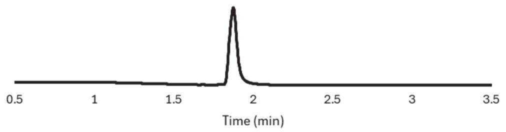


$^1$H NMR (400 MHz, DMSO-$d_6$): δ = 9.21 (brs, 1H), 8.19 (d, *J* = 1.4 Hz, 1H), 8.17-8.00 (m, 5H), 7.66 (dd, *J* = 7.9, 1.4 Hz, 1H), 7.57 (d, *J* = 8.5 Hz, 1H), 7.54-7.43 (m, 1H), 7.30 (d, *J* = 2.0 Hz, 1H), 7.16 (d, *J* = 9.5 Hz, 1H), 7.07-7.01 (m, 3H), 6.95 (d, *J* = 8.5 Hz, 1H), 6.73 (dd, *J* = 9.5, 1.6 Hz, 1H), 6.62 (d, *J* = 8.1 Hz, 2H), 4.57 (td, *J* = 7.2, 6.7 Hz, 1H), 4.31-4.20 (m, 2H), 4.12 (dd, *J* = 8.2, 7.1 Hz, 1H), 2.95 (dd, *J* = 13.8, 5.6 Hz, 1H), 2.79 (dd, *J* = 13.8, 8.8 Hz, 1H), 2.44-2.38 (m, 2H), 2.38-2.32 (m, 2H), 2.01-1.90 (m, 1H), 1.26-1.13 (m, 6H), 0.83-0.71 (m, 6H), 0.58-0.52 (m, 3H), 0.52-0.44 (m, 3H).

HRMS (ESI$^-$): *m/z* Calcd. for $[M-H]^-$ 991.2679, Found: 991.2632.

**Synthesis of Fmoc-Abu-PAB-OH**

Fmoc-Abu-OH (1.0 g, 3.1 mmol), DMT-MM (1.0 g, 3.7 mmol) and *p*-aminobenzyl alcohol (0.46 g, 3.7 mmol) were dissolved in MeCN (30 mL) and the mixture was stirred at 25°C for 14 h. The precipitate was filtered and the product was purified by column chromatography (silica, Hexane/AcOEt 100:0 to 0:100) to afford Fmoc-Abu-PAB-OH as a colorless solid (970 mg, 2.3 mmol, yield 73%).

Fmoc-Abu-PAB-OH

$^1$H NMR (400 MHz, DMSO-$d_6$): δ = 9.99 (s, 1H), 7.90 (d, *J* = 7.8 Hz, 2H), 7.78-7.72 (m, 2H), 7.65 (d, *J* = 8.2 Hz, 1H), 7.55 (d, *J* = 8.2 Hz, 2H), 7.42 (dd, *J* = 7.3, 7.3 Hz, 2H), 7.33 (ddd, *J* = 7.3, 7.3, 3.2 Hz, 2H), 7.24 (d, *J* = 8.2 Hz, 2H), 5.11 (t, *J* = 5.0 Hz, 1H), 4.43 (d, *J* = 5.0 Hz, 2H), 4.33-4.15 (m, 3H), 4.07 (td, *J* = 8.2, 5.5 Hz, 1H), 1.80-1.56 (m, 2H), 0.23 (t, *J* = 7.3 Hz, 3H).

$^{13}$C NMR (100 MHz, DMSO-$d_6$): δ = 170.9, 156.2, 143.9, 143.8, 140.7, 137.6, 137.4, 127.7, 127.1, 126.9, 125.4, 120.1, 118.9, 65.7, 62.6, 56.8, 46.7, 25.2, 10.6.

HRMS (ESI$^+$): *m/z* Calcd. for $[M+Na]^+$ 453.1785, Found: 453.1774.

**Fmoc-Arg(Boc)$_2$-PAB-OH**

Fmoc-Arg(Boc)$_2$-OH (1.0 g, 1.7 mmol), DMT-MM (0.55 g, 2.0 mmol) and *p*-aminobenzyl alcohol (0.25 g, 2.0 mmol) were dissolved in MeCN (30 mL) and the mixture was stirred at 25°C for 16 h. The precipitate was filtered and the product was purified by column chromatography (silica, Hexane/AcOEt 100:0 to 0:100) to afford Fmoc-Arg(Boc)$_2$-PAB-OH as a colorless solid (450 mg, 0.64 mmol, yield 38%).

$^1$H NMR (400 MHz, DMSO-$d_6$): δ = 11.5 (s, 1H), 10.0 (s, 1H), 8.33 (t, *J* = 5.5 Hz, 1H), 7.89 (d, *J* = 7.3 Hz, 2H), 7.78-7.70 (m, 3H), 7.54 (d, *J* = 8.7 Hz, 2H), 7.41 (dd, *J* = 7.8, 7.8 Hz, 2H), 7.32 (ddd, *J* = 7.3, 7.3, 3.2 Hz, 2H), 7.24 (d, *J* = 8.7 Hz,

2H), 5.11 (t, *J* = 5.9 Hz, 1H), 4.43 (d, *J* = 5.9 Hz, 2H), 4.30-4.10 (m, 4H), 3.33-3.25 (m, 2H), 1.75-1.50 (m, 4H), 1.47 (s, 9H), 1.38 (s, 9H).

$^{13}$C NMR (100 MHz, DMSO-$d_6$): δ = 170.9, 163.2, 156.1, 155.3, 152.1, 143.9, 143.8, 140.8, 137.5, 137.5, 127.7, 127.1, 126.9, 125.4, 120.2, 119.0, 82.9, 78.2, 65.7, 62.6, 55.1, 46.7, 29.3, 28.0, 27.7, 25.4.

HRMS (ESI$^+$): *m/z* Calcd. for [M+H]$^+$ 702.3497, Found: 702.3468.

**Fmoc-Val-PAB-OH**

Fmoc-Val-OH (1.0 g, 2.9 mmol), DMT-MM (0.98 g, 3.5 mmol) and *p*-aminobenzyl alcohol (0.44 g, 3.5 mmol) were dissolved in MeCN (30 mL) and the mixture was stirred at 25°C for 14 h. The precipitate was filtered and the product was purified by column chromatography (silica, Hexane/AcOEt 100:0 to 0:100) to afford Fmoc-Val-PAB-OH as a colorless solid (1.0 g, 2.3 mmol, yield 78%).

$^1$H NMR (400 MHz, DMSO-$d_6$): δ = 10.04 (s, 1H), 7.90 (d, *J* = 7.8 Hz, 2H), 7.79-7.72 (m, 2H), 7.65 (d, *J* = 9.2 Hz, 1H), 7.56 (d, *J* = 8.2 Hz, 2H), 7.41 (ddd, *J* = 7.3, 7.3, 3.2 Hz, 2H), 7.31 (ddd, *J* = 7.8, 7.3, 3.2 Hz, 2H), 7.25 (d, *J* = 8.2 Hz, 2H), 5.12 (brs, 1H), 4.43 (s, 2H), 4.33-4.18 (m, 3H), 3.98 (t, *J* = 8.3 Hz, 1H), 2.02 (m, 1H), 0.97-0.87 (m, 6H).

$^{13}$C NMR (100 MHz, DMSO-$d_6$): δ = 170.5, 156.3, 143.9, 143.8, 140.7, 137.6, 137.5, 127.7, 127.1, 127.0, 125.5, 120.2, 119.1, 65.8, 62.6, 61.1, 46.7, 30.4, 19.2, 18.7.

HRMS (ESI$^+$): *m/z* Calcd. for [M+Na]$^+$ 467.1941, Found: 467.1924.

**Synthesis of Suc-AAPAbu-PABC-sHMRG**

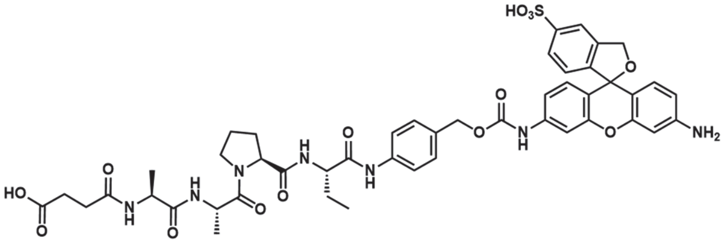


Procedures (2)-(3) were performed using an automated peptide synthesizer (Syro I, Biotage) equipped with a heating block. 30 μmol of sHMRG-Cpz was used, but the reactions were performed as parallel procedures of small scale reactions (1 μmol/tube and 1.5 μmol/reactor) as shown below.

(1) Conjugation: In 1.5 mL plastic tube, Fmoc-Abu-PAB-OH (4.3 mg, 10 μmol) was mixed with triphosgene (2.8 mg, 10 μmol) in THF (500 μL) and *N*,*N*-diisopropyl-*N*-ethylamine (DIEA, 2.8 μL, 15 μmol) was added dropwise. After mixing for 5 min, a solution of sHMRG-Cpz (1 μmol) in *N*-methylpyrrolidone (NMP, 10 μL) was added, and THF was removed *in vacuo* using a centrifugal evaporator over 2 h at 25°C.

(2) Capture and wash: In each reactor, Tentagel-azide beads[7] (150 mg) was suspended in Tris-HCl buffer (300 mM, pH 7.4; 300 μL). Reaction mixture of (1) corresponding to 1.5 μmol/reactor was diluted in *t*-BuOH (400 μL) and added into the reactor. $CuSO_4$ (10 mM in $H_2O$, 100 μL), tris[(1-benzyl-1*H*-1,2,3-triazol-4-yl)methyl]amine (TBTA, 30 mM in DMSO, 100 μL) and sodium ascorbate (30 mM in $H_2O$, 100 μL) were added to the reactor and stirred at 25°C for 4 h. The beads were washed six times with DMF.

(3) The peptide synthesis was performed by treating beads with conditions for (a) Fmoc deprotection and (b) amino acid coupling, successively until the full peptide sequence was prepared. (a) Piperidine (40% in DMF; 1200 μL) was added to beads and stirred at 25°C for 3 min. After removing the solvent, piperidine (40% in DMF; 600 μL) and DMF (600 μL) were added and stirred for 12 min. Beads were washed six times with DMF. (b) Fmoc-AA (0.4 M in DMF, 800 μL), HATU (0.4 M in DMF, 840 μL), DIEA (1.6 M in NMP, 400 μL) were added to beads and stirred at 30°C for 40 min. Beads were washed three times with DMF. Mono-*tert*-butyl succinate was used as a building block of Suc capping.

(4) Cleavage and purification: 90% TFA *aq.* (200 μL) was added to the beads and stirred at 25°C for 15 min. The solution was collected, and the beads were washed three times with acetone (200 μL). The combined solution from all reactors was collected, diluted with $H_2O$ and directly injected into a prep. MPLC column for purification ($C_{18}$; A/B = 95:5 isocratic 10-20 min, then the gradient was changed to 0:100 over 15 min; A = $H_2O$ + 0.1% TFA; B = MeCN + 0.1% TFA). The fractions containing the desired product were combined, concentrated *in vacuo*, and freeze-dried (10.8 μmol, yield 36%).

LC chromatogram (A/B = 99:1 to 0:100 over 3.5 min; A = $H_2O$ + 0.1% TFA; B = MeCN/$H_2O$ (80:20, v/v) + 0.1% TFA; λ = 500 nm).

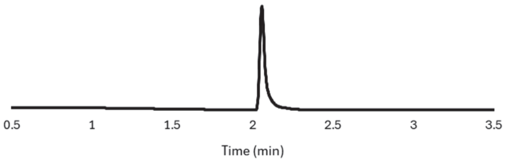


$^1$H NMR (400 MHz, DMSO-$d_6$ + 2 drops of 1M NaOD in $D_2O$): δ = 7.61 (s, 1H), 7.48 (d, *J* = 8.0 Hz, 1H), 7.28 (d, *J* = 1.8 Hz, 1H), 7.22 (d, *J* = 8.0 Hz, 2H), 7.00 (d, *J* = 8.0 Hz, 2H), 6.73 (dd, *J* = 1.8, 8.5 Hz, 1H), 6.67 (d, *J* = 8.0 Hz, 1H), 6.47 (d, *J* = 8.4 Hz, 1H), 6.39 (d, *J* = 8.5 Hz, 1H), 6.30 (d, *J* = 2.0 Hz, 1H), 6.23 (dd, *J* = 2.0, 8.4 Hz, 1H), 5.08 (s, 2H), 4.73 (s, 2H), 4.33-4.21 (m, 2H), 4.01-3.90 (m, 2H), 3.72-3.61 (m, 1H), 3.61-3.52 (m, 1H), 2.28-2.18 (m, 1H), 2.18-2.03 (m, 3H), 2.00-1.89 (m, 1H), 1.89-1.55 (m, 5H), 1.16-1.01 (m, 6H), 0.74 (t, *J* = 7.1 Hz, 3H).
HRMS (ESI$^-$): *m/z* Calcd. for [M-H]$^-$ 968.3142, Found: 968.3107.

**General scheme for preparation of dsSiR-based probes with PABC linkers**

Procedures (2)-(4) were performed using an automated peptide synthesizer (Syro I, Biotage) equipped with a heating block. The reactions were performed as parallel procedures of small scale reactions (1 μmol/tube and 1.5 μmol/reactor) as shown below.

(1) Conjugation: In 1.5 mL plastic tube, Fmoc-Val-PAB-OH or Fmoc-Arg(Boc)$_2$-PAB-OH (10 μmol) was mixed with triphosgene (2.8 mg, 10 μmol) in THF (500 μL), and DIEA (2.8 μL, 15 μmol) was added dropwise. After 5 min, a solution of leuco dsSiR-Ert (1 μmol) in NMP (10 μL) was added and the resultant mixture was placed *in vacuo* in a centrifugal evaporator for 2 h.

(2) Capture and wash: Tentagel-azide beads (100 mg) were suspended in Tris-HCl buffer (300 mM, pH 7.4;

300 μL) in the reactor. Reaction mixture of (1) was diluted in *t*-BuOH (400 μL) and added into the reactor. $CuSO_4$ (10 mM in $H_2O$, 100 μL), TBTA (30 mM in DMSO, 100 μL) and sodium ascorbate (30 mM in $H_2O$, 100 μL) were added to the reactor and stirred at 25°C for 4 h. The beads were washed six times with DMF.

(3) The peptide synthesis was performed by treating beads with conditions for (a) Fmoc deprotection and (b) amino acid coupling, successively until the full peptide sequence was prepared. (a) Piperidine (40% in DMF; 1200 μL) was added to beads and stirred at 25°C for 3 min. After removing the solvent, piperidine (40% in DMF; 600 μL) and DMF (600 μL) were added and stirred for 12 min. Beads were washed six times with DMF. (b) Fmoc-AA (0.4 M in DMF, 800 μL), HATU (0.4 M in DMF, 840 μL), DIEA (1.6 M in NMP, 400 μL) were added to beads and stirred at 30°C for 40 min. Beads were washed three times with DMF. Mono-*tert*-butyl succinate was used as a building block of Suc capping.

(4) Oxidation: 2,3-Dichloro-5,6-dicyano-1,4-benzoquinone (DDQ) (10 mg, 0.044 mmol) dissolved in acetone (1 mL) was added to the beads and stirred at 25°C for 30 min. Beads were washed six times with acetone.

(5) Cleavage: 90% TFA *aq.* (200 μL) was added to the beads and stirred at 25°C for 15 min. The solution was collected, and the beads were washed three times with acetone (200 μL). The combined solution from all reactors were pooled, and directly injected into a prep. MPLC column for purification ($C_{18}$; A/B = 95:5 isocratic 10-20 min, then the gradient was changed to 0:100 over 15 min; A: $H_2O$ + 0.1% TFA; B: MeCN + 0.1% TFA). The product was further purified over prep. HPLC twice (1st separation: $C_{18}$; A/B = 100:0 to 0:100 over 30 min; A = $H_2O$ + 0.1% TFA; B = MeCN + 0.1% TFA. 2nd separation: $C_{18}$; A/B = 100:0 to 0:100 over 30 min; A = 10 mM ammonium formate; B = MeCN + 10 mM ammonium formate). The fractions containing the desired product were collected and freeze-dried.

**Suc-AAPV-PABC-dsSiR**

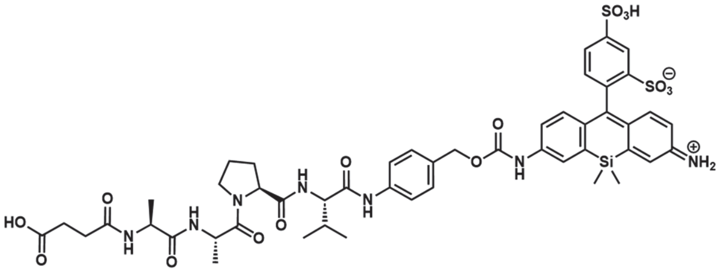


Prepared from leuco dsSiR-Ert (63 μmol). Yield 7%.

LC chromatogram (A/B = 99:1 to 0:100 over 3.5 min; A = $H_2O$ + 0.1% TFA; B = MeCN/$H_2O$ (80:20, v/v) + 0.1% TFA; λ = 500 nm).

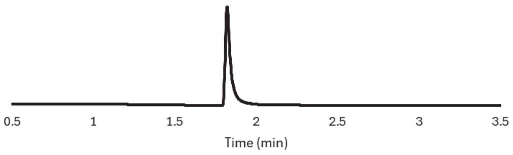

$^1$H NMR (400 MHz, DMSO-$d_6$ + 2 drops of 10 mM NaOD in $D_2O$): δ = 8.19 (d, *J* = 1.4 Hz, 1H), 8.01 (d, *J* = 2.3 Hz, 1H), 7.71 (dd, *J* = 1.8, 7.8 Hz, 1H), 7.62 (d, *J* = 8.2 Hz, 2H), 7.41-7.35 (m, 3H), 7.31 (d, *J* = 2.3 Hz, 1H), 7.15 (d, *J* = 9.6 Hz, 1H), 7.10 (d, *J* = 7.8 Hz, 1H), 6.92 (d, *J* = 8.7 Hz, 1H), 6.72 (dd, *J* = 2.3, 9.6 Hz, 1H), 5.14 (s, 2H), 4.51-4.40 (m, 2H), 4.30-4.19 (m, 2H), 3.56-3.48 (m,2H), 2.45-2.38 (m, 2H), 2.37-2.30 (m, 2H), 2.08-1.96 (m, 2H), 1.94-1.76 (m, 3H), 1.24-1.11 (m, 6H), 0.90 (d, *J* = 6.8 Hz, 6H), 0.53 (s, 3H), 0.46 (s, 3H).

HRMS (ESI$^-$): *m/z* Calcd. for $[M-H]^-$ 1074.3051, Found: 1074.3068.

**Ac-IGGR-PABC-dsSiR**

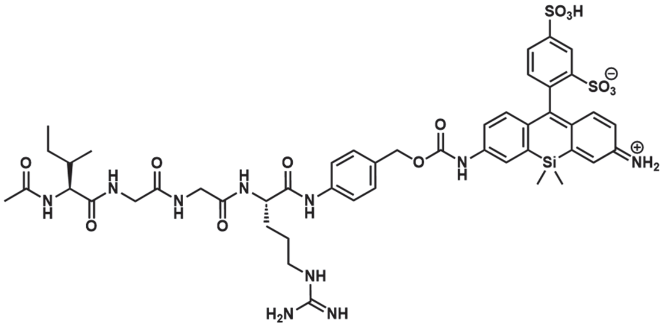

Prepared from leuco dsSiR-Ert (16 μmol). Yield 8%.

LC chromatogram (A/B = 99:1 to 0:100 over 3.5 min; A = $H_2O$ + 0.1% TFA; B = $MeCN/H_2O$ (80:20, v/v) + 0.1% TFA; λ = 500 nm).

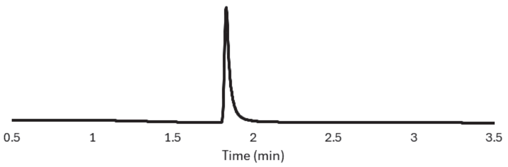


HRMS (ESI$^-$): *m/z* Calcd. for $[M-H]^-$ 1061.3323, Found: 1061.3296.

**Synthesis of γGlu-sSiR**

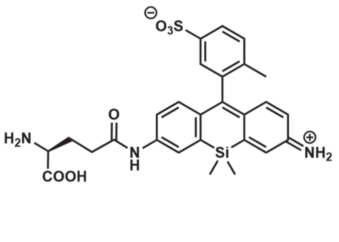

Leuco-sSiR was prepared according to the literature[2]. Leuco-sSiR (2.0 mg, 4.7 μmol) was mixed with Boc-Glu-O*t*Bu (1.0 mg, 3.3 μmol), ethyl 2-cyano-2-((dimethyliminio)(morpholino)methyloxyimino)acetate hexafluorophosphate (COMU, 1.4 mg, 3.3 μmol) and DIEA (5.0 μL, 28 μmol) in DMF (100 μL) and the reaction

mixture was stirred at 25°C for 12 h. DMF (200 μL) and chloranil (2.3 mg, 93 μmol) were added, and the reaction mixture was stirred at 25°C for 4 h. TFA/$H_2O$ (9:1, 1.0 mL) was added, and the reaction mixture was stirred at 25°C for 2 h. The mixture was diluted with $H_2O$ and the product was purified over MPLC ($C_{18}$; A/B = 100:0 to 0:100; A = $H_2O$ + 0.1% TFA; B = MeCN + 0.1% TFA). The fractions containing the desired product were flash frozen and lyophilized. γGlu-sSiR was acquired as a purple solid (2.4 μmol, yield 51%).
LC chromatogram (A/B = 99:1 to 0:100 over 3.5 min; A = $H_2O$ + 0.1% TFA; B = MeCN/$H_2O$ (80:20, v/v) + 0.1% TFA; λ = 500 nm).

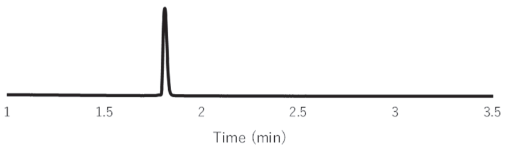


HRMS (ESI$^-$): *m/z* Calcd. for $[M-H]^-$ 550.1474, Found: 550.1459.

#### Synthesis of sTG-PHBA-P(Et)-GR

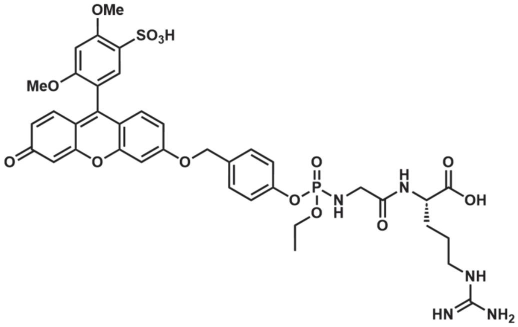


Br-PHBA-P(Et)-GR(Pbf)-O*t*Bu was synthesized following the reported procedure[21]. sTG (3.0 mg, 7.0 μmol) was dissolved in NMP (15 μL) and DIPEA (15 μL), and dry THF (25 μL) was added. Br-PHBA-P(Et)-GR(Pbf)-O*t*Bu (15 mg, 18 μmol) was added to the mixture. The resultant mixture was stirred at 25°C for 16 h. TFA/$H_2O$ (9:1, 2.0 mL) was added, and the reaction mixture was stirred at 25°C for 4 h. The mixture was diluted with $H_2O$ and the product was purified over MPLC (C18; A/B = 100:0 to 0:100; A = $H_2O$ + 0.1% TFA, B = MeCN + 0.1% TFA,) and prep. HPLC (C18; A/B = 100:0 to 0:100, 30 min: A = 5 mM triethylammonium acetate aq., B = MeCN). The fractions containing the desired product were collected and freeze-dried (0.30 μmol, yield 4%).
LC chromatogram (A/B = 99:1 to 0:100 over 3.5 min; A = $H_2O$ + 0.1% TFA; B = MeCN/$H_2O$ (80:20, v/v) + 0.1% TFA; λ = 500 nm).

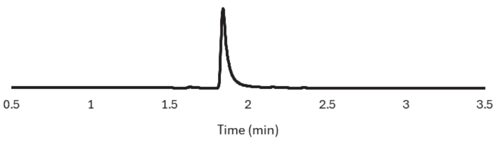

HRMS ($ESI^-$): $m/z$ Calcd. for $[M-H]^-$ 854.2114, Found: 854.2081.

**Synthesis of Res-PHBA-P(Et)-AK**

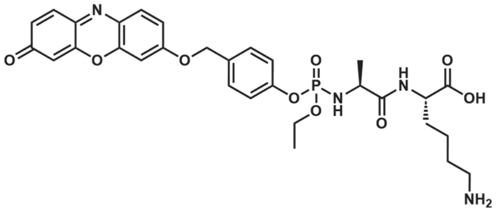


Br-PHBA-P(Et)-AK(Boc)-O*t*Bu was synthesized following the reported procedure[21]. Resorufin (20 mg, 94 µmol) was dissolved in NMP (150 µL) and DIPEA (150 µL), and dry THF (250 µL) was added. Br-PHBA-P(Et)-AK(Boc)-O*t*Bu (85 mg, 0.13 mmol) was added to the mixture. The resultant mixture was stirred at 25°C for 48 h. Column chromatography (silica; A/B = 100:0 to 0:100 over 15 min; A = hexane; B = AcOEt) was performed and the fractions containing the desired product were collected and evaporated. TFA/$H_2O$ (9:1, 2.0 mL) was added, and the reaction mixture was stirred at 25°C for 4 h. The mixture was diluted with $H_2O$ and the product was purified over MPLC (C18; A/B = 100:0 to 0:100; A = $H_2O$ + 0.1% TFA; B = MeCN + 0.1% TFA) and prep. HPLC (C18; A/B = 100:0 to 0:100, 30 min: A = 5 mM triethylammonium acetate aq., B = MeCN). The fractions containing the desired product were collected and freeze-dried (1.9 µmol, yield 2%).

LC chromatogram (A/B = 99:1 to 0:100 over 3.5 min; A = 10 mM Ammonium formate aq.; B = MeCN/ 10 mM Ammonium formate aq. (80:20, v/v) ; λ = 450 nm).

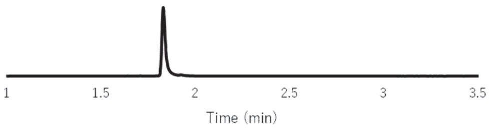


HRMS ($ESI^+$): $m/z$ Calcd. for $[M+H]^+$ 627.2214, Found: 627.2211.

**Synthesis of sTG-mADP**

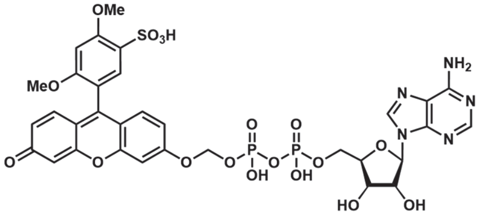


sTG-mPhos was prepared according to the literature[9]. To a solution of sTG-mPhos (15 mg, 0.028 mmol), AMP-Imidazole (22 mg, 0.056 mmol) and triethylamine (3.9 µL, 0.028 mmol) in dry DMF (1.2 mL) was added $ZnCl_2$ (76 mg, 0.56 mmol, 20 eq.) under $N_2$. The reaction mixture was stirred at 25°C for 72 h. The product was purified by preparative HPLC ($C_{18}$; A/B = 99:1 to 0:100 over 25 min; A = $H_2O$ + 0.1% TFA; B = MeCN + 0.1% TFA) to obtain sTG-mADP as a yellow solid (22 mg, 26 µmol, yield 91%).

$^1$H NMR (500 MHz, DMSO-$d_6$): δ 8.65 (d, *J* = 6.5 Hz, 1H), 8.43 (d, *J* = 6.5 Hz, 1H), 7.67–7.61 (m, 2H), 7.54–7.46 (m, 2H), 7.29–7.24 (m, 1H), 7.06–7.00 (s, 1H), 6.97–6.90 (m, 2H), 5.94 (dd, *J* = 5.7, 6.1 Hz, 1H), 5.83 (d, *J* = 11.8 Hz, 2H), 4.56–4.50 (m, 1H), 4.25–4.12 (m, 4H), 3.96 (s, 3H), 3.77 (s, 3H).
$^{13}$C NMR (125 MHz, DMSO-$d_6$): δ 176.4, 164.3, 160.0, 159.4, 158.5, 158.4, 158.2, 157.9, 156.4, 150.2, 145.4, 141.9, 133.1, 131.7, 131.1, 128.5, 118.5, 118.0, 117.4, 116.8, 116.5, 114.2, 109.7, 103.0, 97.1, 88.1, 87.8, 74.1, 70.0 (*J* = 9.96 Hz), 66.0, 56.3, 56.0.
HRMS (ESI$^+$): *m/z* Calcd. for [M+H]$^+$ 868.0938, Found: 868.0959.

**Synthesis of sTG-diPrAMP**

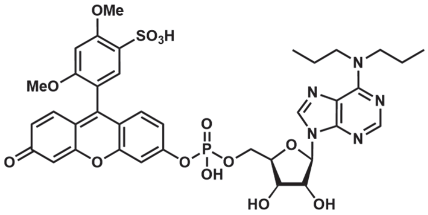


sTG was prepared according to the literature[9]. To a solution of sTG (15 mg, 0.033 mmol, 1.0 eq.), diPrAMP (62 mg, 0.14 mmol, 4.3 eq.), DMAP (4.0 mg) and $NEt_3$ (5 drops) in *t*-BuOH/$H_2O$ (3 mL/1 mL) was added 2 N NaOH aq. (100 μL) and EDCI (13 mg, 0.067 mmol, 2.0 eq.) at R.T. The reaction mixture was refluxed for 15 min, then repeatedly added EDCI (13 mg, 0.067 mmol, 2.0 eq.) eight times every 15 min during stirring. The reaction mixture was further refluxed for 1 h, then evaporated. The residue was purified by preparative HPLC (A:B = 99:1 to 99:1 (3 min) to 0:100 (23 min); A = 50 mM TEAA buffer; B = MeCN). The combined solution was evaporated and the concentrated solution was further purified by preparative HPLC (A:B = 90:10 to 90:10 (5 min) to 0:100 (25 min); A = $H_2O$ + 0.1% TFA; B = MeCN + 0.1% TFA) to obtain sTG-diPrAMP as an orange solid (8.3 mg, 9.6 μmol, yield 29%).
$^1$H NMR (500 MHz, DMSO-$d_6$): δ 8.46 (s, 1H), 8.28 (s, 1H), 7.62 (s, 1H), 7.56 (d, *J* = 9.9 Hz, 1H), 7.41-7.33 (m, 2H), 7.32–7.23 (m, 1H), 6.92–6.85 (m, 2H), 6.72–6.63 (m, 1H), 5.99–5.95 (m, 1H), 4.61–4.55 (m, 1H), 4.26–4.10 (m, 4H), 3.95 (s, 3H), 3.76 (d, J = 7.8 Hz, 3H), 1.68–1.56 (m, 4H), 0.94–0.82 (m, 6H).
$^{13}$C NMR (125 MHz, DMSO-$d_6$): δ 184.0, 158.8, 158.5, 158.2, 153.6, 153.6, 153.0, 152.0, 150.6, 150.6, 147.7, 147.4, 138.1, 131.0, 130.4, 129.3, 129.0, 128.7, 118.9, 118.8, 118.1, 114.8, 110.3, 104.5, 96.8, 95.8, 86.6, 74.0, 71.1, 56.0, 55.8, 45.7, 11.1, 8.7. HRMS (ESI$^+$): *m/z* Calcd. for [M+H]$^+$ 842.2108, Found: 842.2138.

**Preparation of fluorogenic probes for other biomarker candidate enzymes**

sTG-PHBA-P(Et)-AY and Res-PHBA-P(Et)-GL were prepared as reported[21]. Ac-GP-sHMRG, A-sHMRG, EP-sSiR, and Y-sSiR were prepared as reported[2]. γGlu-sHMRG was prepared as reported[7]. sTG-mPhos, sTM-Phos, and sTM-dCMP were prepared as reported[9].

The fluorogenic probe for amylase was synthesized according to previously established principles for glycosidase probes. The specific amylase-recognition moiety was conjugated to the fluorophore following a standardized protocol. Detailed synthetic procedures and full structural characterization of this probe will be reported elsewhere. The reactivity and specificity of the probe toward its target enzyme were validated as shown in Supplementary **Fig. S18**.

In all cases, LC-MS analysis was performed to confirm that the chemical purity of the compound (450 nm for sTG and resorufin-based probes, 500 nm for sHMRG, sSiR and sTM-based probes) was higher than 95%.

**NMR charts of fluorogenic probes used to measure final candidate biomarkers.**

**Suc-RPY-sHMRG**

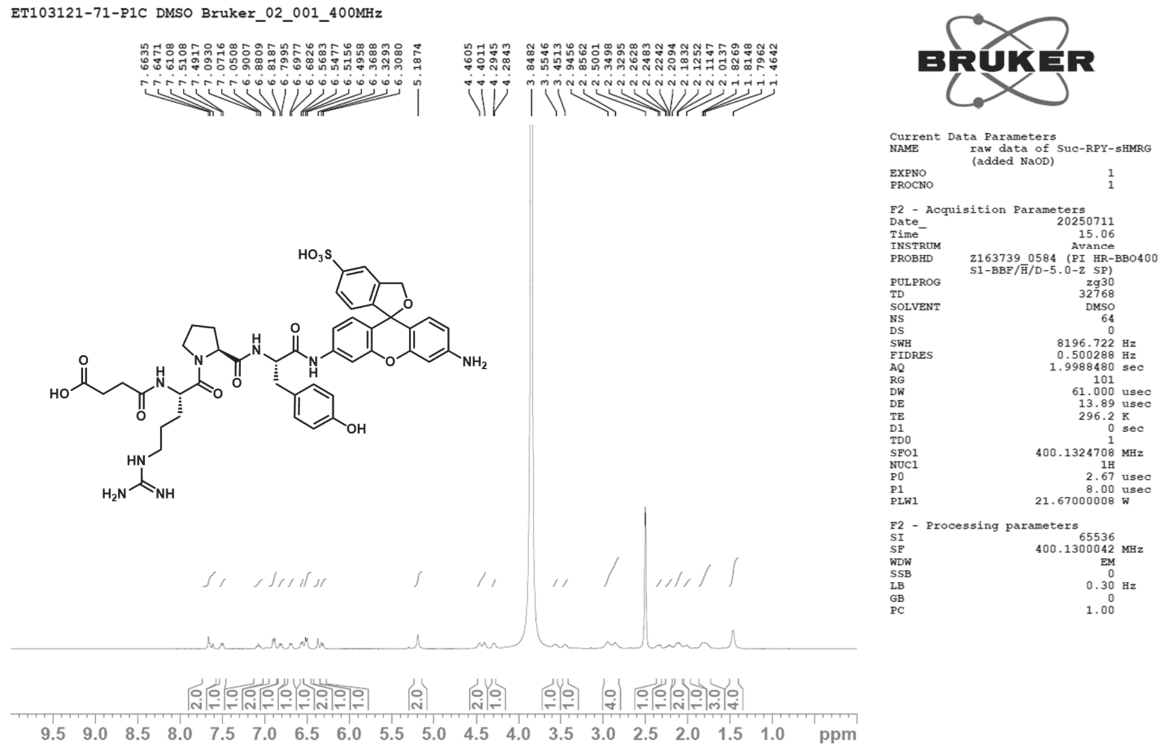


**Suc-AAVY-dsSiR**

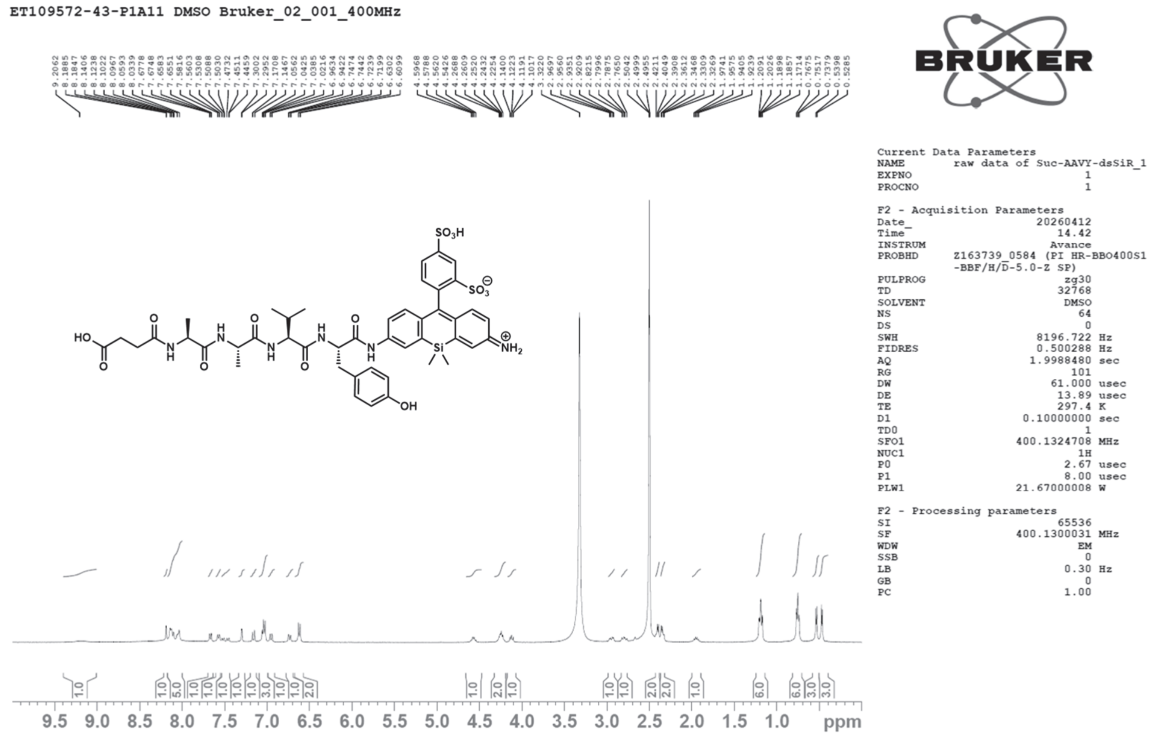

**Suc-AAPAbu-PABC-sHMRG**

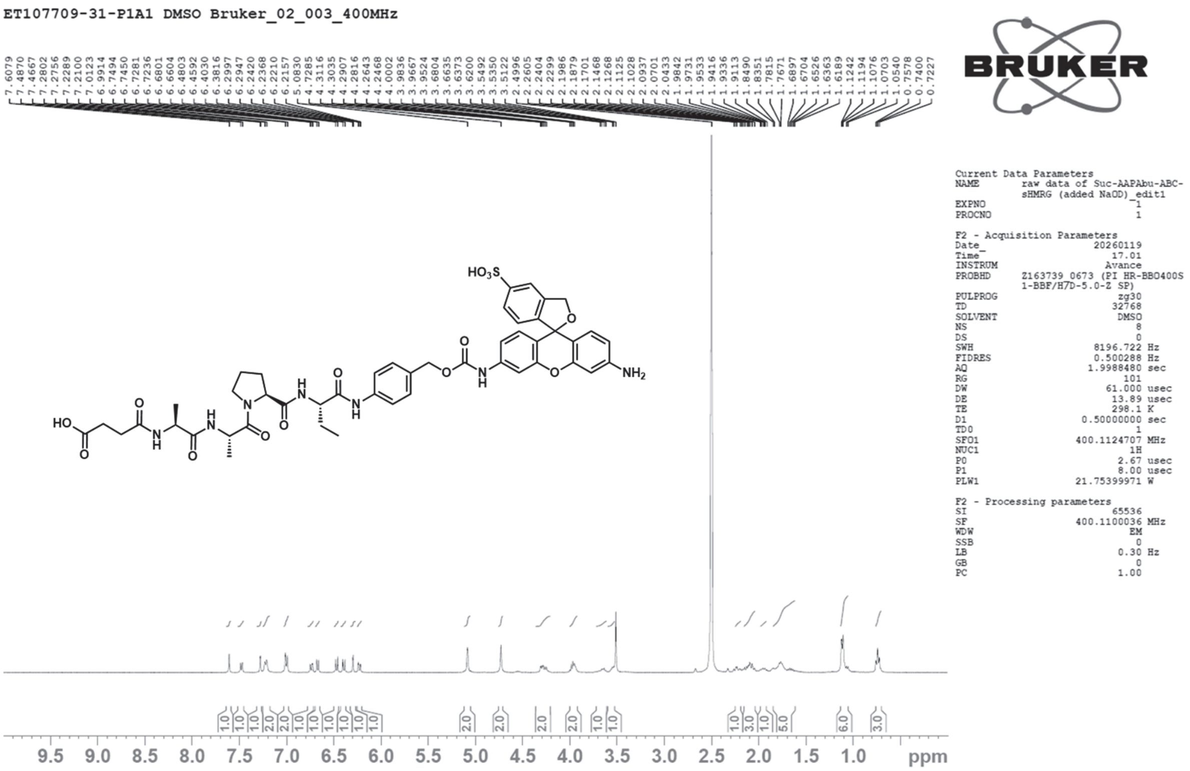


**Suc-AAPV-PABC-dsSiR**

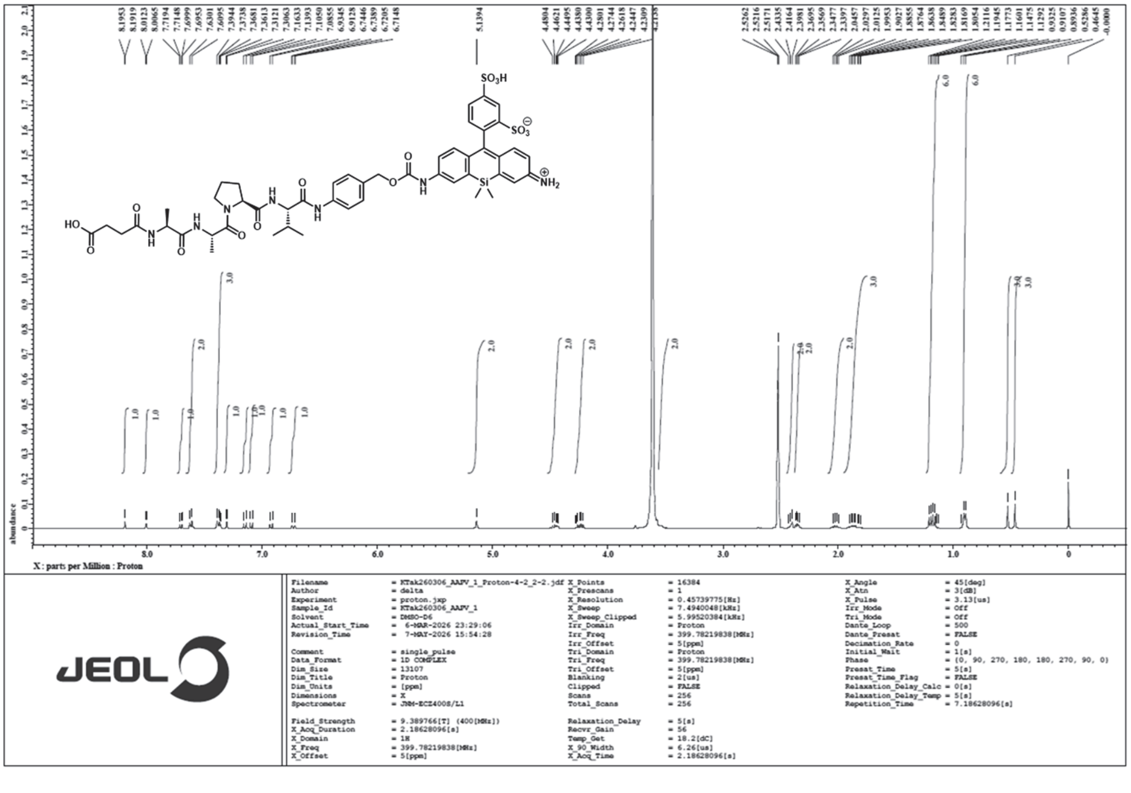
